# Recombinant SIRT1 expression and SIRT1–METTL3 interaction analysis with computational identification of phytochemical SIRT1 activators and inhibitors

**DOI:** 10.64898/2026.10.18.759935

**Authors:** Md Kazy Ebnul Hasan, Md. Nahid Hasan Joy, Liqing Lin, Md. Sanowar Hossan, Shahad Shahriar, Md. Mahedi Hasan Sourov, Md. Naymur Rahman, Md. Faruk Hasan, Md. Enamul Haque, Abdullah-Al Jubayer, Mamudul Hasan Razu, Md. Samiul Islam, Mengqi Lv, Yunyu Shi

**Author notes:** Corresponding Author: Md Kazy Ebnul Hasan.

## Abstract

Sirtuin 1 (SIRT1), a NAD-dependent histone deacetylase, regulates diverse biological processes including aging, stress response, and cancer progression. Owing to its dual roles in tumor suppression and promotion, selective modulators of SIRT1 have attracted considerable therapeutic interest. This study explored the interaction of SIRT1 with METTL3 and screened phytochemicals from Indian medicinal plants for potential SIRT1 activation or inhibition. Human SIRT1 (residues 183–665) was cloned, expressed in E. coli, purified, and validated by isothermal titration calorimetry (ITC) to confirm binding with METTL3. Computational approaches, including molecular docking, ADMET profiling, molecular dynamics (MD) simulations, MM-PBSA free energy estimation, and frontier molecular orbital (FMO) analysis, were employed to characterize ligand interactions. Docking revealed Catechin, Epicatechin, Kaempferol, and Bisdemethoxycurcumin as potent inhibitors, showing stronger binding than the reference inhibitor Selisistat by stabilizing the catalytic pocket via key hydrogen bonds and π–π stacking interactions. Among activators, Ginsenosides and Carnosol demonstrated higher affinities than resveratrol, binding the allosteric site through hydrophobic and electrostatic interactions. MD simulations highlighted Catechin and Ginsenosides as the most stable modulators, supported by reduced RMSD, compact Rg values, and favorable hydrogen bond occupancy. MM-PBSA confirmed their strong binding free energies, while ADMET analysis predicted favorable pharmacokinetic properties and low toxicity, particularly for Catechin and Ginsenosides.

Overall, this study integrates expression and purification of the protein and computational modeling to identify phytochemicals as potential SIRT1 modulators and provides novel insights into the SIRT1–METTL3 regulatory axis. These findings highlight Catechin and Ginsenosides as promising scaffolds for therapeutic strategies targeting cancer and metabolic disorders.

## 1. Introduction

The sirtuin family has become recognized as key regulators of age-related diseases, neurodegeneration, obesity, inflammation, heart disease, and cancer. In mammals, there are seven sirtuin members (SIRT1–SIRT7), with SIRT1 being the most extensively studied over the last decade [1]. SIRT1, a NAD+-dependent histone deacetylase, has been shown to have a regulatory function in energy homeostasis, inflammation, oxidative stress, mitochondrial biogenesis, cell apoptosis, and autophagy [2,3]. Structurally, SIRT1 contains three main regions: the N-terminal region (residues 183–229), the catalytic core region (residues 229–516), and a C-terminal regulatory segment (CTR) located between residues 641–665 [4]. The N-terminal region (residues 183–229) is crucial for binding sirtuin-activating compounds (STACs), while the catalytic domain, where substrate binding occurs, is composed of a larger NAD+ binding Rossmann-fold domain and a smaller zinc-binding domain [5].

Therefore, SIRT1 is considered a most promising therapeutic target for a broad range of diseases, encompassing cancer, as well as cardiovascular, respiratory, and other pathological conditions.

SIRT1 exerts dual, context-dependent effects on cell proliferation by regulating the expression and acetylation mechanisms of key proteins [6]. Its impact varies significantly depending on the cell type and the specific downstream molecules involved. For example, SIRT1 facilitates the proliferation of endometrial cancer cells by decreasing the acetylation levels of the autophagy-associated protein LC [6]. It also drives proliferation in oligodendrocyte progenitor cells by deacetylating the retinoblastoma (Rb) protein within the Rb/E2F1 complex, leading to E2F1 release [7]. Furthermore, SIRT1 directly regulates transcription factors like E2F1 (stimulating macrophage proliferation) and p53 (promoting hepatocellular carcinoma cell proliferation), where p53 is a key target in cancer therapy [8]. When Sirt1 becomes activated, it can inhibit apoptosis by deacetylating tumor-suppressor proteins [9]. As a result, certain Sirt1 inhibitors that prevent p53 deacetylation demonstrate anticancer potential [10,11]. Activation of Sirt1 also reduces the tumor-suppressive function of p53 at the transcriptional level by deacetylating the proto-oncogene BCL6 [12]. In addition, Sirt1 can deacetylate members of the FOXO family, thereby suppressing FOXO-dependent transcription and apoptosis pathways. SIRT1 also regulates DNA repair pathways: it deacetylates Ku70 and influences Bax-mediated apoptosis [13]. Ku70 binds to Bax, preventing its translocation to mitochondria, thereby inhibiting apoptosis and extending cell survival [13]. Similarly, Sirt1-induced deacetylation of KAP1 may contribute to its anti-apoptotic effects, as KAP1 plays a role in non-homologous end joining (NHEJ), a primary pathway for repairing DNA double-strand breaks [14].

Given the diverse roles of SIRTs in numerous biological processes, researchers have developed both inhibitors and activators as tools to investigate their functions and as potential therapeutic agents for various diseases and conditions. In general, activators are considered to have greater therapeutic promise than inhibitors, possibly due to their higher target selectivity within the enzyme family and reduced likelihood of adverse effects [15]. Nonetheless, the availability of activators remains relatively limited compared to inhibitors. SIRT1 activators are generally divided into two groups: natural polyphenols and structurally unrelated synthetic compounds. Among these, resveratrol, a polyphenolic compound abundantly found in grapes and red wine, was identified in 2003 as the first activator of SIRT1 [16]. Resveratrol functions as an allosteric activator of SIRT1, capable of enhancing its activity by 50% (EC1.5) at a concentration of 46.2 μM. It has been demonstrated to extend the lifespan of diverse organisms, ranging from yeast to mammals [16].

The role of sirtuins in the progression of various diseases has drawn significant attention, and inhibiting these deacetylases is being explored as a potential therapeutic approach. Many of the known inhibitors target Sirt1/2 isoforms. Splitomicin, an inhibitor, acts on human SIRT1 and SIRT2 with IC values of 96 μM and 113 μM, respectively [17]. Treatment with splitomicin decreases deacetylase activity, increases tissue factor mRNA expression in stimulated endothelial cells, and promotes NF-κB/p65 nuclear translocation. Moreover, nicotinamide blocks SIRT1 activity, thereby promoting p53-dependent apoptosis in cancer and neurodegeneration models [18].

However, the relationship between SIRT1 and METTL3 plays a pivotal role in regulating cancer metastasis through post-translational modifications. SIRT1, a class III histone deacetylase, directly deacetylates METTL3 at lysine 177 (K177), a methyltransferase involved in RNA modification [19]. This deacetylation is crucial for METTL3’s nuclear localization [20], where it promotes m6A methylation on specific mRNAs, thereby influencing cancer cell survival, proliferation, and metastasis by enhancing metastatic pathways [19]. Once in the nucleus, METTL3 acts as the catalytic core of the m6A methyltransferase complex, facilitating m6A modification on specific mRNA transcripts. This modification enhances mRNA stability, splicing, translation, and gene expression associated with tumorigenesis and metastasis, with elevated m6A levels being linked to increased cancer cell invasion and metastasis. Inhibiting SIRT1 increases METTL3 acetylation by blocking its deacetylation at the K177 residue, which impairs METTL3’s nuclear translocation and causes it to accumulate in the cytoplasm. This retention in the cytoplasm reduces METTL3’s ability to modify m6A in the nucleus, directly affecting its oncogenic functions, particularly those involved in metastasis, as it can no longer enhance the translation of pro-metastatic mRNAs like IL-6, NRP2, and CAV1. By inhibiting SIRT1, nuclear accumulation of METTL3 is curtailed, reducing its m6A methylation activity and lowering the expression of mRNAs that promote metastasis, ultimately suppressing cancer cell invasion and migration [21]. Furthermore, combining SIRT1 inhibition with treatments like aspirin, which directly acetylates METTL3, enhances acetylation and blocks METTL3’s oncogenic potential, providing a multi-faceted approach to reducing metastasis.

Overall, SIRT1 has dual roles in tumor suppression and progression, highlighting the importance of selective modulators with improved safety and specificity. Besides, the SIRT1–METTL3-mediated m6A methylation axis represents a promising but underexplored avenue for such intervention. Bioactive natural phytochemicals, with their structural diversity and low toxicity, offer a promising source of novel SIRT1 activators and inhibitors. In this study, we combined experimental and computational approaches to validate SIRT1 expression, characterize its interaction with METTL3, and screen plant-derived compounds from Indian medicinal plants for modulatory potential. We aim to provide mechanistic insights and identify phytochemical scaffolds that could guide the development of new therapeutic strategies for cancer and metabolic disorders.

## 2. Materials and methods

### 2.1. Cloning and transformation

A DNA fragment encoding the human NAD-dependent protein deacetylase sirtuin-1 (SIRT1) protein (residues 183-665, UniProt accession number Q96EB6) was amplified from a human brain cDNA library using PCR with specific primers. The PCR was conducted on a Bio-Rad (T100) thermal cycler following this protocol: Step 1 at 95°C for 30 seconds, Step 2 at 55°C for 30 seconds, and Step 3 at 72°C for 90 seconds, with Step 2 repeated for a total of 35 cycles. The amplified product was then extracted from the gel, digested with NdeI and XhoI (TaKaRa), and cloned into the pET-28a expression vector (Novagen). The SIRT1 coding sequence was cloned into a recombinant plasmid and subsequently introduced into Rosetta competent cells to facilitate protein expression.

### 2.2. Protein Expression and Purification

Overnight cultivation at 37 °C in 5 mL of LB medium was initiated from a single bacterial colony. The next day, 5 mL of the culture was transferred into 100 mL of LB medium and incubated for approximately 3 hours. A 100 mL bacterial culture was transferred into 1 L of LB medium supplemented with 20 μg/mL kanamycin. Protein expression was induced by the addition of 0.4 mM isopropyl-β-D-thiogalactopyranoside (IPTG), followed by incubation at 16 °C for 20 hours. Cells were collected via centrifugation at 5000 rpm for 10 minutes, and the resulting pellet was resuspended in lysis buffer containing 20 mM Tris and 1 M NaCl, adjusted to pH 8.0. Lysis was performed by sonication at 4 °C, after which the lysate was clarified through centrifugation at 13,000 rpm for 30 minutes. The supernatant was subjected to an additional centrifugation step at 14,000 rpm for 30 minutes at 4 °C. Protein purification was carried out using a Ni-NTA column (QIAGEN), followed by size-exclusion chromatography on a Superdex 200 column (GE Healthcare). Analysis by SDS-PAGE revealed a distinct protein band with an approximate molecular mass of 54 kDa.

### 2.3. Isothermal Titration Calorimetry (ITC)

Isothermal titration calorimetry (ITC) assays were conducted at 20°C using a MicroCal PEAQ-ITC instrument (Malvern). METTL3, at a concentration of approximately 30 μM, was added to the sample cell, while SIRT1, at a concentration of around 300 μM, was placed in the syringe. The ITC experiment involved 19 injections: an initial 1 μL injection followed by 18 injections of 2 μL of the protein solution. The resulting heat data were analyzed using a one-site binding model through the MicroCal PEAQ-ITC Analysis Software provided by the manufacturer.

### 2.4. Retrieval and Protein Preparation

The three-dimensional crystal structures of SIRT1 bound with inhibitors Kaempferol, Catechin, Epicatechin, and Bisdemethoxycurcumin (PDB ID: 4I5I), as well as activators Ginsenosides and Carnosol (PDB ID: 5BTR), were obtained in PDB format from the Protein Data Bank. The protein structure was processed by eliminating all heteroatoms and water molecules with the help of PyMol and Discovery Studio Client. To refine the structure, key aspects such as missing hydrogen atoms, side-chain geometry, and incorrect bond orders were optimized and subjected to energy minimization using the GROMOS 43B1 force field in Swiss-PDB Viewer [22].

### 2.5. Preparation of Ligand

A comprehensive literature review identified 41 Indian medicinal plants and their phytochemical compounds, as determined by GC-MS (**Table S1**). The compounds were subsequently retrieved in 3D SDF format from the PubChem database. Ligands underwent energy minimization using the MMFF94 force field and the steepest descent optimization algorithm with 2000 steps in PyRx software prior to conducting molecular docking studies.

### 2.6. Molecular Docking Study

A docking-based computational analysis was carried out to explore how candidate compounds interact with the binding pocket of the target protein. Virtual screening with PyRx was conducted to pinpoint the amino acid residues of the target protein involved in ligand interactions [23]. For molecular docking, the protein and ligand structures in PDB format were processed in PyRx to generate the required PDBQT files. Redocking refers to the process of extracting the ligand molecule from the receptor model. To validate the docking results, the 4I5I structure was re-docked with its crystallographic inhibitors Kaempferol, Catechin, Epicatechin, and Bisdemethoxycurcumin. Similarly, 5BTR was re-docked with its activators, Ginsenosides and Carnosol. The re-docked poses were then compared with the ligand conformations in the crystal structure to assess any differences for docking the compounds against 4I5I; the center of the grid box was kept at X: 26.6289, Y: - 15.3706, Z: 22.7738, and dimensions were kept at X: 73.8785, Y: 79.6727, and Z: 51.0347 Å. Additionally, docking was performed against the activator 5BTR using a grid box with the center at X: −21.969, Y:68.4250, Z:4.4549, and dimensions at X:66.7105, Y:55.9884, and Z:64.7260 Å.

Ligand-protein binding strength was quantified in kcal/mol, with lower (negative) scores indicating stronger interactions [24]. The compounds were assessed through docking, and their binding strengths were ranked based on the most negative values, reflecting the highest affinity for the target [25],[26]. The interactions between the drug molecules and the active site residues of the M protein were confirmed by examining the results of both docking and re-docking at each binding position.

### 2.7. ADMET and Bioactivity Analysis

The ADME/T properties were predicted using the SwissADME server, which took SMILES (Simplified Molecular Input Line Entry System) notation of the compounds as input. The pharmacokinetic and pharmacodynamic characteristics, including metabolism, absorption, distribution, excretion, and toxicity, were assessed. The Molinspiration server was used to analyze the biological activities of the compounds. A bioactivity score greater than 0 indicates a biologically active compound, a score between 5.0 and 0 suggests moderate activity, and a score below −5.0 indicates biological inactivity.

### 2.8. Frontier molecular orbital

A quantum physics method called Density Functional Theory (DFT) may be able to accurately depict electron configurations in molecules. Molecular quantities that can be calculated in this manner include electronic properties, structural configurations, and energy levels. Density Functional Theory (DFT) at the B3LYP/6-31G(d) level of theory was used to assess the electronic characteristics of the chosen molecules in their singlet ground state under gas-phase conditions, ignoring solvent effects and external disturbances. The Gaussian computational software was used to optimize geometry and compute frontier molecular orbitals via the WebMO graphical user interface [27]. The DFT approach investigated a number of reactivity characteristics to evaluate molecular reactivity. The characteristics provided included chemical softness (ζ), chemical hardness (η), ionization potential, electronegativity (χ), electrophilicity (ϋ), and electronic potential (μ) [28]. A thorough assessment of the molecular reactivity was made possible by using the DFT technique to examine a variety of reactivity descriptors. The characteristics included ionization potential, electron affinity, chemical hardness (η), chemical softness (ζ), electronegativity (χ), electronic potential (μ), and electrophilicity (ω). These descriptors were computed using Koopmans’ theory, based on the energies of the frontier molecular orbitals [29].

### 2.9. Molecular Dynamics Simulation

Molecular Dynamics (MD) simulation is used to assess and confirm the structural flexibility and entropic effects of protein-ligand complexes in a simulated environment. Before conducting the production simulations, energy minimization and equilibration simulations were carried out to eliminate unfavorable interactions between the solute and solvent water molecules [30]. In the initial step of energy minimization, harmonic restraints were applied to the solute and lipid chains, while in the second step, all atoms were allowed to move freely [31]. Molecular dynamics simulations were performed using YASARA Dynamics software to validate the predictions made from the docking study [32]. The MMFF94 force field, commonly used for macromolecular systems, was applied in this simulation study [33]. Selisistat (EX527) was used as the control system to assess and compare the structural stability of the complexes. Long-range electrostatic interactions were calculated using the Particle Mesh Ewald (PME) method, while short-range van der Waals and Coulomb interactions were evaluated with a cutoff radius of 8Å [34]. The system’s overall environmental conditions were maintained at a temperature of 298 K, a pH of 7.4, and 0.9% NaCl concentration. The steepest descent algorithm was applied for system minimization [25].A Monte Carlo barostat was utilized to regulate pressure, while a Langevin thermostat was employed to control the temperature[35].The MD simulation trajectories were recorded at intervals of 100 ps with a time step of 1.25 fs [36]. To ensure simulation convergence, multiple parallel trajectories were performed for each complex[37]. The MD simulation was run for 100 ns to evaluate the root mean square deviation (RMSD), root mean square fluctuation (RMSF), radius of gyration (RG), solvent-accessible surface area (SASA), and the number of hydrogen bonds [34],[38–40]. The MD plots were generated using Matplotlib.

### 2.10. Free Energy Landscape Analysis

A free energy landscape (FEL) illustrates the potential energy of a system in relation to its pertinent variables [41], providing insights into the energy barriers and favorable states a molecule encounters during transitions between different conformations or states [42]. RMSD and RG values were calculated for each frame based on the simulation trajectories, generating a density matrix that defined a specific region within the conformational space. This density matrix, combined with the principles of statistical mechanics, was then used to determine the Gibbs free energy surface [43]. Using Matplotlib v3.7.2, comprehensive 2D and 3D visual plots were rendered for data analysis.

### 2.11. Probability Density Function

A probability density function was conducted to evaluate how structural properties varied along the trajectory, offering a detailed view of the complex’s dynamic behavior [44]. To represent a PDF analysis, the RG and RMSD histogram values were scaled accordingly. The probability density function of the variables was non-parametrically estimated using Gaussian kernel density estimation, which was implemented in scikit-learn v1.3. Using NumPy v1.25.2, arrays were consolidated and sorted for analysis.

## 3. Results and Discussion

### 3.1 Human SIRT1 NHL domain cloning, expression, and purification

SIRT1 (Sirtuin 1) is an NAD+-dependent histone deacetylase that regulates gene expression, metabolism, DNA repair, and stress response, playing key roles in aging, longevity, and cellular homeostasis. Its structure comprises four main domains (**Fig. 1**): the N-terminal Domain (NTD, 1–150), which is intrinsically disordered and modulated by post-translational modifications; the Catalytic Core Domain (CCD, 151–400), responsible for deacetylase activity; the C-terminal Regulatory Segment (CTR, 401–747), which regulates activity and stability; and the STAC-Binding Domain (SBD) within the NTD, which enhances deacetylase activity through allosteric activation.

**Fig. 1.**
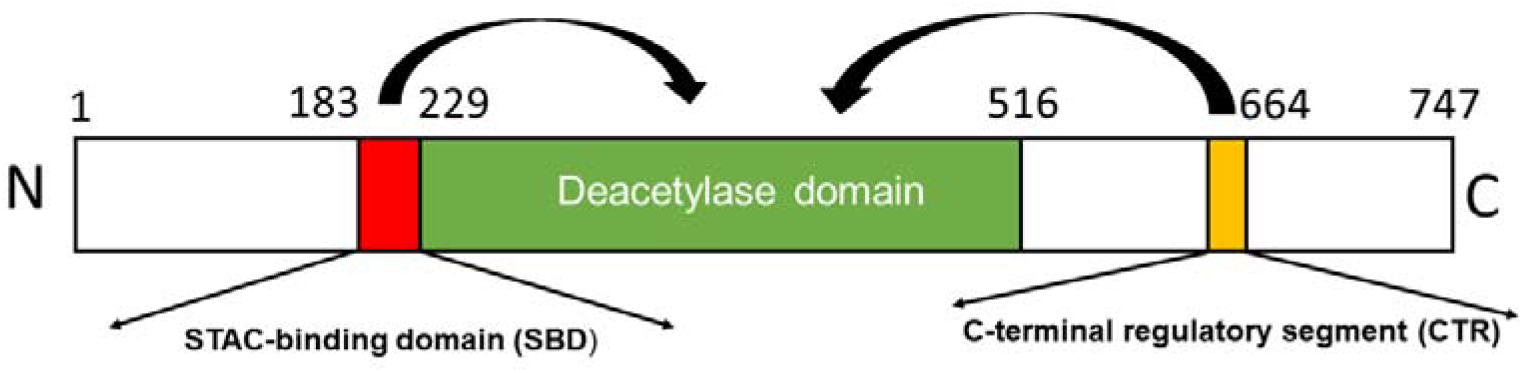
The SIRT1 protein consists of several distinct domains arranged from the N-terminus to the C-terminus. The N-terminal Domain (NTD) spans amino acids 1–150 and is intrinsically disordered, containing sites for post-translational modifications, including the STAC-binding Domain (SBD) involved in allosteric activation. The Catalytic Core Domain (CCD), located from amino acids 151–400, contains the active site responsible for the deacetylase activity of SIRT1. The C-terminal Regulatory Segment (CTR) spans residues 401–747 and plays a role in regulating SIRT1’s activity, stability, and localization through post-translational modifications and protein interactions. The color-coding in the figure highlights the specific positions and boundaries of each domain within the SIRT1 protein structure.

A DNA fragment encoding truncated human SIRT1 (residues 183–665, UniProt Q96EB6) was amplified from a human brain cDNA library (**Fig. 2**), cloned into the P28 expression vector, and transformed into Rosetta cells. Expression of the protein in soluble form was achieved, followed by purification using a two-step approach. Initial purification via Ni-NTA affinity chromatography enriched the protein, and further purification using size-exclusion chromatography (SEC) on a Superdex 200 column ensured homogeneity. The SEC profile showed a single peak at 15 mL, corresponding to a molecular mass of 54 kDa (**Fig. 3**), indicating a monomeric state in solution. SDS-PAGE analysis confirmed the protein purity exceeded 95%, demonstrating the effectiveness of the purification process and yielding a stable, homogeneous SIRT1 protein suitable for downstream studies.

**Fig. 2.**
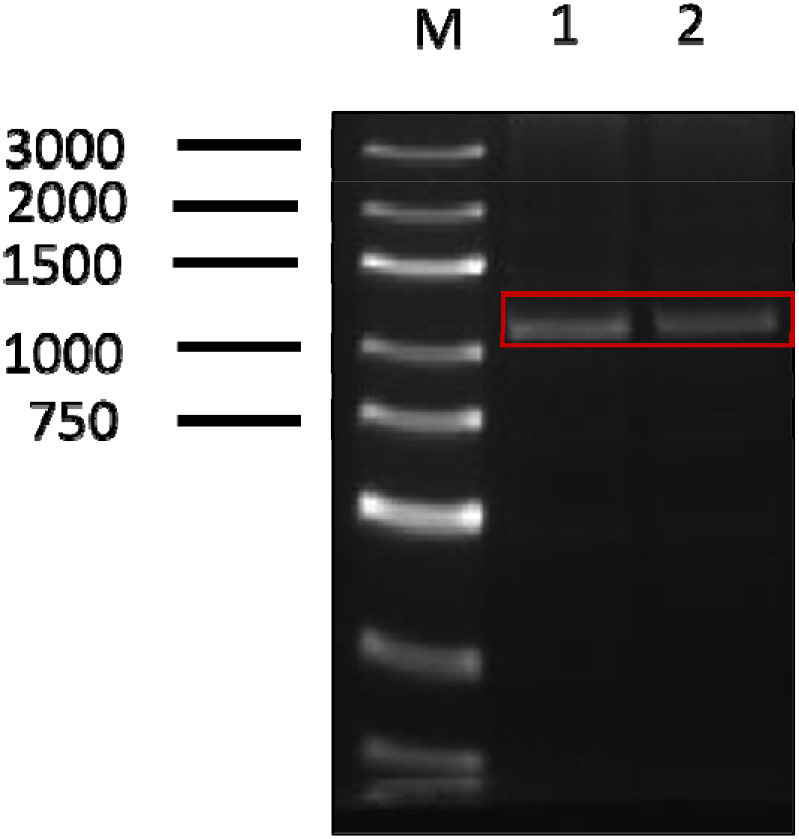
Agarose Gel Electrophoresis Results of Bacterial Culture PCR: Lanes 1 and 2 show the PCR results using bacterial culture as the template. The amplified hSIRT1 183-665 fragment is 1425 nt in length.

**Fig. 3.**
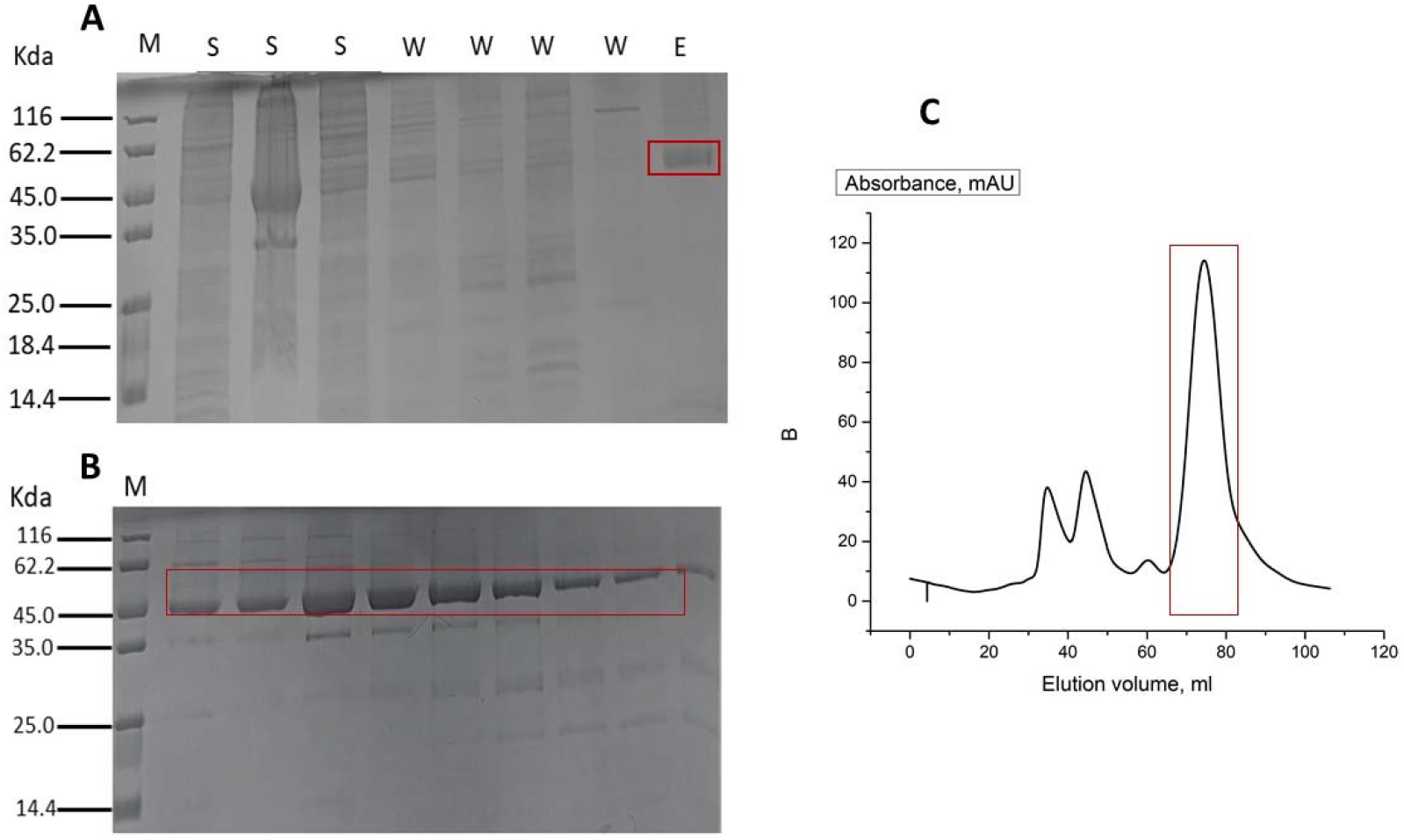
Purification of SIRT1 expressed in E. coli. (A) SDS-PAGE of Ni-NTA affinity chromatography showing supernatant (S), wash (W), and elution (E) fractions with the target protein (∼54 kDa). (B) SDS-PAGE confirming purity after size-exclusion chromatography (SEC) using a HiLoad 16/60 Superdex 200 column. (C) SEC chromatogram highlighting the elution peak of purified SIRT1.

### 3.2 Interaction between SIRT1 and METTL3

The interaction between SIRT1 and METTL3 is a key regulatory mechanism in cancer metastasis, particularly influencing post-translational modifications that govern RNA methylation and protein localization. SIRT1, a class III histone deacetylase, exerts its function by deacetylating METTL3 at lysine 177 (K177), a critical step that promotes METTL3’s nuclear localization. Once METTL3 enters the nucleus, it serves as the catalytic core of the m6A methyltransferase complex, which modifies specific mRNAs via m6A methylation. This epigenetic modification significantly impacts mRNA stability, splicing, translation, and expression of genes involved in tumorigenesis and metastasis. Elevated m6A levels have been shown to enhance cancer cell invasion, survival, and proliferation, further underscoring the oncogenic role of METTL3 in cancer progression. To understand this interaction at the molecular level, both computational docking using HADDOCK and experimental thermodynamics using Isothermal Titration Calorimetry (ITC) were performed (**Fig. 4**), offering complementary insights. The HADDOCK docking simulations identified Cluster 1 as the most favorable binding configuration between SIRT1 and METTL3, with a HADDOCK score of - 126.5. This indicates a highly stable interaction. The large buried surface area (BSA) of 2854.3 Å² in this cluster highlights an extensive interface between the two proteins, suggesting a robust and stable interaction. Moreover, electrostatic interactions, with an energy contribution of −481.7 kcal/mol, play a dominant role in stabilizing the SIRT1-METTL3 complex. These charged residues likely form a network of interactions that reinforce the binding affinity, critical for the biological activity of the complex. The ITC data corroborate the docking results, providing experimental confirmation of the strong binding affinity between SIRT1 and METTL3. The heat change recorded over time during the titration of METTL3 into SIRT1 indicates a high-affinity interaction, with rapid saturation occurring at a 1:1 molar ratio. The binding isotherm in the lower panel of the ITC data shows a steep rise followed by a plateau, consistent with a tight interaction (**Fig. 5**). The negative enthalpy change (ΔH) observed in the ITC experiment further supports the exothermic nature of the interaction, indicative of strong electrostatic forces, mirroring the dominant electrostatic contribution observed in the HADDOCK analysis. The stoichiometry of 1:1 derived from ITC also aligns with the docking data, supporting a direct one-to-one interaction between SIRT1 and METTL3.

**Fig. 4.**
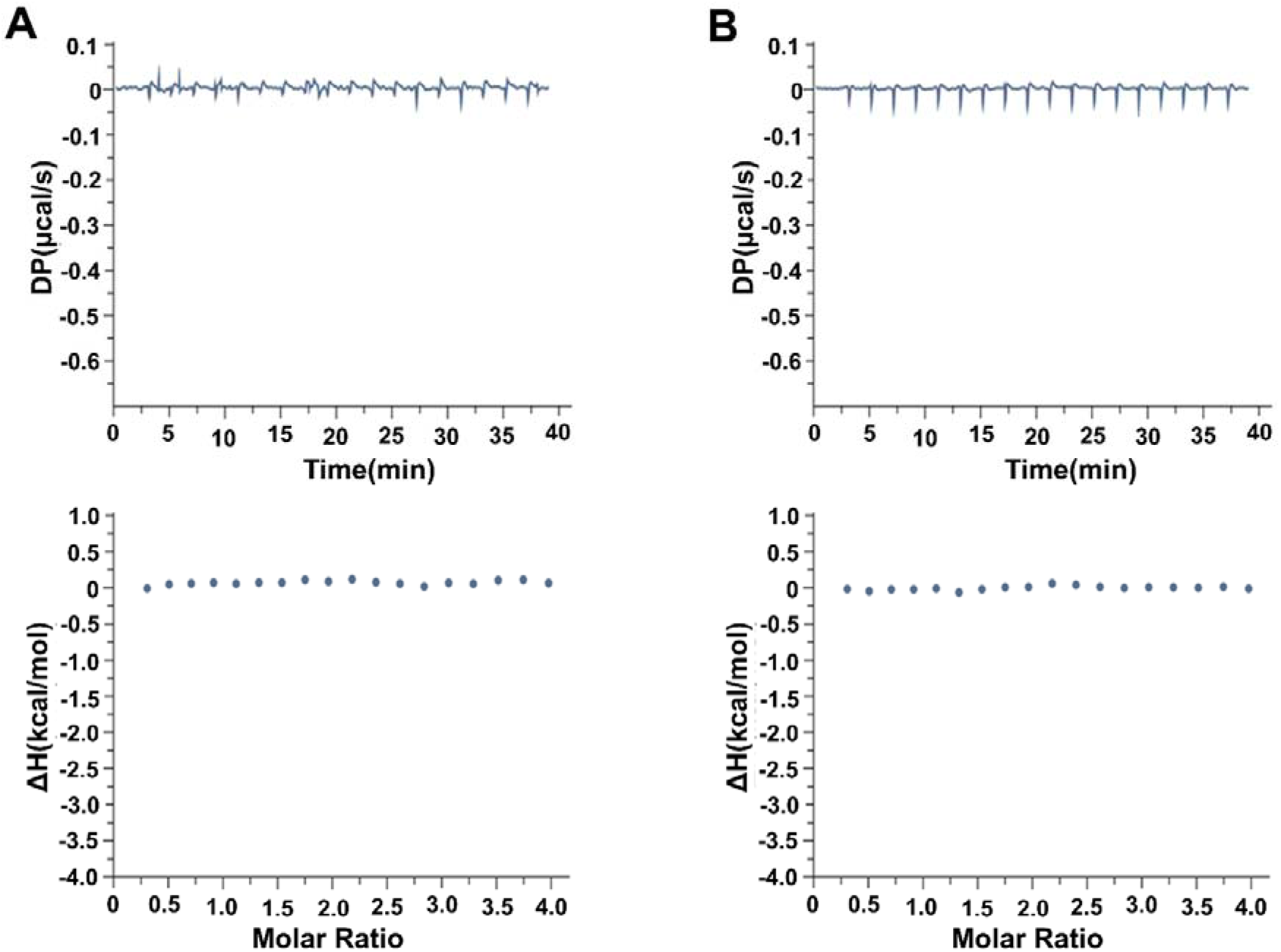
Isothermal titration calorimetry (ITC) analysis of SIRT1 interaction with wild-type METTL3 and the METTL3 K177AC mutant. The top panels of Figures A and B depict the differential power (DP) versus time during the titration of SIRT1 into wild-type METTL3 and the METTL3 K177AC mutant, respectively. The lower panels display the integrated heat changes as ΔH (enthalpy change) plotted against the molar ratio. Both wild-type METTL3 and the K177AC mutant show no detectable binding or heat change, suggesting that either post-translational modifications or specific residue states are required for interaction with SIRT1. The acetylation mimic at K177 (K177AC) further highlights the importance of K177 in mediating METTL3’s interaction with SIRT1.

**Fig. 5.**
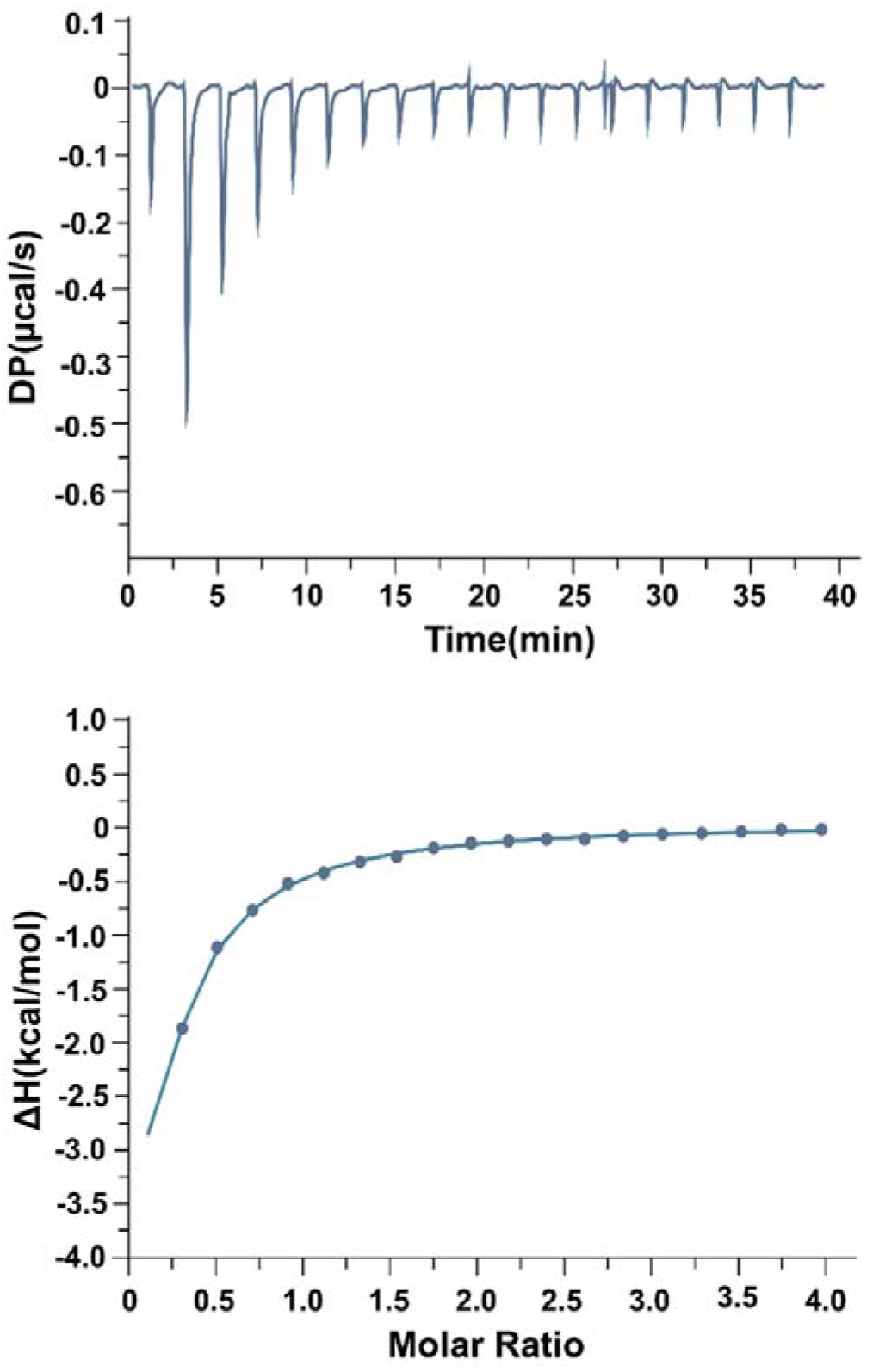
ITC analysis of SIRT1 interaction with the METTL3 K177Q mutant. The top panel represents the DP signal during SIRT1 titration into the METTL3 K177Q mutant, which mimics a constitutively deacetylated state. The lower panel illustrates the binding isotherm, with a noticeable enthalpy change indicating binding. These results confirm that deacetylation at K177 by SIRT1 facilitates the interaction, emphasizing the critical role of K177 in this regulatory mechanism.

The combined insights from HADDOCK and ITC provide a detailed picture of the SIRT1-METTL3 interaction, essential for their function in cancer metastasis. The deacetylation of METTL3 by SIRT1 at K177 enables METTL3 to translocate into the nucleus, where it facilitates m6A methylation of mRNAs that regulate metastasis and tumorigenesis. The computational docking results reveal a strong electrostatic interaction between these proteins, while the ITC results experimentally validate this by showing a high binding affinity and exothermic interaction profile. The consistency between these results emphasizes the biological relevance of the SIRT1-METTL3 interaction in cancer biology.

### 3.3 Evaluation of Selected Compounds Targeting SIRT1 Using Molecular Docking

In molecular docking, the inhibitors’ protein docking (PDB ID: 4I5I) and activators-proteins (PDB ID: 5BTR) have been performed to find lead compounds with higher binding affinity. The structure of the SIRT1 protein has three structural regions: the 183-229 residues N-terminal region, the 229-516 residues contain the area of the catalytic core, and the 641-665 residues known as the C-terminal region. The N-terminal region acts as the sirtuin activation compound binding site, and the core catalytic region plays a pivotal role in the sirtuin inhibitory binding site [4,5].

To identify the potential inhibitors, a blind docking strategy comprising high-throughput virtual screening (HTVS) was employed from an Indian medicinal plants database. This approach identified seven inhibitor compounds with binding affinities stronger than the reference inhibitor Selisistat (−8.1 kcal/mol) (**Table 1**): Catechin and Bisdemethoxycurcumin (−9.4 kcal/mol), Epicatechin (−9.2 kcal/mol), Kaempferol (−9.1 kcal/mol), Beta-eudesmol (−9.0 kcal/mol), Tristin (−8.7 kcal/mol), and Champacol (−8.4 kcal/mol). The top inhibitors, including Catechin, Bisdemethoxycurcumin, Epicatechin, and Kaempferol, exhibited consistent interactions with SIRT1’s catalytic residues (**Fig. 6**). Hydrogen bonds with His363 (a key residue in the deacetylase mechanism) and Gln345 stabilized their positioning, while π-π stacking interactions with Phe297 and Phe273 anchored these ligands within the hydrophobic NAD-binding pocket. These observations align with Zhao et al [45] who emphasized the role of hydrophobic interactions in stabilizing SIRT1 inhibitors deep within the active site. Structural analysis revealed high complementarity between the ligands and the catalytic cavity, mimicking the binding pose of the co-crystallized ligand (CL) and the known inhibitor. Notably, Catechin, Epicatechin, and Kaempferol bind with hydrogen bonding within the active sites Arg274, His363, Ser442, Gln345, and Val412. Cathecine binds with the active sites Arg274, His363, Val412, and Gln345 via its oxygen atom to form a hydrogen bond and also forms Pi-cation, Pi-Pi Stacked, and alkyl bonds with the active sites Arg274, Phe273, and Val445, respectively. Epicatechin forms a remarkable number of hydrogen bonds with the active sites Ala262, Phe414, Ser442, Ser441, and Gly261, with bond distances of 2.403, 2.98, 2.634, 2.514, and 2.625, respectively. Bisdemethoxycurcumin forms a hydrogen bond with the active sites Gln345, Asp272, and Gly261. Kaempferol forms hydrogen bonds, Pi-cation, Pi-Pi Stacked, and Pi-Alkyl bonding with the active site of the SIRT1 protein. Hydrogen bonding occurs with the active site residues Arg274, His363, Ser442, Gln345, Val412, and His363. The flexible loop region adjacent to the catalytic site adopted a conformation reminiscent of NAD-bound states, a phenomenon previously linked to NAD - induced structural rearrangements. Previous findings underscore the potential of natural compounds to act as competitive SIRT1 inhibitors, blocking the NAD-binding site through hydrophobic anchoring (Phe297/Phe273) and residue-specific H-bonding (His363, Gln345, Asp348) while stabilizing catalytically inactive conformations [46].

**Fig. 6.**
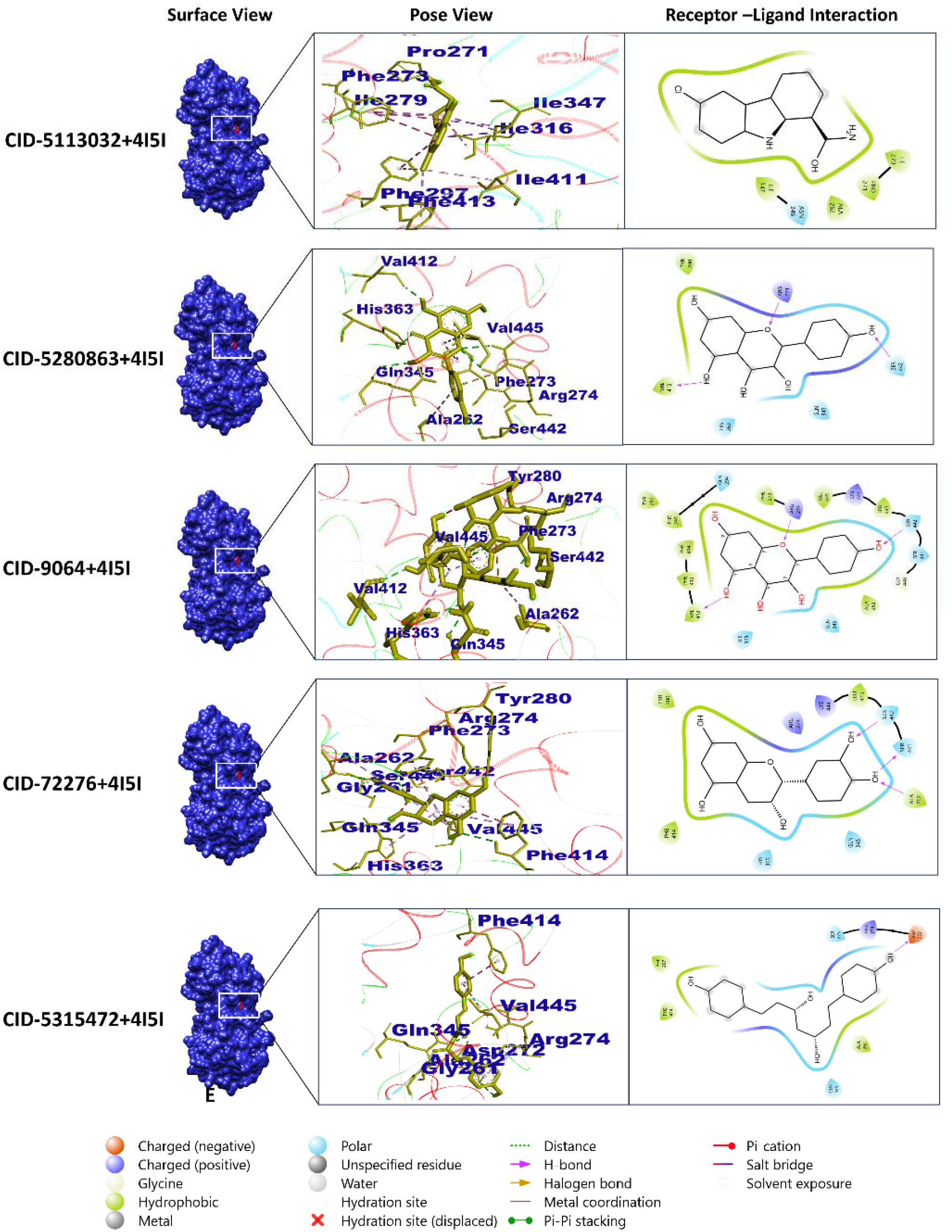
Molecular docking analysis of SIRT1 protein with four selected inhibitor compounds and a control: Selisistat (5113032), Kaempferol (5280863), Catechin (9064), Epicatechin (72276), and Bisdemethoxycurcumin (5315472). The interactions are illustrated using surface view, pose view, and receptor-ligand interaction diagrams for each compound. Surface View: Represents the SIRT1 protein as a molecular surface in blue, highlighting the binding pocket where the respective compounds interact. Pose View: Displays the binding residues of the compounds (shown in blue) within the binding pocket of SIRT1, emphasizing their spatial orientation and fit. Receptor–Ligand Interaction Diagram: Shows the specific interactions between the SIRT1 protein and each compound, including hydrogen bonds, hydrophobic interactions, and π-π stacking interactions, represented with corresponding labels and colors. (Catechin): Demonstrates key binding interactions within the active site. (Epicatechin): Shares a similar interaction pattern as Catechin with subtle differences in bond specificity. (Kaempferol): Exhibits extensive hydrogen bonding and aromatic interactions with the binding pocket. (Bisdemethoxycurcumin): Shows significant hydrophobic and aromatic stacking interactions due to its planar structure.

**Table 1.**
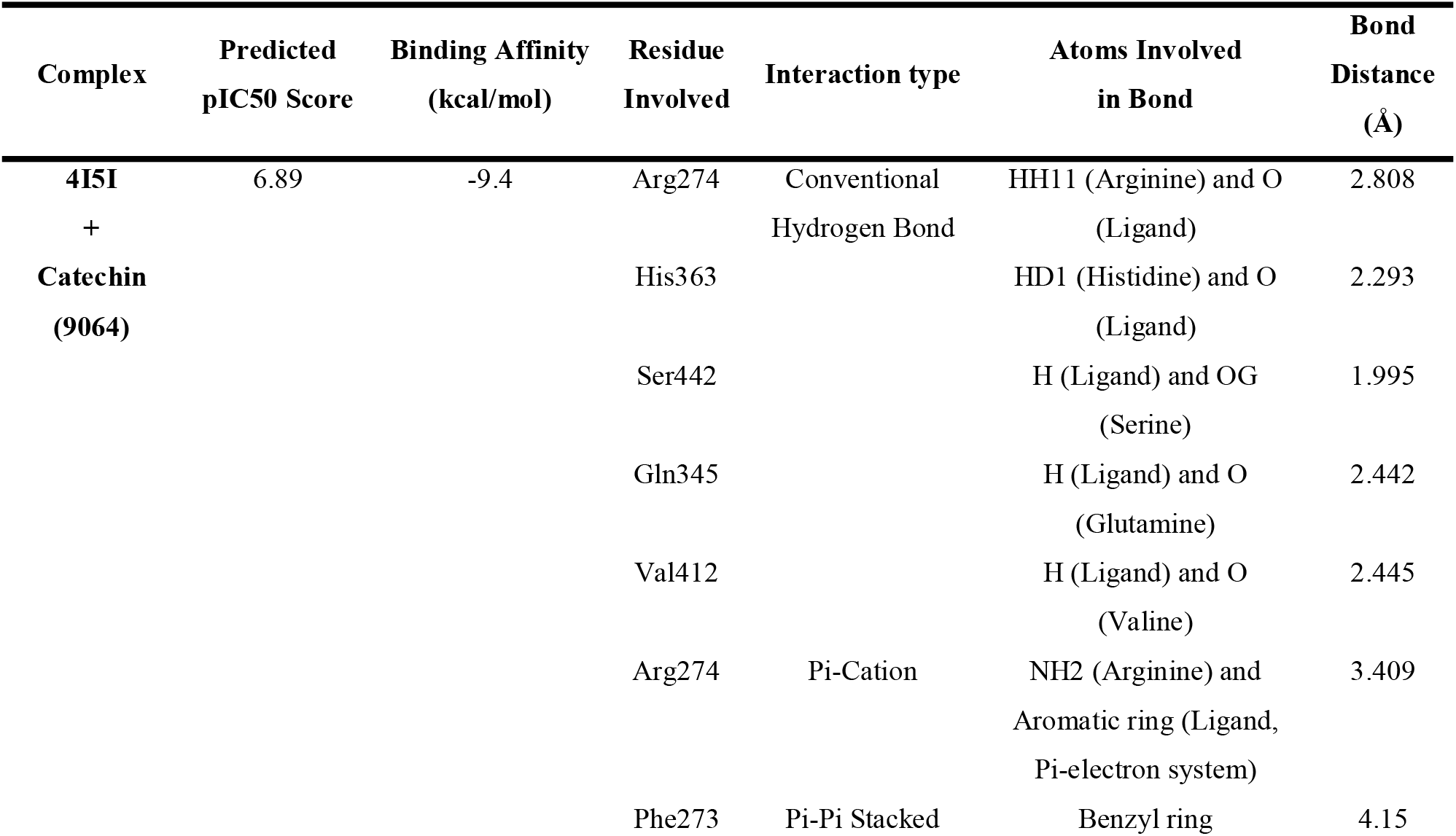
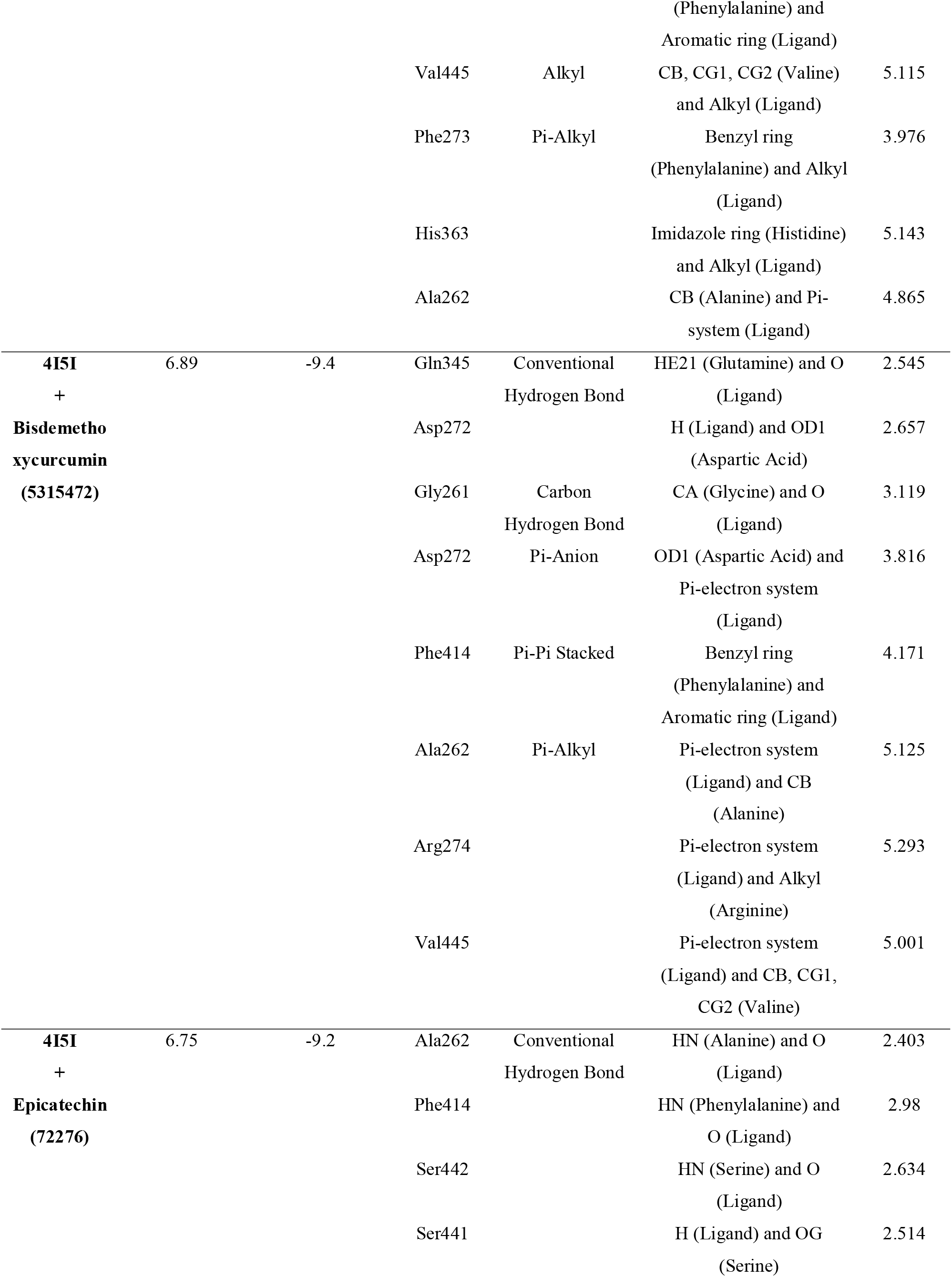
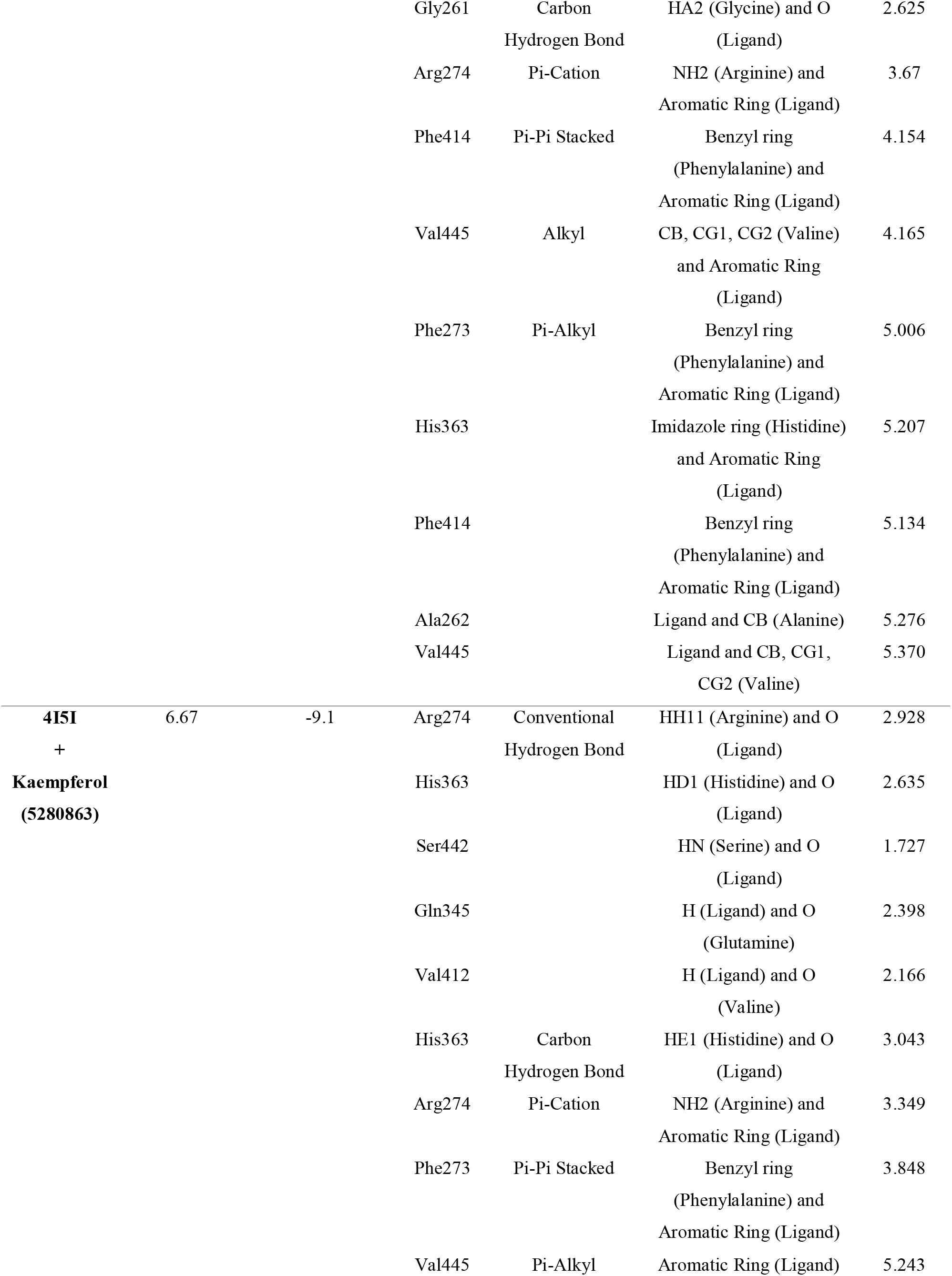
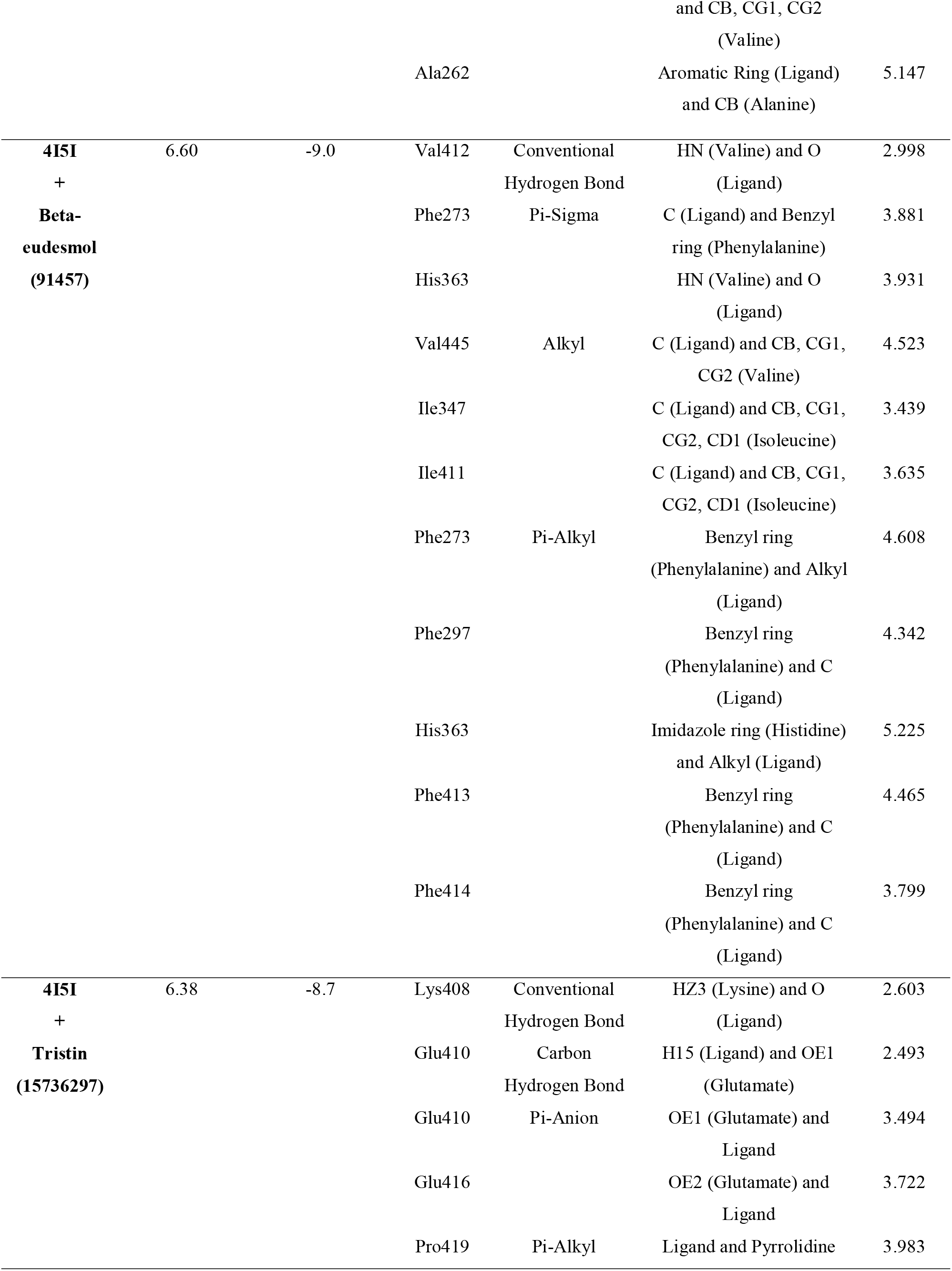
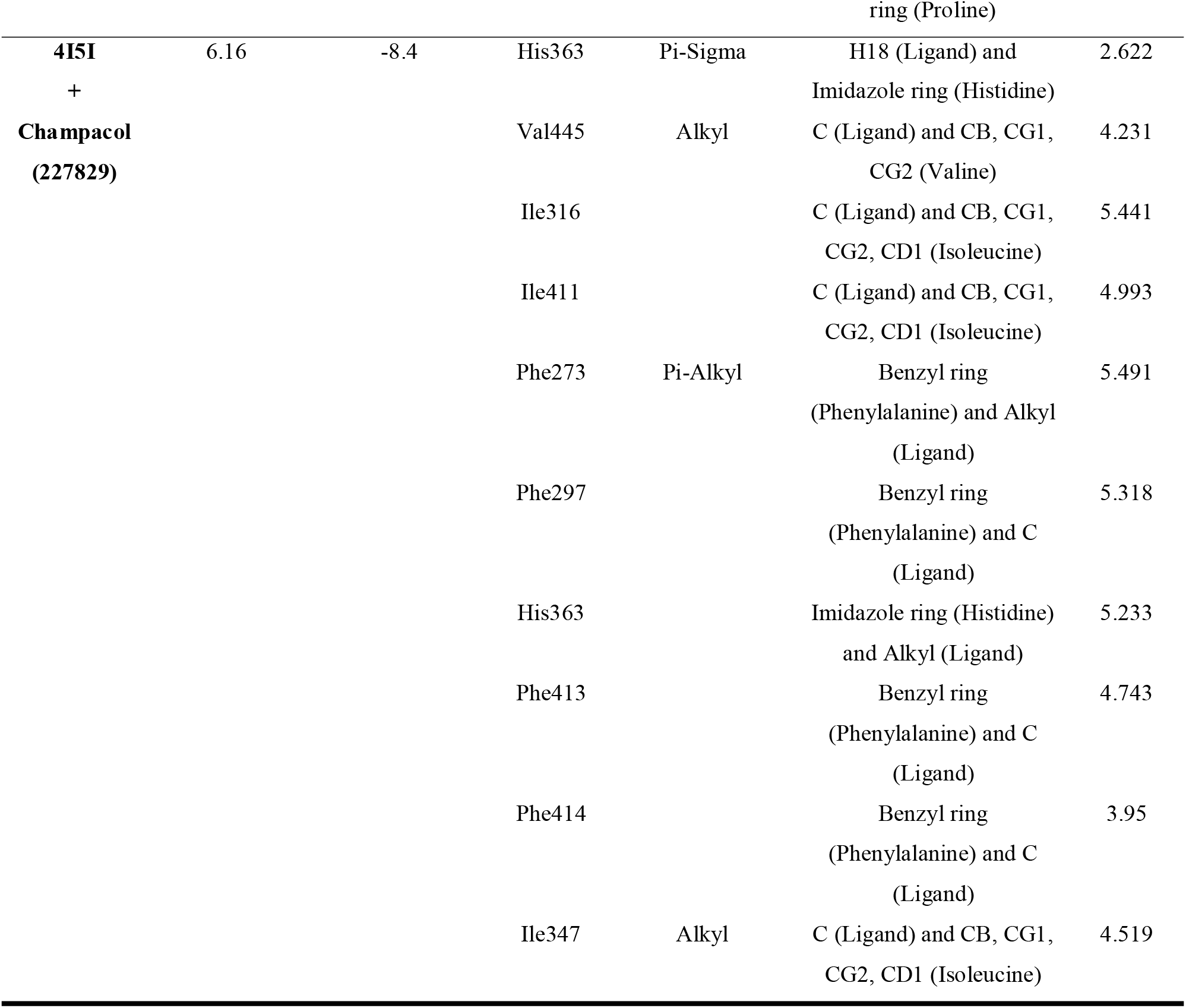
Binding affinity, predicted IC50 score, and non-bond interactions between the SIRT1 protein and the top 7 inhibitor compounds.

To demonstrate how the protein (PDB ID: 5BTR) and activator chemicals interact, another docking study was conducted (**Table 2**). The activator lead compounds were identified according to their binding affinity score, bonding interactions to the allosteric site of the protein, and bonding distances. The activator’s docking procedure occurred according to the previous literature [47–49]. The binding residues identified in the allosteric binding region predominantly consisted of aromatic and charged residues, suggesting that these activators were stabilized by alkyl, electrostatic, and stacking interactions (**Fig. 7**). Ginsenosides show a −9.6 kcal/mol binding affinity and form alkyl interactions within Pro212, Arg446, Ile223, Pro211, Leu450, Pro447, and Phe414 active sites; the nitrogen atom of the compound binds with the protein residue. Another lead compound, Carnosol, exhibits a −8.1 binding affinity score; the benzyl ring of the Phe414 residue binds with the compound and forms a Pi-Pi stacked bond. The beta carbon (CB) and gamma carbon (CG) of Pro212, Ile223, and Leu202 residues of the protein form alkyl bonding with the compound. Berberine, Fisetin, and Apigenin show −8.2, −8.5, and −8.0 kcal/mol binding affinity, respectively. Berberine binds to the allosteric site of the protein, specifically the Pro212 and Lys304 residues, to form an alkyl bond. Fisetin binds to Pro419, and Apigenin binds to Pro212 and Pro291 to form a Pi-alkyl interaction.

**Fig. 7.**
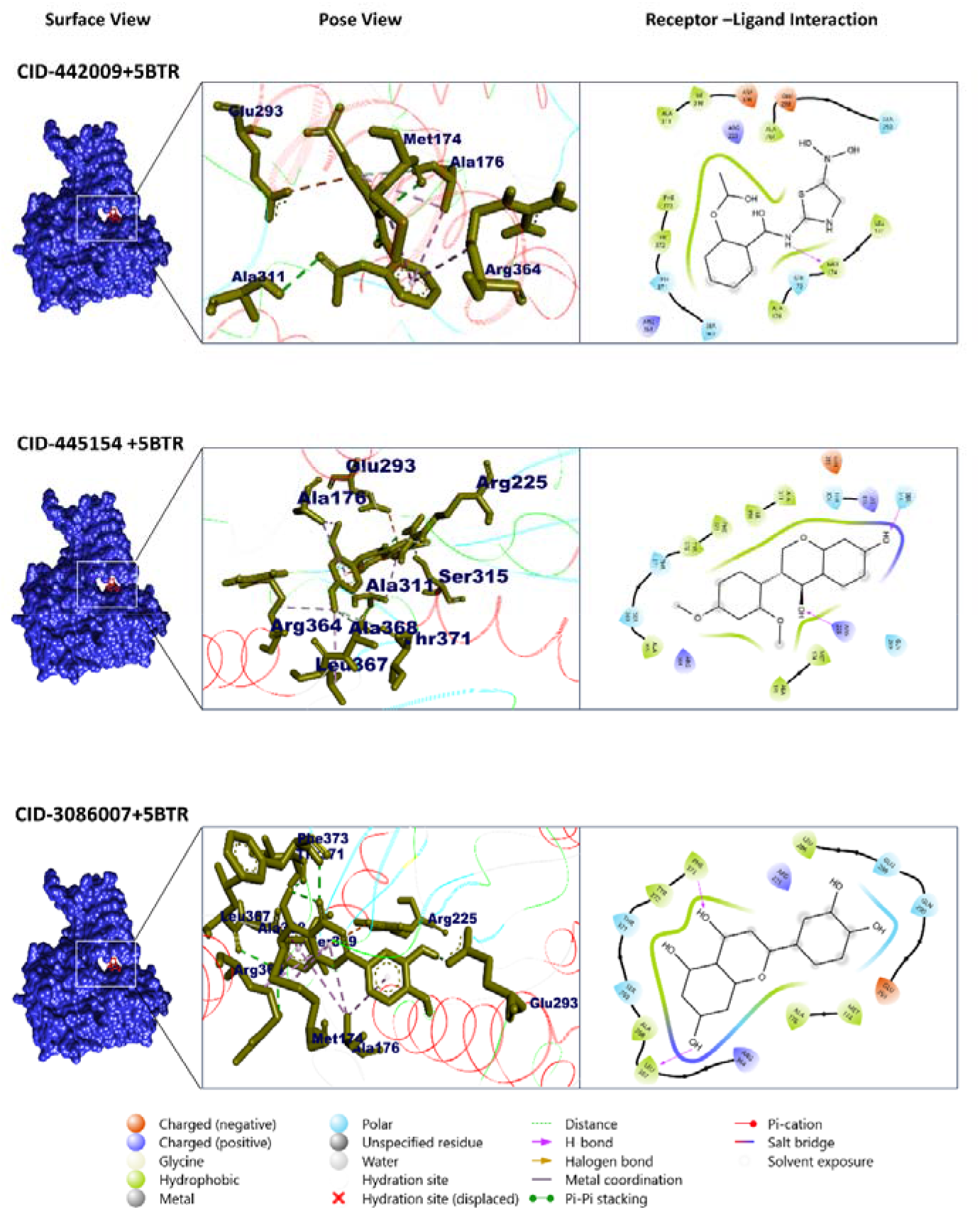
Molecular docking analysis of SIRT1 protein with two selected activator compounds and a control compound: Carnosol (CID 442009), Resveratrol (CID 445154), and Ginsenosides (CID 3086007). The interactions are illustrated using surface view, pose view, and receptor-ligand interaction diagrams for each compound. Surface View: Represents the SIRT1 protein as a molecular surface in blue, highlighting the binding pocket where the respective compounds interact. Pose View: Displays the binding residues of the compounds (shown in blue) within the binding pocket of SIRT1, emphasizing their spatial orientation and fit. Receptor–Ligand Interaction Diagram: Shows the specific interactions between SIRT1 protein and each compound, including hydrogen bonds, hydrophobic interactions, and π-π stacking interactions, represented with corresponding labels and colors. (Catechin): Demonstrates key binding interactions within the active site.

**Table 2.**
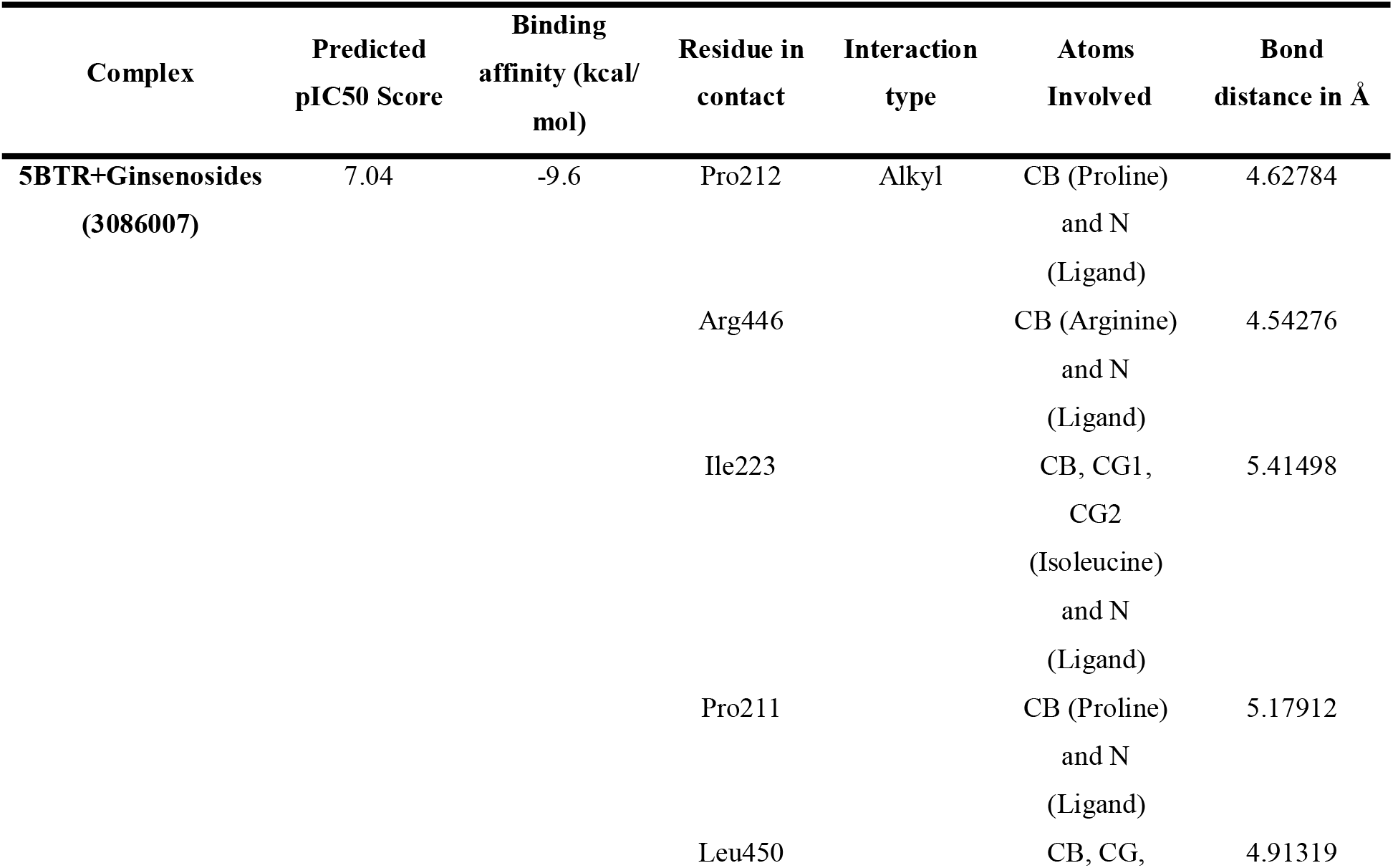
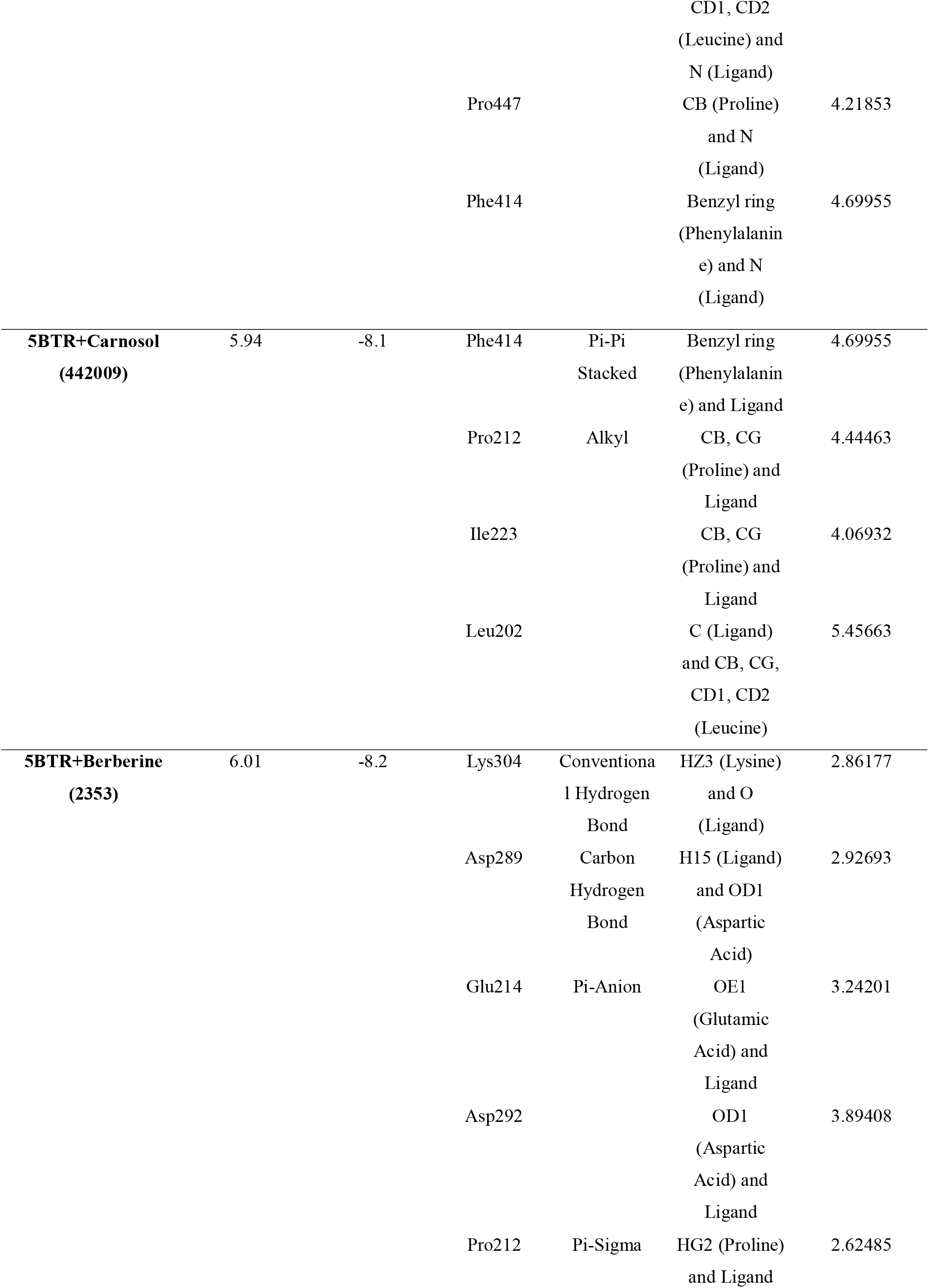
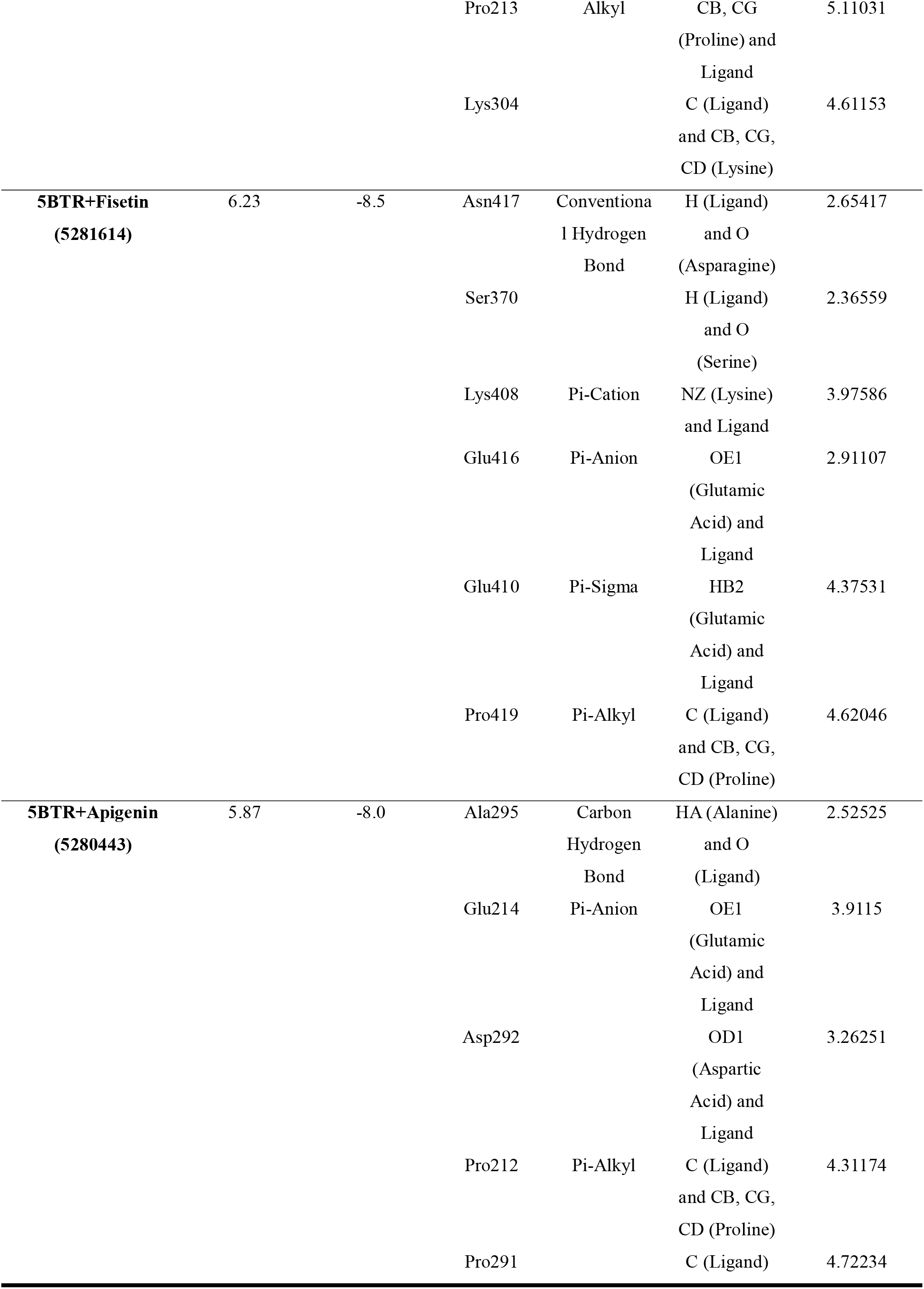
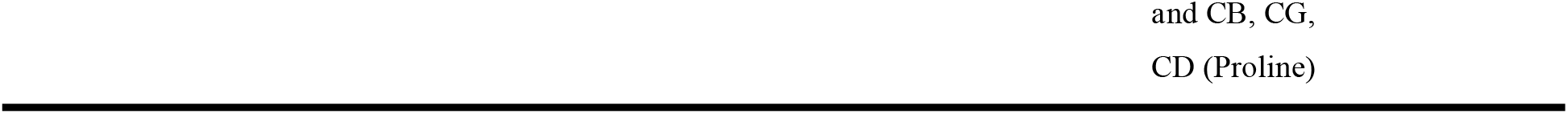
Binding affinity, predicted IC50 score, and non-bond interactions between SIRT1 protein and the top 5 lead activator compounds.

### 3.4 ADME/Tox and bioactivity analysis

Evaluating safety concerns, including toxicities and adverse drug reactions, is a critical aspect of drug development. While clinical trials are mandatory, computational predictive models for various toxicities have greatly enhanced the drug design process [50]. In activator compounds, Ginsenosides and Carnosol both are typically the same in results of physicochemical properties, water solubility, lipophilicity, and pharmacokinetics (**Table 3**), but for toxicity class, Ginsenosides shows better results than Carnosol.

**Table 3.**
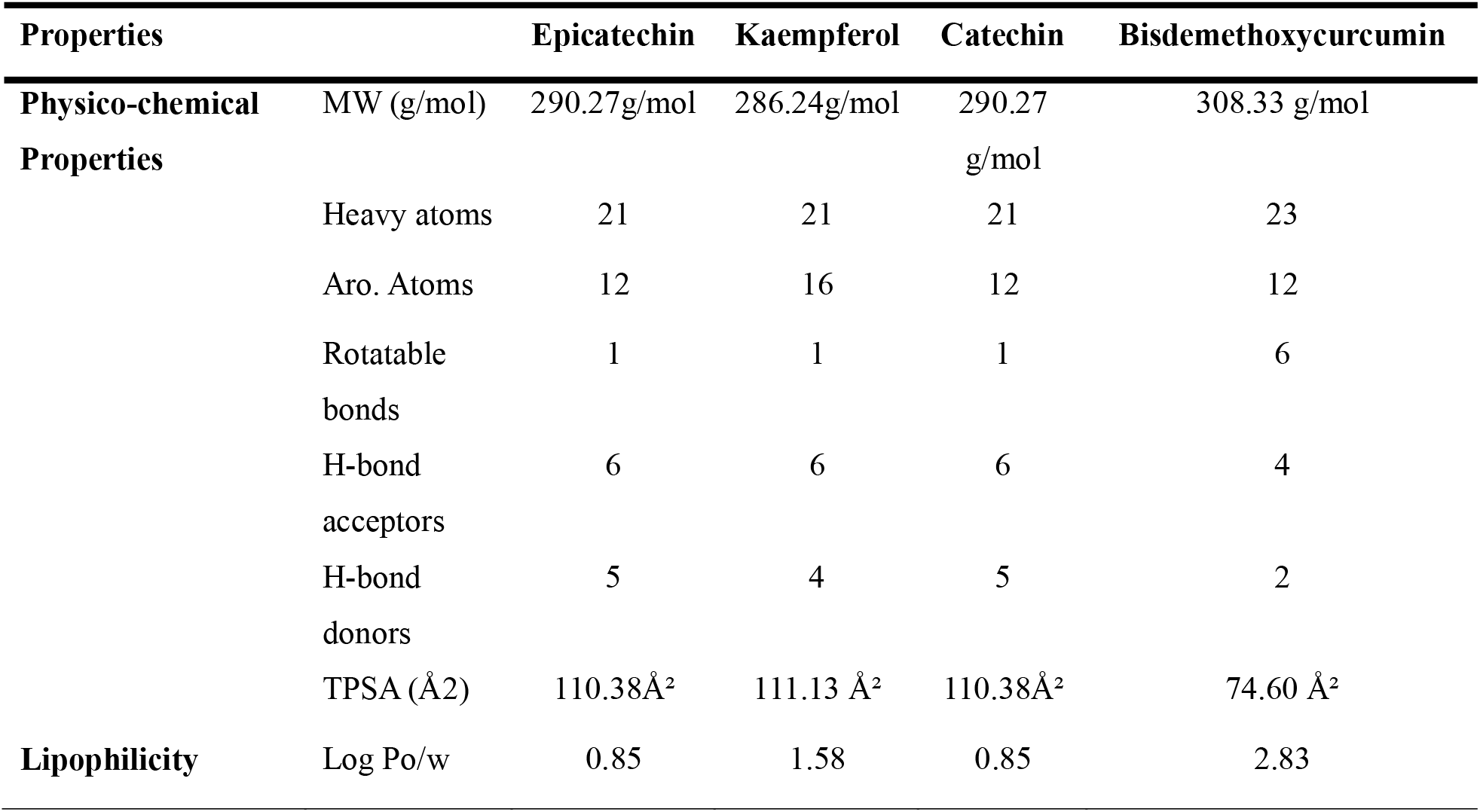
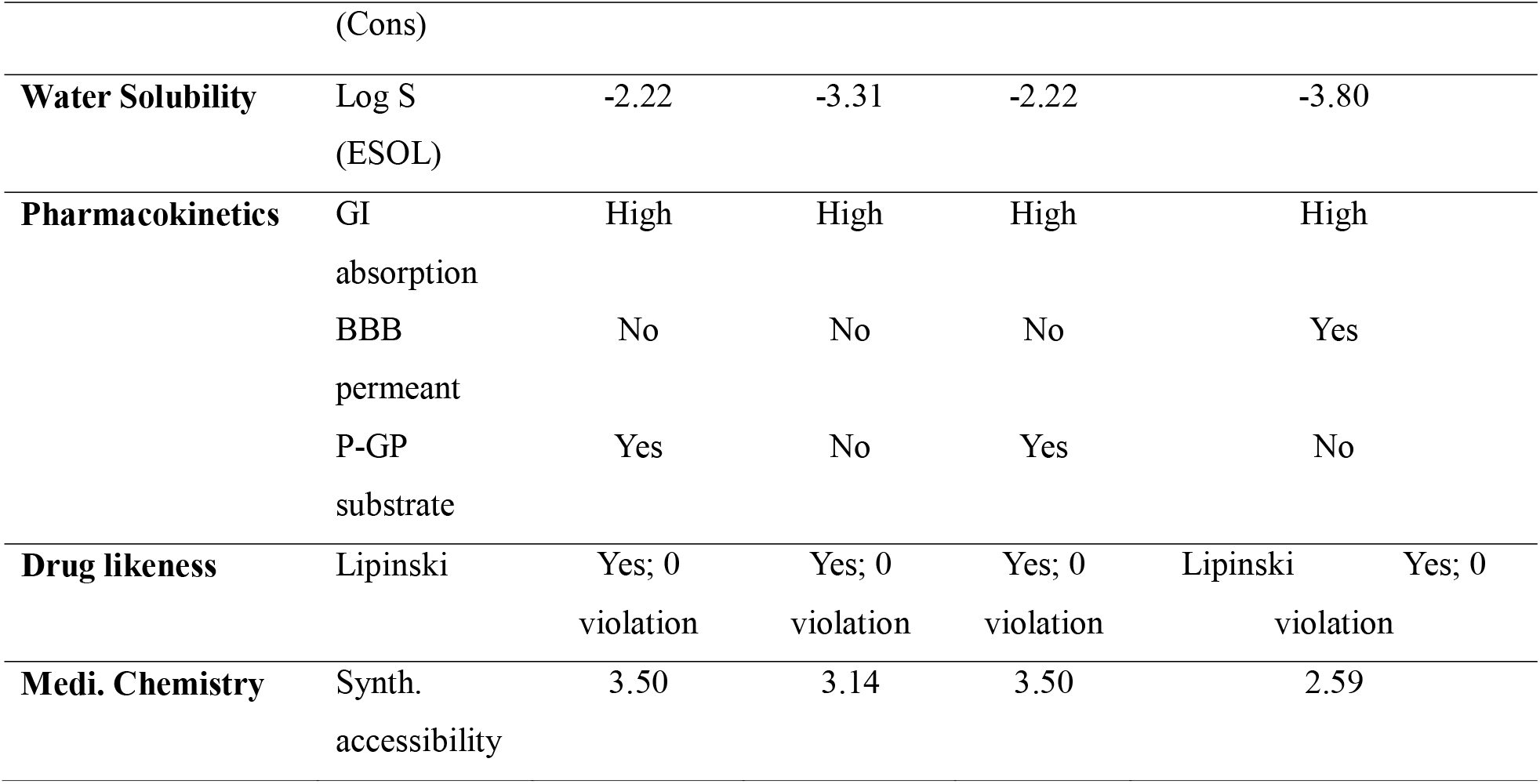
Pharmacological properties analysis of top four lead inhibitor compounds shows drug likeness characteristics.

In addition, the high rate of failure of drug candidates during clinical trials is primarily attributed to toxicity, thereby emphasizing its role as a key determinant in drug development [51]. The toxicity assessment encompassed multiple properties such as hepatotoxicity, carcinogenicity, immunotoxicity, mutagenicity, cytotoxicity, oral rat acute toxicity, LD50 value, and toxicity class for four inhibitors and two activators, with the corresponding results summarized in **Table 4** and **Table 6**, respectively. These computational methods have been employed in the construction of classifiers and regression frameworks aimed at estimating LD50 and categorizing compounds based on toxicity levels [52,53]. All the tested compounds were inactive in organ toxicity endpoints. Predicted LD50 of Epicatechin, Kaempferol, Catechin, and Bisdemethoxycurcumin were 10000, 3919, 10000, and 2560 mg kg^−1^. Ginsenosides and Carnosol were 2260 and 1500 mg kg^−^ ^1,^ making them nearly non-toxic to biological systems. Catechin and epicatechin, compared with Kaempferol and Bisdemethoxycurcumin, showed advantageous pharmacokinetic, biological, and toxicity properties, implying a minimal likelihood of unsuccessful outcomes in clinical testing. In terms of activators, Ginsenosides show better results in toxicity class analysis than Carnosol.

**Table 4.** Toxicological profile of the inhibitor compounds shows that inactive hepatotoxicity, carcinogenicity, immunotoxicity, mutagenicity and cytotoxicity.

| Endpoints | Target | Kaempferol | Catechin | Epicatechin | Bisdemethoxycurcumin |
| --- | --- | --- | --- | --- | --- |
| **Organ toxicity and toxicity endpoints** | Hepatotoxicity | Inactive | Inactive | Inactive | Inactive |
|  | Carcinogenicity | Inactive | Inactive | Inactive | Inactive |
|  | Immunotoxicity | Inactive | Inactive | Inactive | Inactive |
|  | Mutagenicity | Inactive | Inactive | Inactive | Inactive |
|  | Cytotoxicity | Inactive | Inactive | Inactive | Inactive |
|  | LD50 (mg kg<sup>-1</sup>) | 3919 | 10000 | 10000 | 2560 |
|  | Toxicity class | 5 | 6 | 6 | 5 |

These methods offer significant advantages, such as high accuracy and the ability to assess potential risks before a compound is synthesized [54]. To determine the effectiveness and safety of lead compounds, toxicity evaluations and pharmacokinetic properties must be examined [55]. The four lead compounds were further assessed for their pharmacological activity, a key factor in understanding their behavior within the human body, using ADME/T technology, which has recently been extensively reviewed for its applications [50]. The inhibitor compounds (Epicatechin, Kaempferol, Catechin, and Bisdemethoxycurcumin) along with the activator compounds (Ginsenosides and Carnosol) were systematically evaluated with respect to key pharmacokinetic and toxicity parameters, including Central Nervous System (CNS) permeability, hepatotoxicity, P-glycoprotein inhibition, and Cytochrome P450 (CYP) inhibition, among others. Central Nervous System (CNS) permeability denotes the ability of a drug to traverse the semipermeable blood–brain barrier, a physiological structure designed to safeguard the CNS from potentially harmful substances. A CNS permeability value greater than −4 is generally considered indicative of a compound’s ability to cross the blood–brain barrier.

The molecular weights of Epicatechin, Kaempferol, Catechin and Bisdemethoxycurcumin were 290.27, 286.24, 290.27, 308.33 g/mol respectively. The molecular weights of Ginsenosides and Carnosol were 444.73 and 330.42 g/mol respectively (**Table 5**). All four inhibitors are in agreement with Lipinski and Ghoses’ rule [56] of drug-likeness. In terms of activators Ginsenosides and Carnosol both are in agreement with Lipinski and Ghoses’ rule of drug-likeness. Consequently, Catechin, Epicatechin showed better results in physiochemical properties, water solubility, lipophilicity, pharmacokinetics and toxicity parameters concerning drug-likeness compared to Kaempferol, Bisdemethoxycurcumin.

**Table 5.** Pharmacological properties analysis of top two lead activator compounds.

| Properties |  | Ginsenosides | Carnosol |
| --- | --- | --- | --- |
| Physico-chemical Properties | MW (g/mol) | 444.73 g/mol | 330.42 g/mol |
|  | Heavy atoms | 32 | 24 |
|  | Aro. Atoms | 0 | 6 |
|  | Rotatable bonds | 4 | 1 |
|  | H-bond acceptors | 2 | 4 |
|  | H-bond donors | 2 | 3 |
|  | TPSA (Å<sup>2</sup>) | 40.46 Å<sup>2</sup> | 66.76 Å<sup>2</sup> |
| Lipophilicity | Log Po/w (Cons) | 6.75 | 3.72 |
| Water Solubility | Log S (ESOL) | -7.71 | -4.77 |
| Pharmacokinetics | GI absorption | Low | High |
|  | BBB permeant | No | Yes |
|  | P-GP substrate | No | Yes |
| Drug likeness | Lipinski | Yes; 1 violation: MLOGP>4.15 | Yes; 0 violation |
| Medi. Chemistry | Synth. accessibility | 5.98 | 4.88 |

**Table 6.** Toxicological profile of the inhibitor compounds shows that inactive hepatotoxicity, carcinogenicity, mutagenicity, and cytotoxicity, but active immunotoxicity.

| Endpoints | Target | Ginsenosides | Carnosol |
| --- | --- | --- | --- |
| **Organ toxicity and toxicity endpoints** | Hepatotoxicity | Inactive | Inactive |
|  | Carcinogenicity | Inactive | Inactive |
|  | Immunotoxicity | Active | Active |
|  | Mutagenicity | Inactive | Inactive |
|  | Cytotoxicity | Inactive | Inactive |
|  | LD50 (mg kg<sup>-1</sup>) | 2260 | 1500 |
|  | Toxicity class | 5 | 4 |

### 3.5 Frontier molecular orbital study

In quantum chemistry, the Frontier Molecular Orbital (FMO) analysis method is employed to investigate the geometries, energy sources, and electron distributions associated with two molecular orbitals: the lowest unoccupied (LUMO) and the highest occupied (HOMO). Molecular orbital behavior, particularly in organic molecules, may be better understood using this approach. The reactivity of molecular orbitals was assessed by analyzing the HOMO-LUMO levels, and it has been shown that molecules with a minimal or insignificant HOMO-LUMO gap generally display increased activity potential. Possible activators and inhibitors were the focus of this investigation at the subatomic level. The electron affinity and ionization potential of inhibitors such as Epicatechin, Kaempferol, Catechin, and Bisdemethoxycurcumin, and activators such as Ginsenosides and Carnosol were evaluated using FMO calculations. Key electronic properties, including ionization potential, hardness, EHOMO, electronegativity, electrophilicity, ELUMO, chemical potential, ΔEgap, softness, electronic potential, and electron affinity, were included in the computations using the Gaussian search engine, Hartree-Fock theory, and the 6-31G(d) basis set.

The outcomes of the inhibitors are shown in **Fig. 8**. Electronic energy, or Eh for short, is one of the most crucial measurements in quantum chemistry for comprehending molecule behavior. Changing Eh means that the security and bonding links between molecules are changing, while higher energy levels mean that the molecules are reacting more. Among the analyzed compounds, Bisdemethoxycurcumin had the greatest dipole moment (5.4579 Debye), signifying substantial polarity, while Epicatechin displayed the lowest dipole moment (1.3459 Debye), suggesting comparatively reduced polarity compared to the control molecule CID 5113032. The disparities in charge distribution and chemical polarity were emphasized by the variations in dipole moments. Bisdemethoxycurcumin exhibited a high degree of polarity, while Epicatechin exhibited a significantly reduced degree of polarity. The determined frontier molecular orbital (FMO) energies indicate unique electronic characteristics for each compound. The EHOMO and ELUMO values for Kaempferol are −0.305 au and 0.056 au, respectively, which leads to an energy gap (ΔEgap) of −0.361 au (**Fig. 8B**). Catechin exhibits an EHOMO of −0.307 au and an ELUMO of 0.152 au, resulting in a ΔEgap of −0.459 au (**Fig. 8C**). In the meantime, Epicatechin exhibits an EHOMO of −0.318 au and an ELUMO of 0.137 au, resulting in a ΔEgap of −0.455 au (**Fig. 8D**), and Bisdemethoxycurcumin shows an EHOMO of −0.304 au and an ELUMO of 0.060 au, resulting in an energy gap of −0.364 au. The energy gap values indicate the electronic stability and potential reactivity of the compounds. The molecular orbital results of the activators are shown in **Fig. 9** in comparison with the control molecule, Resveratrol. Carnosol had the highest dipole moment (5.0978 Debye), a higher value than the control, and Ginsenosides had the lowest dipole moment (2.1479 Debye), a lower value than the control. The EHOMO and ELUMO values are −0.244 au and 0.264 au, respectively, for Carnosol, leading to an energy gap (ΔEgap) of −0.508 au (**Fig. 9B**). Ginsenosides show an EHOMO of −0.343 au and an ELUMO value of 0.0483 au, resulting in a ΔEgap of −0.386 au (**Fig. 9C**).

**Fig. 8.**
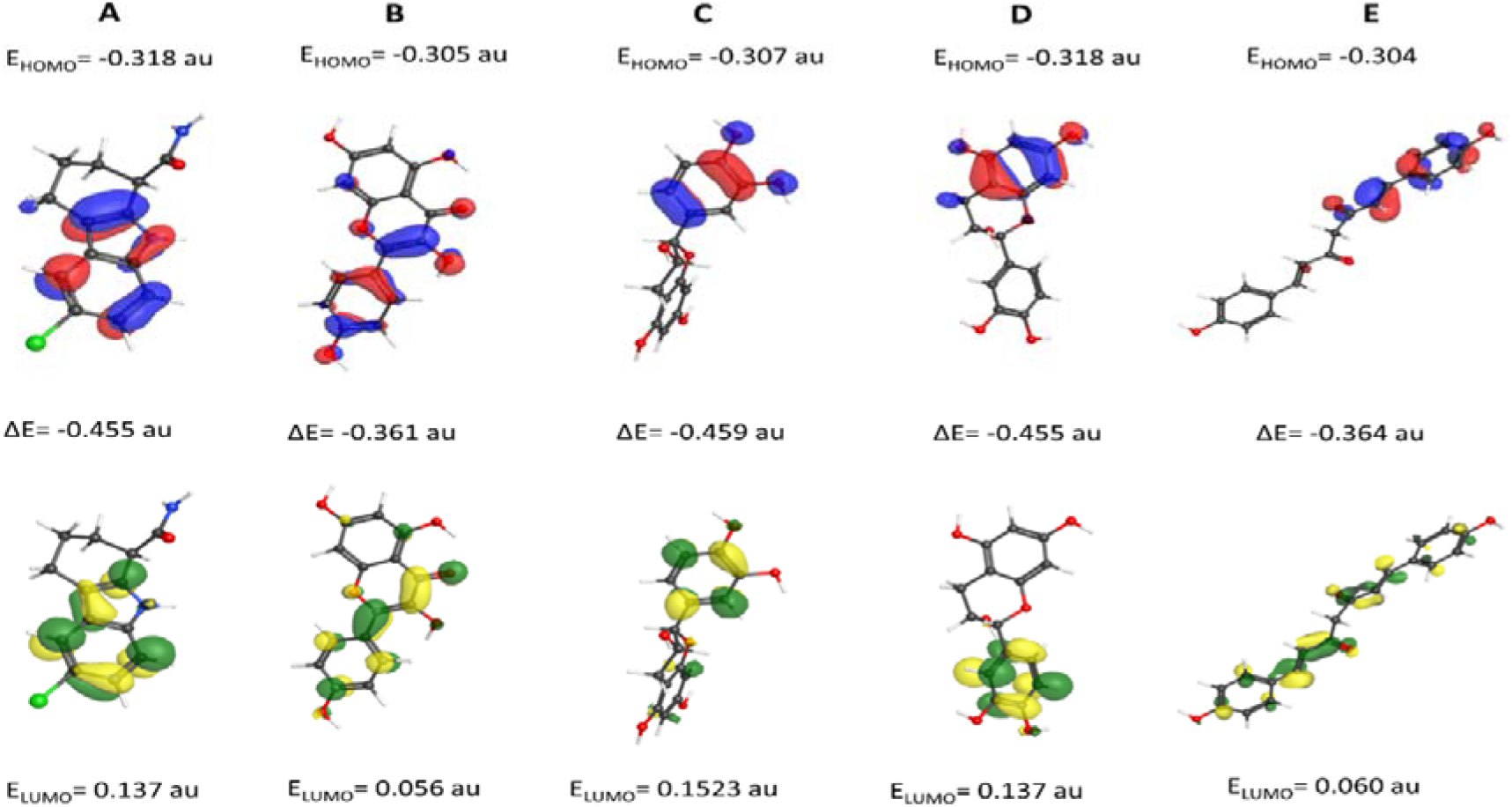
Molecular orbital distributions of inhibitors compounds A) Selisistat (Control compound), B) Kaempferol (CID 5280863), C) Catechin (CID 9064), D) Epicatechin (CID 72276), and E) Bisdemethoxycurcumin (CID 5315472).

**Fig. 9.**
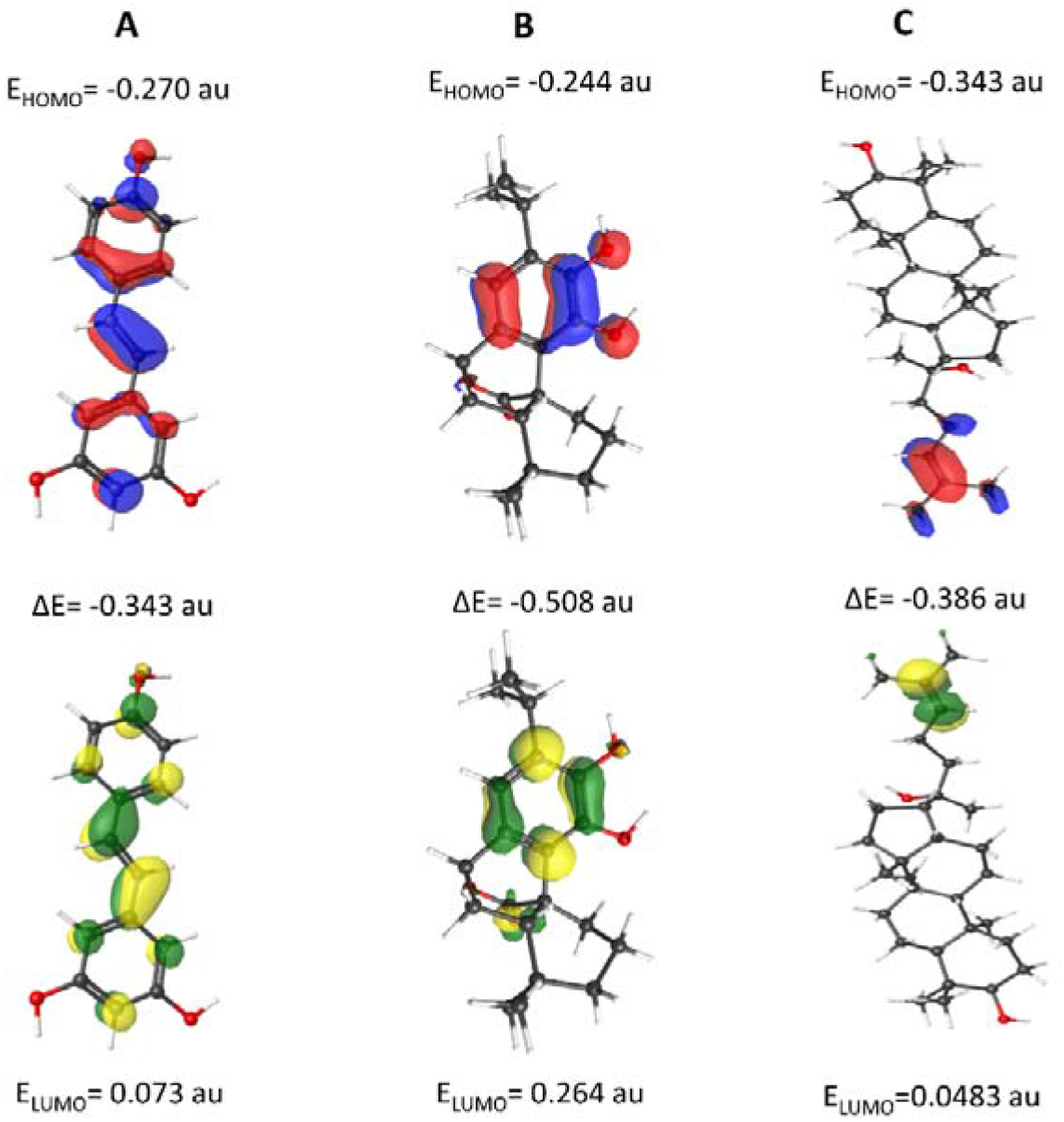
Molecular orbital distributions of activators (A) Resveratrol (Control compound), (B) Carnosol (CID 442009), and (C) Ginsenosides (CID 3086007).

### 3.6 Molecular Dynamics Analysis of Selective SIRT1-Targeting Compounds

The molecular dynamics (MD) simulation of biomolecular structures in solution provides insights into their behavior across different time intervals. Calculating thermal averages of molecular parameters enables the simulation of their time-dependent behavior, providing a close approximation to experimental ensemble observations [57]. The method assesses bulk fluid parameters and free energy changes, which play a pivotal role in phenomena like ligand binding [58]. Molecular simulations additionally examine reachable conformational states, thereby aiding processes like ligand docking [59]. The chosen complexes underwent 100 ns molecular dynamics simulations to examine their structural integrity and stability, aiming to pinpoint potential inhibitors and activators. RMSD, RMSF, Rg, and SASA analyses were carried out for the best-scoring ligand–protein complex.

#### 3.6.1 Evaluation of RMSD in Protein–Ligand Complexes

RMSD analysis was utilized to assess conformational changes and overall stability of the docked protein–ligand complex over the course of the simulation.Lower RMSD shifts values reflect a more stable atomic backbone, with reduced RMSD generally considered a favorable indicator of the complex’s conformational integrity [60]. The RMSD values of Cα atoms were analyzed to examine the structural stability of 4I5I in complex with four inhibitors (CID 5280863, CID 9064, CID 72276, and CID 5315472) relative to the control compound (CID 5113032). The RMSD mean values for the inhibitor complexes were 3.56 Å, 2.25 Å, 4.72 Å, and 4.07 Å, corresponding to each complex, whereas the control exhibited a higher average RMSD of 4.96 Å, indicating greater structural deviation. Among all compounds, CID 9064 displayed the lowest RMSD, indicating the highest structural stability. It initiated at 0.366 Å, peaked at 2.991 Å around 74.2 ns, and stabilized between 2.0–2.5 Å after the first 20 ns, maintaining a final RMSD of 2.878 Å. CID 5280863 showed a rapid increase to 4.5 Å within the first 10 ns, reached a maximum of 5.827 Å at 95.8 ns, and stabilized between 3.0–4.5 Å, concluding at 4.161 Å, indicative of moderate flexibility following an initial adjustment phase.CID 72276 started at 0.385 Å, reached a peak of 7.076 Å at 13.4 ns, and ended at 3.881 Å, suggesting a flexible yet stable conformation. CID 5315472 exhibited similar dynamics, beginning at 0.368 Å, peaking at 6.062 Å (31.8 ns), and fluctuating between 3.5–5.5 Å, ultimately stabilizing at 4.869 Å. The control compound displayed the highest deviation, with RMSD ranging between 4.5–5.5 Å after peaking at 6.191 Å (46.5 ns), and finishing at 5.601 Å. Overall, CID 9064 exhibited the most stable binding profile, characterized by the lowest average and peak RMSD values. These findings suggest that CID 9064 confers the greatest conformational stability to 4I5I among the tested inhibitors, underscoring its role as a reliable stabilizing compound.

Alternatively, the mean Cα RMSD readings for the activator complexes of CID 442009 and CID 3068007 bound to 4I5I were 5.26 Å and 4.07 Å in sequential order, whereas the control compound (CID 445154) displayed a lower overall RMSD of 3.04 Å, as shown in **Fig. 10**. Notably, CID 3068007 demonstrated less fluctuation (range: 6.500 Å–0.588 Å), closely aligning with the control complex, indicating a more stable conformation during the simulation period. In contrast, CID 442009 showed the highest average RMSD (5.26 Å) and the widest fluctuation range (10.581 Å; max: 11.132 Å, min: 0.551 Å), suggesting significant structural deviations, particularly around 39.9 ns. These findings reflect the varying degrees of structural stability among the protein–ligand complexes, with CID 3068007 exhibiting the most stable binding conformation among the activator compounds.

**Fig. 10.**
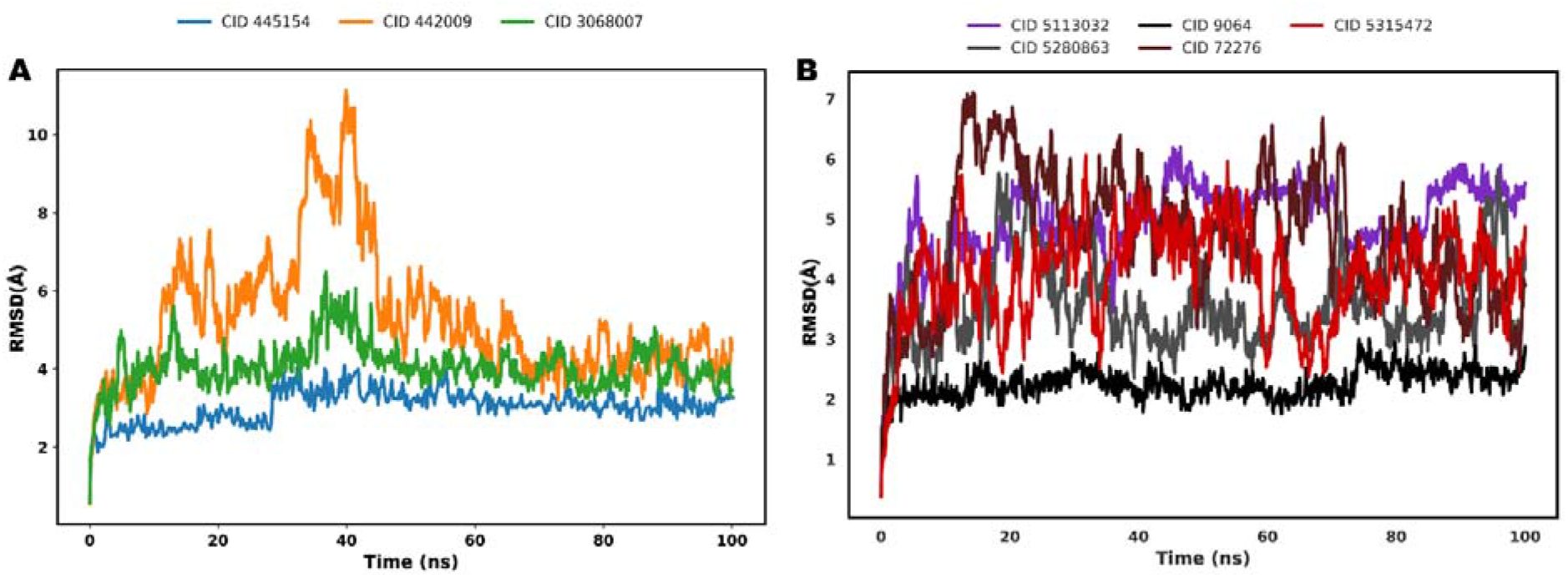
Molecular dynamics simulation analysis of the SIRT1 protein in complex with four inhibitor compounds and a control, as well as two activator compounds and a corresponding control. (A) RMSD plots depicting the stability of SIRT1 in complex with two activator compounds and the control [C0 – Resveratrol (CID 445154), C1 – Carnosol (CID 442009), and C2 – Ginsenosides (CID 3068007)]. (B) Root mean square deviation (RMSD) over a 100 ns simulation, illustrating the stability of SIRT1 bound to four distinct compounds [Selisistat (Control 5113032), Kaempferol (CMPND 5280863), Catechin (CMPND 9064), Epicatechin (CMPND 72276), and Bisdemethoxycurcumin (CMPND 5315472)]. The plots highlight structural fluctuations of the protein upon ligand binding, thereby demonstrating the influence of each compound on the conformational stability of SIRT1.

#### 3.6.2 Evaluation of RMSF in Protein–Ligand Complexes

Through RMSF analysis, the positional fluctuations of protein residues were quantified during ligand binding over the simulation trajectory. RMSF quantifies the extent to which atomic coordinates deviate from their mean values over the course of MD simulations or related dynamic analyses [61]. RMSF analysis was applied to probe the residue-specific conformational adaptability of the 4I5I protein across all complexes. Notable regions of increased fluctuation were observed at residues GLU36, LYS80, VAL263, and LYS272, which likely reflect intrinsic conformational dynamics or local rearrangements induced by ligand binding. The mean RMSF values for the protein bound to CID 5280863, CID 9064, CID 72276, and CID 5315472 amounted to 1.82 Å, 1.39 Å, 1.83 Å, and 1.53 Å, respectively. In contrast, the control compound CID 5113032 and the apo form of 4I5I exhibited higher average fluctuations of 1.74 Å and 2.93 Å, respectively. Among the tested inhibitors, CID 9064 conferred the highest conformational stability, as evidenced by the lowest average RMSF (1.39 Å), followed by CID 5315472 (1.53 Å). Both CID 5280863 and CID 72276 displayed slightly elevated fluctuations (∼1.8 Å), indicating relatively less stabilization. The significantly higher flexibility observed in the apo form underscores the stabilizing influence of ligand binding. Additionally, the control compound induced lower fluctuations than the apo form but was less effective than the inhibitors, particularly CID 9064. Collectively, the RMSF data reinforce the RMSD findings, highlighting CID 9064 as the most effective stabilizer of 4I5I by minimizing residue-level fluctuations and preserving structural integrity throughout the simulation.

By comparison, the residue-specific RMSF of the activators was calculated over a 100 ns molecular dynamics simulation. Specifically, residues ASN1, CYS13, GLN105, ALA160, LEU195, GLU227, and GLU237 exhibited significant flexibility, and these residues are known to play key roles in ligand binding, as illustrated in **Fig. 11**. Among these residues, CYS13 displayed the highest 5 8 RMSF value, reaching 14.54 Å. The mean RMSF for 4I5I bound to CID 442009, CID 3068007, and the control compound amounted to 2.95 Å, 2.34 Å, and 1.62 Å in that order. Conversely, the apo form exhibited a heightened RMSF of 3.52 Å, suggesting reduced receptor flexibility following interaction with the selected ligands and implying increased structural stability of the resulting protein–ligand complexes.

**Fig. 11.**
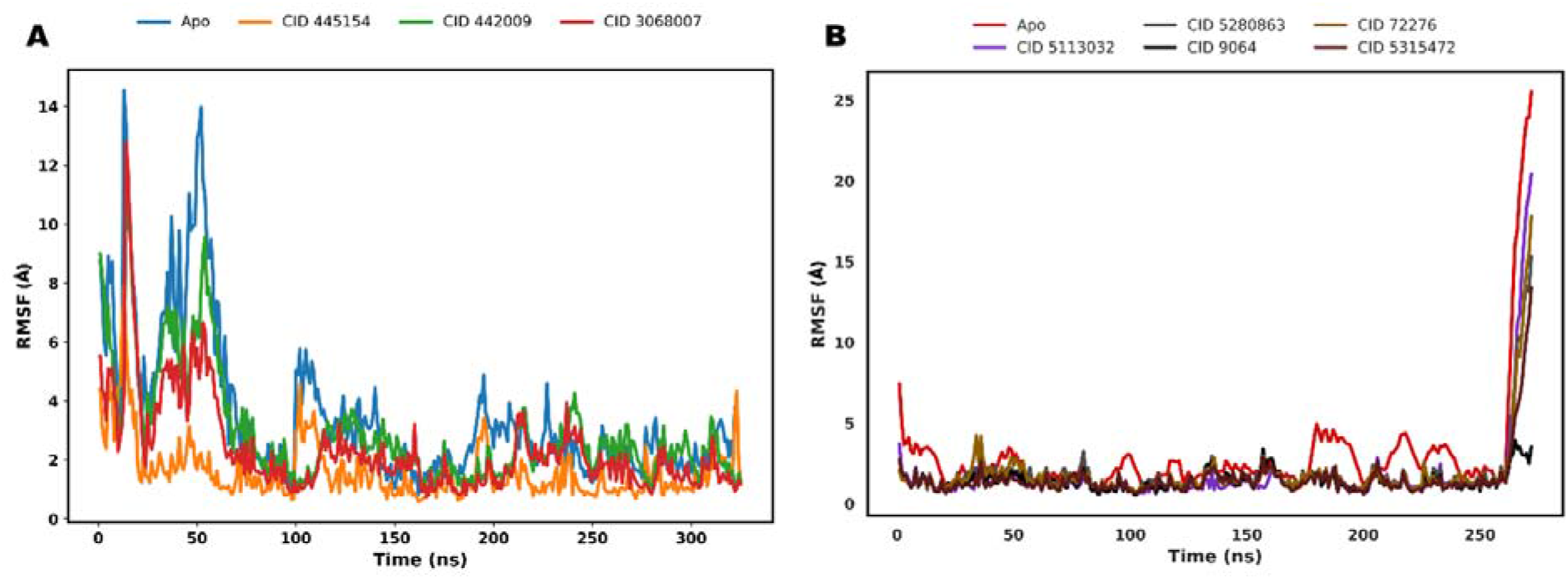
Molecular dynamics investigation of the SIRT1 protein bound to activator and inhibitor molecules, along with a control. (A) RMSF profile of SIRT1 in the presence of activators, showing residue-level flexibility and structural compactness induced by the ligands. The apo form of SIRT1 is compared with the SIRT1-compound complexes, highlighting the dynamic behavior of the protein in response to different ligands. (B) RMSF profile for the SIRT1 protein with inhibitors, illustrating the residue-specific movements and structural compactness in the presence of the compounds. The apo form of SIRT1 is compared with the SIRT1-compound complexes, highlighting the dynamic behavior of the protein in response to different ligands.

#### 3.6.3 Rg analysis of protein–ligand complexes

Radius of gyration (Rg) provides insight into the structural compactness and spatial arrangement of protein atoms within ligand-bound complexes over the course of the simulation [62]. Additionally, it provides an overview of the protein’s structural configuration by reflecting changes in compactness or expansion of the system. Reduced Rg values indicate increased structural compactness [63]. During protein unfolding, the radius of gyration exhibits temporal fluctuations. Accordingly, the stability of 4I5I in association with the phytochemical ligands CID 5280863, CID 9064, CID 72276, and control CID 5113032 (inhibitor complexes presented in **Fig. 12**, along with activator complexes CID 442009 (c1), CID 3068007 (c2), and control CID 445154 (c0) was assessed by analyzing their radius of gyration over the course of a 100 ns MD trajectory. Rg analysis was conducted to characterize the protein’s folding and global stability in complexes of 4I5I with each inhibitor. The mean Cα Rg values for the complexes with CID 5280863, CID 9064, CID 72276, CID 5315472, and the control compound CID 5113032 were 20.41 Å, 20.09 Å, 20.87 Å, 20.32 Å, and 20.20 Å, respectively. Among all ligands, CID 9064 promoted the most compact conformation of the protein, as indicated by its lowest mean Rg (20.09 Å), suggesting enhanced conformational stability. CID 5315472 (20.32 Å) and CID 5280863 (20.41 Å) also maintained compact structures with slightly higher Rg values, indicative of moderate stabilizing effects. Conversely, CID 72276 exhibited the highest Rg (20.87 Å), pointing to increased structural flexibility or a less compact conformation. The control compound CID 5113032 (20.20 Å) yielded an intermediate Rg value, offering less stability than CID 9064 but more than CID 72276. Overall, the narrow range of Rg values (20.09–20.87 Å) indicates that the global fold of the 4I5I protein remains largely preserved across all complexes. However, the subtle variations in compactness corroborate the RMSD and RMSF results, confirming CID 9064 as the most effective ligand in preserving the protein’s conformational stability and compact structure of 4I5I.

**Fig. 12.**
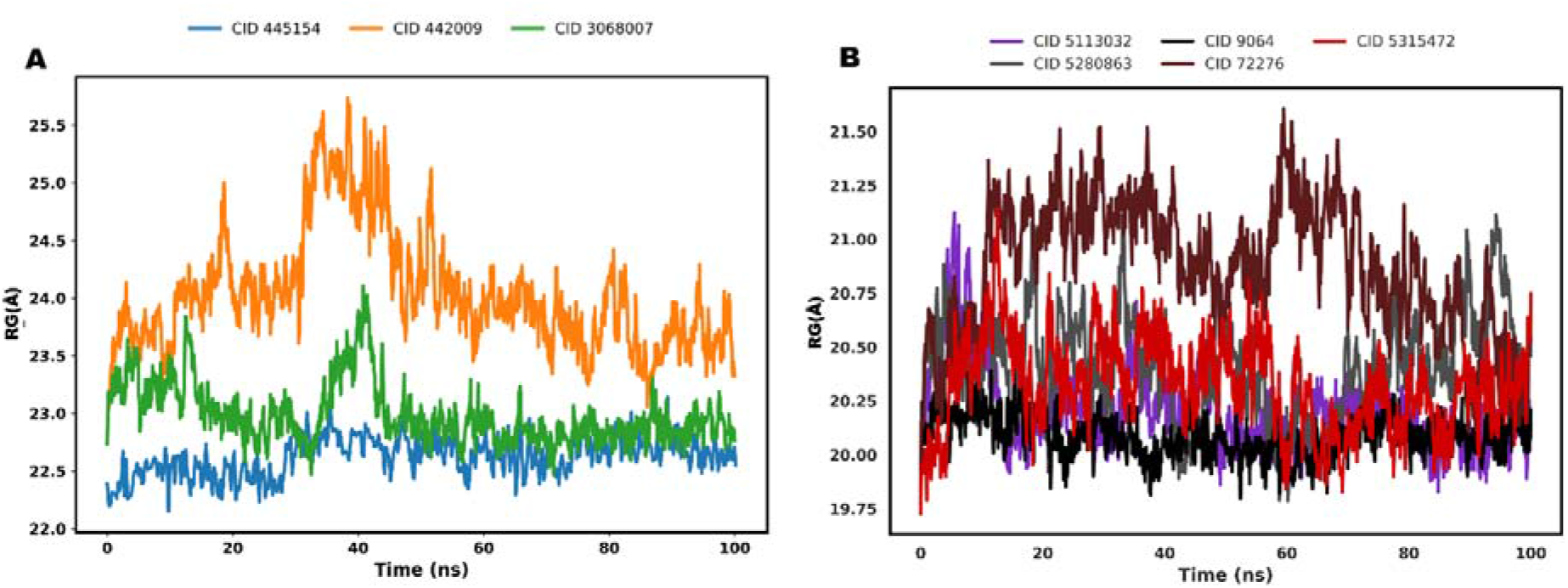
Molecular dynamics investigation of the SIRT1 protein bound to activator and inhibitor molecules, including a control. (A) Rg profile of SIRT1 depicting the structural compactness of the protein over the course of the simulation. The investigation illustrates the impact of each activator [C0 – Resveratrol (CID 445154), C1 – Carnosol (CID 442009), and C2 – Ginsenosides (CID 3086007)] and the control on the structural stability of SIRT1, as indicated by Rg measurements. (B) Rg profile of SIRT1 depicting the protein’s structural compactness over the course of the molecular dynamics simulation. Similarly, the study highlights how each inhibitor [Selisistat (Control5113032), Kaempferol (CMPND5280863), Catechin (CMPND9064), Epicatechin (CMPND72276), and Bisdemethoxycurcumin (CMPND5315472)] and the control influence SIRT1 stability, as reflected in the Rg values.

The mean radius of gyration (Rg) for 4I5I bound to the activator compounds CID 442009, CID 3068007, and the control compound CID 445154 was 24.04 Å, 22.99 Å, and 22.62 Å in that order, suggesting minimal structural rearrangements within the protein’s active site upon ligand association. Notably, the complex with CID 442009 exhibited minor fluctuations between 32 ns to 45 ns, peaking at 25.74 Å before stabilizing within the range of 23.5 24 Å. In contrast, the CID 3068007 complex demonstrated a more stable Rg profile, with a peak value of 24.11 Å observed at 40.8 ns.

#### 3.6.4 SASA analysis of protein–ligand complexes

Solvent-accessible surface area (SASA) evaluates the regions of a protein surface available for interaction with solvent molecules [64]. This study analyzed the spatial arrangement of polar and nonpolar residues within the protein, where elevated SASA values reflect increased exposure of the protein surface. We analyzed the SASA of the 4I5I protein bound to inhibitors CID 5280863, CID 9064, CID 72276, control CID 5113032 represented in **Fig. 13B** activators CID 442009 (c1), CID 3068007 (c2), and control CID 445154 (c0) (**Fig. 13A**) over a 100 ns simulation. The SASA profiles varied with each ligand, highlighting their distinct effects on protein surface exposure and stability. SASA analysis was performed to assess the degree of solvent accessibility and conformational compactness of the 4I5I protein upon ligand binding. Mean SASA values for the 4I5I complexes with CID 5280863, CID 9064, CID 72276, and CID 5315472 were 14,185.68 Å², 13,772.45 Å², 14,434.79 Å², and 13,761.46 Å², respectively. The control compound CID 5113032 exhibited an average SASA of 14,063.58 Å². Among the tested ligands, CID 9064 and CID 5315472 displayed the lowest SASA values (13,772.45 Å² and 13,761.46 Å², respectively), indicating enhanced structural compactness and reduced solvent exposure—features characteristic of stable protein–ligand interactions. Notably, CID 9064 fluctuated within a narrow range (13,154.42–14,578.94 Å²), while CID 5315472 ranged from 13,131.02 to 14,468.92 Å², both reflecting consistent solvents shielding during the simulation. In contrast, CID 72276 exhibited the highest SASA (14,434.79 Å²), with substantial fluctuation (13,805.70–15,195.41 Å²), suggesting increased flexibility or partial unfolding of the protein–ligand complex, possibly due to weaker binding affinity. CID 5280863 showed intermediate behavior with moderate variation (13,269.04–14,718.79 Å²), implying occasional structural adjustments. The control compound CID 5113032 also demonstrated moderate solvent exposure, with values ranging from 13,233.42 to 14,749.26 Å², serving as a comparative benchmark. Overall, the SASA results align with the RMSD, RMSF, and Rg analyses, further supporting CID 9064 and CID 5315472 as the most effective stabilizers of 4I5I, likely due to their strong and consistent binding that minimizes solvent accessibility.

**Fig. 13.**
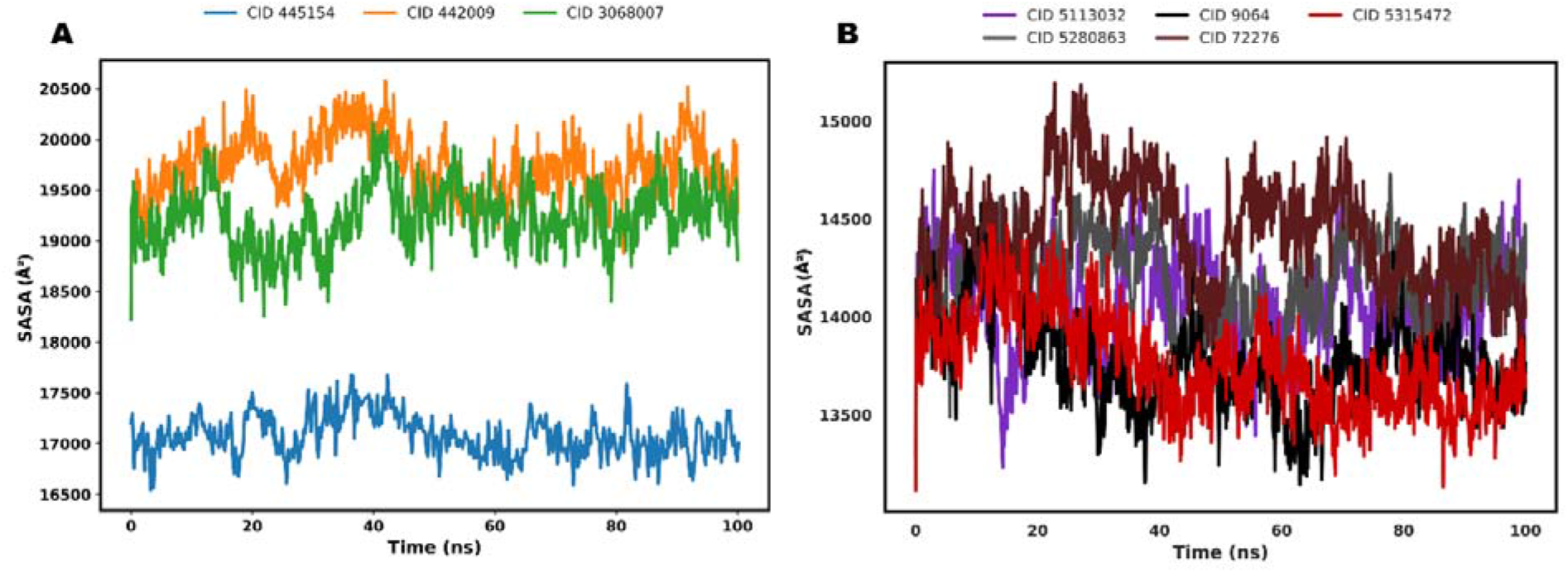
Molecular dynamics investigation of the SIRT1 protein bound to activator and inhibitor molecules, including a control. (A) SASA profile depicting the temporal variation in protein surface exposure of SIRT1 during a 100 ns molecular dynamics simulation with two activators and a control compound [C0 – Resveratrol (CID 445154), C1 – Carnosol (CID 442009), and C2 – Ginsenosides (CID 3086007)]. The observed fluctuations in SASA indicate alterations in protein surface accessibility and compactness upon ligand association. (B) SASA profile illustrating changes in SIRT1 surface exposure over a 100 ns MD trajectory in complexes with four inhibitors and a control compound [Selisistat (Control5113032), Kaempferol (CMPND5280863), Catechin (CMPND9064), Epicatechin (CMPND72276), and Bisdemethoxycurcumin (CMPND5315472)]. Variations in SASA reflect adjustments in protein surface accessibility and structural compactness induced by ligand binding.

The mean SASA values for the 4I5I protein complexes over a 100 ns simulation were 19,733.1 Å² for CID 442009 (c1), 19,229.0 Å² for CID 3068007 (c2), and 17,077.9 Å² for the control compound (c0). Both ligand-bound complexes exhibited no significant conformational changes and maintained structural stability after approximately 45 ns throughout the remainder of the 100 ns simulation.

### 3.7 Free Energy Landscape Analysis

The multifaceted links between conformational states, structural stability, and thermodynamic favorability are depicted in detail by the Free Energy Landscape (FEL) [65]. The dynamic behavior of molecular complexes, transition routes, and lowest energy states may all be found by examining the FEL. **Fig. 14 and Fig. 15** show that compact and structurally stable structures are found in regions on the FEL with reduced Radius of Gyration (RG) and Root-Mean-Square Deviation (RMSD) values. These regions reflect thermodynamically advantageous states when combined with lower Gibbs Free Energy (GFE). By breaking down the overall free energy into its component parts, solvation energy and molecular mechanics energy, the Gibbs Free Energy (GFE) of a protein-ligand complex is determined. Additionally, structural changes within the complex are indicated by regions exhibiting higher root mean square deviation (RMSD) values, which are typically associated with less stable configurations. Elevated GFE in these areas reflects the presence of higher energy barriers and thermodynamically unfavorable states.

**Fig. 14.**
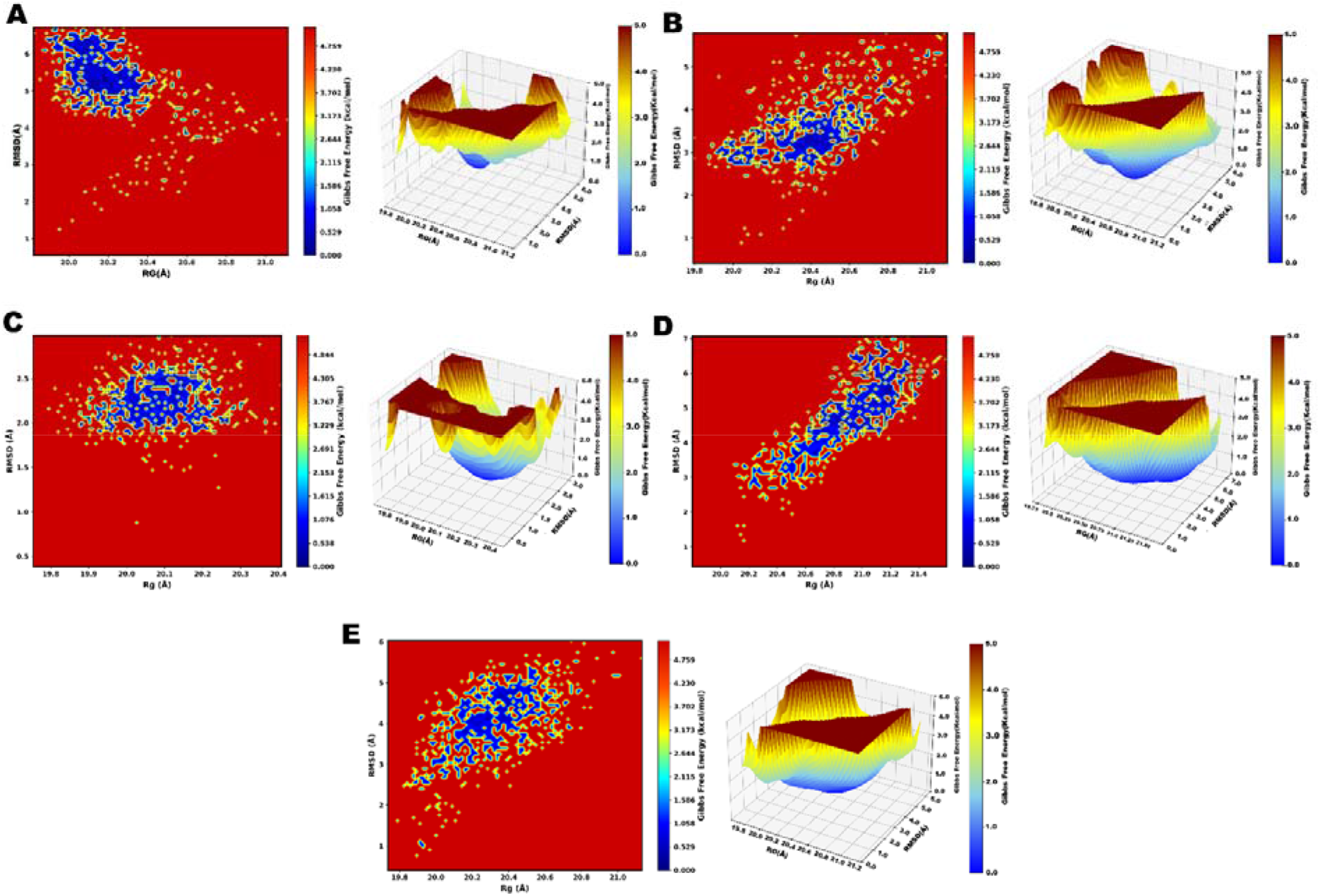
Free Energy Landscape (FEL) analysis of SIRT1 complexes with kaempferol, catechin, epicatechin, and bisdemethoxycurcumin. (A) Represents the FEL of the SIRT1-kaempferol (5280863) complex. The 3D plot illustrates the global free energy distribution as a function of Root Mean Square Deviation (RMSD) and Radius of Gyration (Rg), highlighting the conformational stability of the system. The 2D contour map further visualizes the free energy basins, where the blue regions indicate the most stable low-energy conformations, and the red regions denote high-energy states corresponding to less stable configurations. (B) Depicts the FEL of the SIRT1-catechin (9064) complex. Similar to kaempferol, the 3D plot shows the distribution of free energy across RMSD and Rg values, while the 2D contour map identifies stable (blue) and unstable (red) conformational states, indicating the dynamic behavior of the complex during the simulation. (C) FEL of the SIRT1-epicatechin (72276) complex. The 3D plot shows the global free energy distribution as a function of Root Mean Square Deviation (RMSD) and Radius of Gyration (Rg), illustrating the conformational stability of the system. The 2D contour map highlights stable (blue) and unstable (red) regions, with the blue regions representing low-energy, highly stable conformations. (D) FEL of the SIRT1-bisdemethoxycurcumin (5315472) complex. The 3D plot visualizes the free energy changes based on RMSD and Rg, identifying stable conformational states. The corresponding 2D contour map further emphasizes the energy basins, where blue regions denote stable conformations and red regions indicate higher-energy, less stable states. (E) FEL of the protein-control complex.

**Fig. 15.**
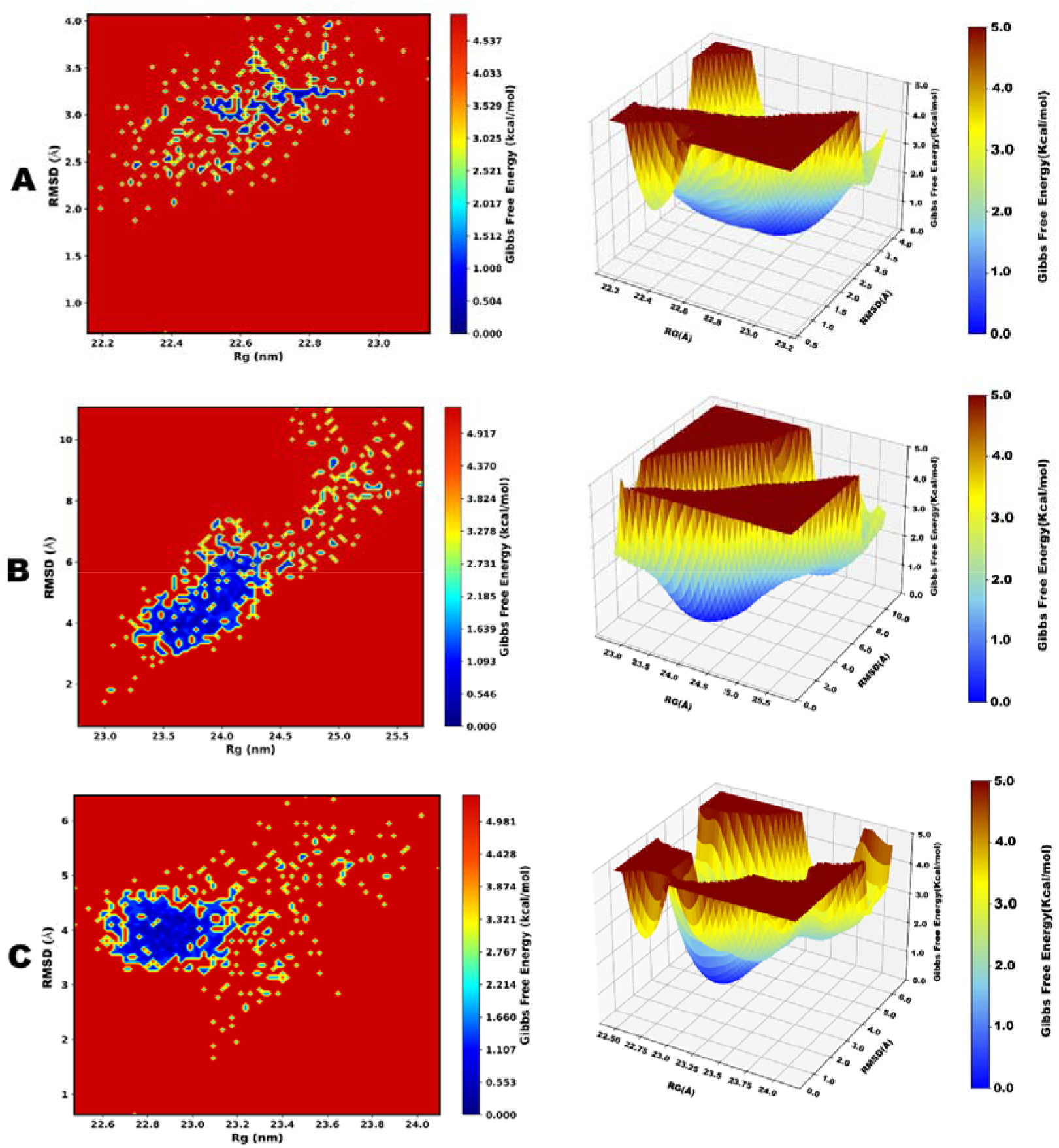
Free Energy Landscape (FEL) analysis of SIRT1 complexes with Resveratrol, Carnosol, and Ginsenosides. (A) Represents the FEL of the SIRT1-Resveratrol (445154) complex. The 3D plot illustrates the global free energy distribution as a function of Root Mean Square Deviation (RMSD) and Radius of Gyration (Rg), highlighting the conformational stability of the system. The 2D contour map further visualizes the free energy basins, where the blue regions indicate the most stable low-energy conformations, and the red regions denote high-energy states corresponding to less stable configurations. (B) Depicts the FEL of the SIRT1-Carnosol (442009) complex. Similar to kaempferol, the 3D plot shows the distribution of free energy across RMSD and Rg values, while the 2D contour map identifies stable (blue) and unstable (red) conformational states, indicating the dynamic behavior of the complex during the simulation. (C) FEL of the SIRT1-Ginsenosides (3086007) complex. The 3D plot shows the global free energy distribution as a function of Root Mean Square Deviation (RMSD) and Radius of Gyration (Rg), illustrating the conformational stability of the system. The 2D contour map highlights stable (blue) and unstable (red) regions, with the blue regions representing low-energy, highly stable conformations.

Potential structural changes or dynamic shifts are signaled by transition states, which are characterized by an increase in both Root Mean Square Deviation (RMSD) and Radius of Gyration (RG) values. The energy barriers associated with transitions between these states are reflected in the steepness of the Free Energy Landscape (FEL) surface, providing valuable insight into the challenges associated with conformational changes.

### 3.8 Probability density function

The Probability Density Function (PDF) analysis was conducted to explore the relationship between Radius of Gyration (RG) and Root-Mean-Square Deviation (RMSD) for four molecular systems [52,66] labeled as A, B, C, and D. These investigations offer important new information about the stability, flexibility, and structural dynamics of the corresponding molecular systems.

The RMSD values of kaempferol in **Fig. 16A** range from 1.0 Å to 6.0 Å, while the RG values range from 19.8 Å to 21.2 Å. The most stable structural condition is represented by the high-density area (in red) near RG ∼20.5 Å and RMSD ∼3.0 Å. There is a significant connection between RG and RMSD, as evidenced by the extended density distribution, which indicates that the system exhibits considerable structural flexibility and variability. The range of radius of gyration (RG) values (19.8 Å to 20.4 Å) and root mean square deviation (RMSD) values (0.5 Å to 3.0 Å) is more restricted in **Fig. 16B**. At approximately RG ∼20.1 Å and RMSD ∼2.0 Å for the catechin, there is a dense concentration that indicates a more compact and stable structural state. The tighter contour lines suggest limited structural flexibility, highlighting a stronger correlation between RG and RMSD.

**Fig. 16.**
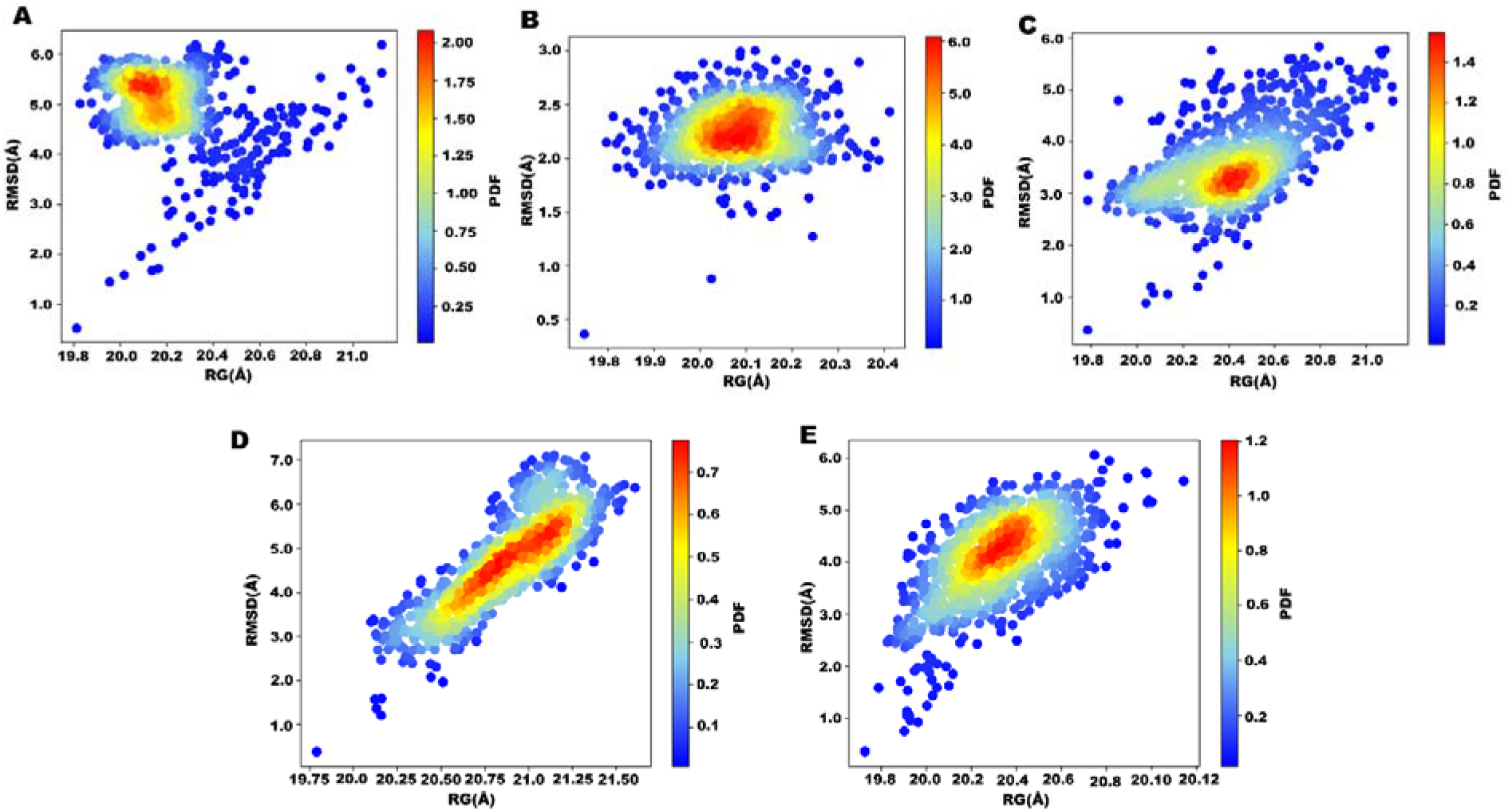
Probability Density Function (PDF) analysis of SIRT1 complexes with kaempferol, catechin, epicatechin, and bisdemethoxycurcumin. (A) PDF analysis of the SIRT1-kaempferol (5280863) complex. The plot shows the probability distribution of conformations based on Root Mean Square Deviation (RMSD) and Radius of Gyration (Rg). Higher probability regions are represented in red, indicating the most frequently sampled conformational states during the simulation, while blue regions correspond to less frequent states. (B) PDF analysis of the SIRT1-catechin (9064) complex. The distribution highlights stable conformational states, with red areas denoting higher probability density, reflecting the preferred conformations of the complex. The blue regions indicate less populated conformational states. (C) PDF analysis of the SIRT1-epicatechin (72276) complex. The plot illustrates the probability distribution of conformational states based on Root Mean Square Deviation (RMSD) and Radius of Gyration (Rg). The red regions represent the most frequently sampled stable conformations, while blue regions indicate less frequent states. (D) PDF analysis of the SIRT1-bisdemethoxycurcumin (5315472) complex. The distribution highlights the preferred conformational states of the complex, with red areas showing higher probability density and blue areas reflecting less populated states. (E) PDF analysis of the protein-control complex, Selisistat.

The PDF of the epicatechin complex, with RG values ranging from 19.75 Å to 21.5 Å and RMSD values between 3.0 Å and 7.0 Å, is presented in **Fig. 16C**. A distinct high-density area is observed at RG approximately 20.75 Å and RMSD around 5.0 Å, indicating the most likely structural arrangement. The bisdemethoxycurcumin complex shows the RG value ranging from 19.8 Å to 21.2 Å and RMSD values between 2 Å and 6 Å, as presented in **Fig. 16D**. The densest area of this heatmap remains within the RG value 20.3 Å to 20.6 Å, indicating structural arrangement and compactness of the complex. **Fig. 16E** shows the result of the protein-control complex, showing a similar RG value range of the kaempferol and a dense area of structural arrangement remains within 4.1 Å to 5.5 Å. The diagonal elongation of the density distribution suggests a positive correlation between RG and RMSD, implying that there is substantial structural variability within the system.

The structural dynamics, flexibility, and stability of the activator-protein complex are represented in **Fig. 17** through the relationship between Radius of Gyration (RG) and Root-Mean-Square Deviation (RMSD) for two compounds and a control. The PDF of the control compound is represented in **Fig. 17A**; the range of radius of gyration (RG) values (22.2 Å to 23.2 Å) and root mean square deviation (RMSD) values (1.8 Å to 3.9 Å). Carnosol (**Fig. 17B**) shows a compact and stable complex structure around 23.8 Å and an RG value of 4.1 Å of RMSD value. The range of the RG value for the compound is 23.0 Å to 25.5 Å, and for the RMSD value, it remains around 2.3 Å to 5.9 Å. Another activator, Ginsenosides, is represented within **Fig. 17C** with the RG value of 22.50 Å to 24.0 Å, and a 3.2 Å to 4.8 Å RMSD range indicates the stability of the complex. A compact and stable region of the complex remains within the 22.85 Å RG value and 3.9 Å RMSD value.

**Fig. 17.**
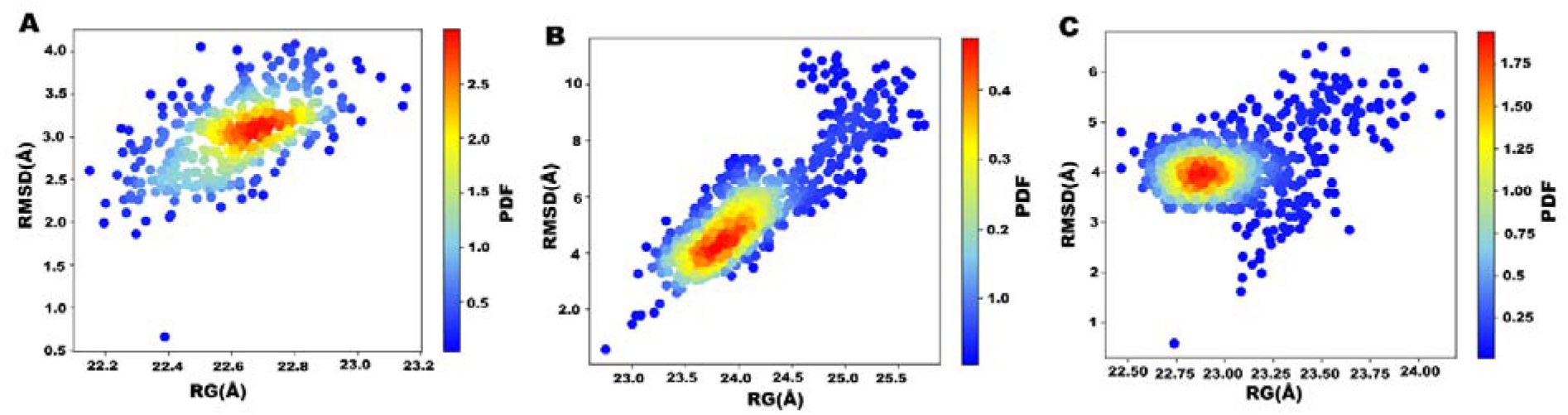
Probability Density Function (PDF) analysis of SIRT1 complexes with Resveratrol, Carnosol, and Ginsenosides. (A) Represents the PDF for the control compound, Resveratrol (CID 445154). (B) The PDF plot analysis for another activator compound, Carnosol (CID 442009). The plot shows the probability distribution of conformations based on Root Mean Square Deviation (RMSD) and Radius of Gyration (Rg). Higher probability regions are represented in red, indicating the most frequently sampled conformational states during the simulation, while blue regions correspond to less frequent states. (C) Represents the PDF analysis of the Ginsenosides (CID 3086007). The plot shows the probability distribution of conformations based on Root Mean Square Deviation (RMSD) and Radius of Gyration (Rg). Higher probability regions are represented in red, indicating the most frequently sampled conformational states during the simulation, while blue regions correspond to less frequent states.

## 4. Conclusions

This study provides an integrated experimental and computational framework for the discovery of novel SIRT1 modulators with potential therapeutic applications. By combining protein expression, purification, and binding validation with high-throughput docking, molecular dynamics simulations, and free energy analyses, we identified Catechin, Epicatechin, Kaempferol, and Bisdemethoxycurcumin as promising inhibitors, while Ginsenosides and Carnosol emerged as strong activators. Among these, Catechin and Ginsenosides demonstrated the most favorable stability, binding energetics, and drug-likeness, supported by robust ADMET and toxicity profiles. Furthermore, the mechanistic insights into the SIRT1–METTL3 regulatory axis underscore the importance of post-translational modifications in shaping oncogenic pathways and provide a rationale for targeting this interaction to suppress metastasis.

Collectively, our findings highlight phytochemicals as viable scaffolds for developing selective SIRT1 modulators capable of influencing cancer progression and metabolic disorders. Future work should include enzymatic assays and in vivo studies to validate their biological efficacy and safety. The therapeutic potential demonstrated here strengthens the case for natural compounds as cost-effective and accessible candidates in precision medicine approaches targeting SIRT1.

## CRediT author statement

**Md Kazy Ebnul Hasan:** Writing – review & editing, Writing – original draft, Visualization, Software, Resources, Project administration, Formal analysis, Data curation, Conceptualization. **Md. Nahid Hasan Joy:** Writing – review & editing, Writing – original draft, Visualization, Validation, Software, Resources, Methodology, Investigation, Formal analysis, Data curation. **Liqing Lin:** Writing – review & editing, Writing – original draft, Visualization, Validation, Software, Methodology, Supervision. **Md. Sanowar Hossan:** Writing – review & editing, Investigation, Formal analysis, Software, Resources. **Shahad Shahriar:** Writing – review & editing, Validation, Software, Resources, Formal analysis. **Md. Mahedi Hasan Sourov:** Writing – review & editing, Software, Formal analysis, Visualization, Validation. **Md. Naymur Rahman:** Writing – review & editing, Visualization, Validation. **Md. Faruk Hasan:** Writing – review & editing, Visualization,

Software, Resources, Investigation, Formal analysis, Data curation. **Md. Enamul Haque:** Writing – review & editing, Visualization, Validation, Software, Resources**. Abdullah-Al-Jubayer:** Writing – review & editing, Visualization, Validation, Software, Resources. **Mamudul Hasan Razu:** Writing – review & editing, Supervision, Validation, Formal analysis. **Md. Samiul Islam:** Writing – review & editing, Formal analysis, Validation. **Mengqi Lv:** Writing – review & editing, Writing – original draft, Validation, Supervision, Formal analysis, Data curation. **Yunyu Shi:** Writing – review & editing, Writing – original draft, Validation, Supervision, Formal analysis, Data curation.

## Availability of data and materials

Structural data used in this study are publicly available from the Protein Data Bank (accession codes 4I5I and 5BTR) and from UniProt (Q96EB6). Chemical structures were retrieved from PubChem under the CIDs cited in the text and in Table S1. Isothermal titration calorimetry raw traces, docking poses and molecular dynamics trajectories generated in this study are available from the corresponding author on reasonable request.

## Ethics statement

This manuscript does not include any data obtained from human subjects, animals, or third parties without proper authorization or consent. In our manuscript, we used publicly available data with proper reference and followed the authorized guidelines in case of data retrieval.

## Acknowledgements

This research was supported by the National Natural Science Foundation of China (32470817, 32090040, and 32293213), the Strategic Priority Research Program of the Chinese Academy of Sciences (XDB37010105 and XDB39010300), and the Fundamental Research Funds for the Central Universities (WK9100000067).

## Declaration of generative AI and AI-assisted technologies in the writing process

During the preparation of this work the authors did not use generative artificial intelligence or AI-assisted technologies to produce scientific content, figures, data or analyses. Where such tools were used, their use was limited to language editing and readability, and the authors reviewed and edited the content as needed and take full responsibility for the content of the published article

## References

[1] Z. Lin, D. Fang, The Roles of SIRT1 in Cancer, Genes Cancer 4 (2013). 10.1177/1947601912475079.

[2] Q.J. Wu, T.N. Zhang, H.H. Chen, X.F. Yu, J. Le Lv, Y.Y. Liu, Y.S. Liu, G. Zheng, J.Q. Zhao, Y.F. Wei, J.Y. Guo, F.H. Liu, Q. Chang, Y.X. Zhang, C.G. Liu, Y.H. Zhao, The sirtuin family in health and disease, Signal Transduct. Target. Ther. 7 (2022). 10.1038/s41392-022-01257-8.

[3] P. Shen, X. Deng, Z. Chen, X. Ba, K. Qin, Y. Huang, Y. Huang, T. Li, J. Yan, S. Tu, SIRT1: A Potential Therapeutic Target in Autoimmune Diseases, Front. Immunol. 12 (2021). 10.3389/fimmu.2021.779177.

[4] H. Dai, A.W. Case, T. V. Riera, T. Considine, J.E. Lee, Y. Hamuro, H. Zhao, Y. Jiang, S.M. Sweitzer, B. Pietrak, B. Schwartz, C.A. Blum, J.S. Disch, R. Caldwell, B. Szczepankiewicz, C. Oalmann, P.Y. Ng, B.H. White, R. Casaubon, R. Narayan, K. Koppetsch, F. Bourbonais, B. Wu, J. Wang, D. Qian, F. Jiang, C. Mao, M. Wang, E. Hu, J.C. Wu, R.B. Perni, G.P. Vlasuk, J.L. Ellis, Crystallographic structure of a small molecule SIRT1 activator-enzyme complex, Nat. Commun. 6 (2015). 10.1038/ncomms8645.

[5] X. Zhao, D. Allison, B. Condon, F. Zhang, T. Gheyi, A. Zhang, S. Ashok, M. Russell, I. MacEwan, Y. Qian, J.A. Jamison, J.G. Luz, Correction: The 2.5 Å Crystal Structure of the SIRT1 Catalytic Domain Bound to Nicotinamide Adenine Dinucleotide (NAD+) and an Indole (EX527 Analogue) Reveals a Novel Mechanism of Histone Deacetylase Inhibition (Journal of Medicinal Chemistry (2013) 56 (963-969) DOI: 10.1021/jm301431y), J. Med. Chem. 59 (2016). 10.1021/acs.jmedchem.6b00130.

[6] S. Huang, Y. Li, G. Sheng, Q. Meng, Q. Hu, X. Gao, Z. Shang, Q. Lv, Sirtuin 1 promotes autophagy and proliferation of endometrial cancer cells by reducing acetylation level of LC3, Cell Biol. Int. 45 (2021). 10.1002/cbin.11549.

[7] B. Jablonska, M. Gierdalski, L.J. Chew, T. Hawley, M. Catron, A. Lichauco, J. Cabrera-Luque, T. Yuen, D. Rowitch, V. Gallo, Sirt1 regulates glial progenitor proliferation and regeneration in white matter after neonatal brain injury, Nat. Commun. 7 (2016). 10.1038/ncomms13866.

[8] F. Imperatore, J. Maurizio, S. Vargas Aguilar, C.J. Busch, J. Favret, E. Kowenz Leutz, W. Cathou, R. Gentek, P. Perrin, A. Leutz, C. Berruyer, M.H. Sieweke, SIRT1 regulates macrophage self renewal, EMBO J. 36 (2017). 10.15252/embj.201695737.

[9] H. Vaziri, D. Sk, N.E. E, I. S, F. RA, P. TK, L. Guarente, W. RA, hSIR2(SIRT1) functions as an NAD-dependent p53 deacetylase, Cell %19;107 (2001).

[10] B. Heltweg, T. Gatbonton, A.D. Schuler, J. Posakony, H. Li, S. Goehle, R. Kollipara, R.A. DePinho, Y. Gu, J.A. Simon, A. Bedalov, Antitumor activity of a small-molecule inhibitor of human silent information regulator 2 enzymes, Cancer Res. 66 (2006). 10.1158/0008-5472.CAN-05-3617.

[11] S. Lain, J.J. Hollick, J. Campbell, O.D. Staples, M. Higgins, M. Aoubala, A. McCarthy, V. Appleyard, K.E. Murray, L. Baker, A. Thompson, J. Mathers, S.J. Holland, M.J.R. Stark, G. Pass, J. Woods, D.P. Lane, N.J. Westwood, Discovery, In Vivo Activity, and Mechanism of Action of a Small-Molecule p53 Activator, Cancer Cell 13 (2008). 10.1016/j.ccr.2008.03.004.

[12] O.R. Bereshchenko, W. Gu, R. Dalla-Favera, Acetylation inactivates the transcriptional repressor BCL6, Nat. Genet. 32 (2002). 10.1038/ng1018.

[13] S. Mazumder, D. Plesca, M. Kinter, A. Almasan, Interaction of a Cyclin E Fragment with Ku70 Regulates Bax-Mediated Apoptosis, Mol. Cell. Biol. 27 (2007). 10.1128/mcb.01448-06.

[14] Y.H. Lin, J. Yuan, H. Pei, T. Liu, D.K. Ann, Z. Lou, KAP1 deacetylation by SIRT1 promotes non-homologous end-joining repair, PLoS One 10 (2015). 10.1371/journal.pone.0123935.

[15] Y. Wang, J. He, M. Liao, M. Hu, W. Li, H. Ouyang, X. Wang, T. Ye, Y. Zhang, L. Ouyang, An overview of Sirtuins as potential therapeutic target: Structure, function and modulators, Eur. J. Med. Chem. 161 (2019). 10.1016/j.ejmech.2018.10.028.

[16] K.T. Howitz, K.J. Bitterman, H.Y. Cohen, D.W. Lamming, S. Lavu, J.G. Wood, R.E. Zipkin, P. Chung, A. Kisielewski, L.L. Zhang, B. Scherer, D.A. Sinclair, Small molecule activators of sirtuins extend Saccharomyces cerevisiae lifespan, Nature 425 (2003). 10.1038/nature01960.

[17] J.A. Park, S. Park, W.Y. Park, M.K. Han, Y. Lee, Splitomicin, a SIRT1 inhibitor, enhances hematopoietic differentiation of mouse embryonic stem cells, Int. J. Stem Cells 12 (2019). 10.15283/ijsc18040.

[18] E.S. Hwang, S.B. Song, Nicotinamide is an inhibitor of SIRT1 in vitro, but can be a stimulator in cells, Cellular and Molecular Life Sciences 74 (2017). 10.1007/s00018-017-2527-8.

[19] Y. Li, X. He, X. Lu, Z. Gong, Q. Li, L. Zhang, R. Yang, C. Wu, J. Huang, J. Ding, Y. He, W. Liu, C. Chen, B. Cao, D. Zhou, Y. Shi, J. Chen, C. Wang, S. Zhang, J. Zhang, J. Ye, H. You, METTL3 acetylation impedes cancer metastasis via fine-tuning its nuclear and cytosolic functions, Nat. Commun. 13 (2022). 10.1038/s41467-022-34209-5.

[20] X. Jin, Y. Wei, F. Xu, M. Zhao, K. Dai, R. Shen, S. Yang, N. Zhang, SIRT1 promotes formation of breast cancer through modulating Akt activity, J. Cancer 9 (2018). 10.7150/jca.24275.

[21] E. Wang, Y. Li, H. Li, Y. Liu, R. Ming, J. Wei, P. Du, X. Li, S. Zong, H. Xiao, METTL3 Reduces Oxidative Stress-induced Apoptosis in Presbycusis by Regulating the N6-methyladenosine Level of SIRT1 mRNA, Neuroscience 521 (2023). 10.1016/j.neuroscience.2023.03.001.

[22] W. Kaplan, T.G. Littlejohn, Swiss-PDB viewer (deep view), Brief. Bioinform. 2 (2001) 195– 197.

[23] S.K. Kondapuram, S. Sarvagalla, M.S. Coumar, Docking-based virtual screening using PyRx Tool: autophagy target Vps34 as a case study, in: Molecular Docking for Computer-Aided Drug Design, Elsevier, 2021: pp. 463–477.

[24] O. Trott, A.J. Olson, AutoDock Vina: Improving the speed and accuracy of docking with a new scoring function, efficient optimization, and multithreading, J. Comput. Chem. 31 (2010) 455–461. 10.1002/jcc.21334.

[25] A.H. Arshia, S. Shadravan, A. Solhjoo, A. Sakhteman, A. Sami, De novo design of novel protease inhibitor candidates in the treatment of SARS-CoV-2 using deep learning, docking, and molecular dynamic simulations, Comput. Biol. Med. 139 (2021) 104967.

[26] S. Murugesan, S. Kottekad, I. Crasta, S. Sreevathsan, D. Usharani, M.K. Perumal, S.N. Mudliar, Targeting COVID-19 (SARS-CoV-2) main protease through active phytocompounds of ayurvedic medicinal plants – Emblica officinalis (Amla), Phyllanthus niruri Linn. (Bhumi Amla) and Tinospora cordifolia (Giloy) – A molecular docking and simulation study, Comput. Biol. Med. 136 (2021). 10.1016/j.compbiomed.2021.104683.

[27] W.F. Polik, J.R. Schmidt, WebMO: Web-based computational chemistry calculations in education and research, Wiley Interdiscip. Rev. Comput. Mol. Sci. 12 (2022). 10.1002/WCMS.1554.

[28] C.R. Legler, N.R. Brown, R.A. Dunbar, M.D. Harness, K. Nguyen, O. Oyewole, W.B. Collier, Scaled quantum mechanical scale factors for vibrational calculations using alternate polarized and augmented basis sets with the B3LYP density functional calculation model, Spectrochim. Acta A Mol. Biomol. Spectrosc. 145 (2015). 10.1016/j.saa.2015.02.103.

[29] R.G. Parr, L. V. Szentpály, S. Liu, Electrophilicity index, J. Am. Chem. Soc. 121 (1999). 10.1021/ja983494x.

[30] T. ting Fu, G. Tu, M. Ping, G. xun Zheng, F. yuan Yang, J. yi Yang, Y. Zhang, X. jun Yao, W. wei Xue, F. Zhu, Subtype-selective mechanisms of negative allosteric modulators binding to group I metabotropic glutamate receptors, Acta Pharmacol. Sin. 42 (2021) 1354–1367. 10.1038/s41401-020-00541-z.

[31] W. Xue, P. Wang, G. Tu, F. Yang, G. Zheng, X. Li, X. Li, Y. Chen, X. Yao, F. Zhu, Computational identification of the binding mechanism of a triple reuptake inhibitor amitifadine for the treatment of major depressive disorder, Physical Chemistry Chemical Physics 20 (2018) 6606–6616. 10.1039/c7cp07869b.

[32] E. Krieger, T. Darden, S.B. Nabuurs, A. Finkelstein, G. Vriend, Making optimal use of empirical energy functions: Force-field parameterization in crystal space, Proteins: Structure, Function and Genetics 57 (2004) 678–683. 10.1002/prot.20251.

[33] T.A. Halgren, Merck molecular force field. I. Basis, form, scope, parameterization, and performance of MMFF94, J. Comput. Chem. 17 (1996) 490–519.

[34] M. Dutta, A.M. Tareq, A. Rakib, S. Mahmud, S.A. Sami, J. Mallick, M.N. Islam, M. Majumder, M.Z. Uddin, A. Alsubaie, A.S.A. Almalki, M.U. Khandaker, D.A. Bradley, M.S. Rana, T. Bin Emran, Phytochemicals from leucas zeylanica targeting main protease of sars-cov-2: Chemical profiles, molecular docking, and molecular dynamics simulations, Biology (Basel). 10 (2021). 10.3390/biology10080789.

[35] G. Tu, T. Fu, F. Yang, J. Yang, Z. Zhang, X. Yao, W. Xue, F. Zhu, Understanding the Polypharmacological Profiles of Triple Reuptake Inhibitors by Molecular Simulation, ACS Chem. Neurosci. 12 (2021) 2013–2026. 10.1021/acschemneuro.1c00127.

[36] E. Krieger, G. Vriend, New ways to boost molecular dynamics simulations, J. Comput. Chem. 36 (2015) 996–1007. 10.1002/jcc.23899.

[37] T. ting Fu, G. Tu, M. Ping, G. xun Zheng, F. yuan Yang, J. yi Yang, Y. Zhang, X. jun Yao, W. wei Xue, F. Zhu, Subtype-selective mechanisms of negative allosteric modulators binding to group I metabotropic glutamate receptors, Acta Pharmacol. Sin. 42 (2021) 1354–1367. 10.1038/s41401-020-00541-z.

[38] S. Mahmud, M.R. Parves, Y.M. Riza, K.M. Sujon, S. Ray, F.A. Tithi, Z.F. Zaoti, S. Alam, N. Absar, Exploring the potent inhibitors and binding modes of phospholipase A2 through in silico investigation, J. Biomol. Struct. Dyn. 38 (2020) 4221–4231.

[39] M.A. Khan, S. Mahmud, A.S.M.R.U. Alam, M.E. Rahman, F. Ahmed, M. Rahmatullah, Comparative molecular investigation of the potential inhibitors against SARS-CoV-2 main protease: a molecular docking study, J. Biomol. Struct. Dyn. (2020) 1–7. 10.1080/07391102.2020.1796813.

[40] M.J. Islam, M.R. Parves, S. Mahmud, F.A. Tithi, M.A. Reza, Assessment of structurally and functionally high-risk nsSNPs impacts on human bone morphogenetic protein receptor type IA (BMPR1A) by computational approach, Comput. Biol. Chem. 80 (2019) 31–45. 10.1016/j.compbiolchem.2019.03.004.

[41] S. Rampogu, R.G. Gajula, G. Lee, M.O. Kim, K.W. Lee, Unravelling the therapeutic potential of marine drugs as SARS-CoV-2 inhibitors: An insight from essential dynamics and free energy landscape, Comput. Biol. Med. 135 (2021) 104525.

[42] G.G. Maisuradze, A. Liwo, H.A. Scheraga, Relation between free energy landscapes of proteins and dynamics, J. Chem. Theory Comput. 6 (2010) 583–595.

[43] E. Papaleo, P. Mereghetti, P. Fantucci, R. Grandori, L. De Gioia, Free-energy landscape, principal component analysis, and structural clustering to identify representative conformations from molecular dynamics simulations: The myoglobin case, J. Mol. Graph. Model. 27 (2009) 889–899. 10.1016/j.jmgm.2009.01.006.

[44] D.S. Jairajpuri, A. Hussain, K. Nasreen, T. Mohammad, F. Anjum, M.T. Rehman, G.M. Hasan, M.F. Alajmi, M.I. Hassan, Identification of natural compounds as potent inhibitors of SARS-CoV-2 main protease using combined docking and molecular dynamics simulations, Saudi J. Biol. Sci. 28 (2021) 2423–2431.

[45] X. Zhao, D. Allison, B. Condon, F. Zhang, T. Gheyi, A. Zhang, S. Ashok, M. Russell, I. MacEwan, Y. Qian, J.A. Jamison, J.G. Luz, The 2.5 Å crystal structure of the SIRT1 catalytic domain bound to nicotinamide adenine dinucleotide (NAD +) and an indole (EX527 analogue) reveals a novel mechanism of histone deacetylase inhibition, J. Med. Chem. 56 (2013). 10.1021/jm301431y.

[46] M. Alrouji, F.A. Alhumaydhi, A. Alsayari, S.E. Sharaf, S. Shafi, S. Anwar, M. Shahwan, A. Atiya, A. Shamsi, Targeting Sirtuin 1 for therapeutic potential: Drug repurposing approach integrating docking and molecular dynamics simulations, PLoS One 18 (2023). 10.1371/journal.pone.0293185.

[47] V.K. Pulla, M. Alvala, D.S. Sriram, S. Viswanadha, D. Sriram, P. Yogeeswari, Structure-based drug design of small molecule SIRT1 modulators to treat cancer and metabolic disorders, J. Mol. Graph. Model. 52 (2014). 10.1016/j.jmgm.2014.06.005.

[48] A. Sharma, V. Gautam, S. Costantini, A. Paladino, G. Colonna, Interactomic and pharmacological insights on human Sirt-1, Front. Pharmacol. 3 MAR (2012). 10.3389/fphar.2012.00040.

[49] I. Autiero, S. Costantini, G. Colonna, Human Sirt-1: Molecular Modeling and Structure-Function Relationships of an Unordered Protein, PLoS One 4 (2013). 10.1371/journal.pone.0007350.

[50] H. Yang, L. Sun, W. Li, G. Liu, Y. Tang, In Silico Prediction of Chemical Toxicity for Drug Design Using Machine Learning Methods and Structural Alerts, Front. Chem. 6 (2018). 10.3389/fchem.2018.00030.

[51] S.S. Swain, T. Hussain, Role of Bioinformatics in Early Drug Discovery: An Overview and Perspective, in: Computation in Bioinformatics: Multidisciplinary Applications, 2021. 10.1002/9781119654803.ch3.

[52] A. Hossain, S. Hossan, S. Prodhan, N.H. Joy, M.R. Siam, E. Rahman, M.N. Matin, Uralenol, Glycyrol, and Abyssinone II as potent inhibitors of fibroblast growth factor receptor 2 from anti-cancer plants: A deep learning and molecular dynamics approach, PLoS One 21 (2026). 10.1371/journal.pone.0341498.

[53] R. Rani, M.E. Hosen, M.N.H. Joy, N. Haq, M.A. Saleh, S. Biswas, A.K. Dutta, Z. Noor, B. Sikdar, M. Khalekuzzaman, M.F. Hasan, Regulation of Shigella effector kinase OspG by Musa acuminata fruit extracts: An in vitro, in vivo, and in silico study, Results Chem. 22 (2026). 10.1016/j.rechem.2026.103127.

[54] M.D. Segall, C. Barber, Addressing toxicity risk when designing and selecting compounds in early drug discovery, Drug Discov. Today 19 (2014). 10.1016/j.drudis.2014.01.006.

[55] J.K. Williams, B. Wang, A. Sam, C.L. Hoop, D.A. Case, J. Baum, Molecular dynamics analysis of a flexible loop at the binding interface of the SARS-CoV-2 spike protein receptor-binding domain, Proteins: Structure, Function and Bioinformatics 90 (2022). 10.1002/prot.26208.

[56] A.A. El-Banna, R.S. Darwish, D.A. Ghareeb, A.M. Yassin, S.A. Abdulmalek, H.M. Dawood, Metabolic profiling of Lantana camara L. using UPLC-MS/MS and revealing its inflammation-related targets using network pharmacology-based and molecular docking analyses, Sci. Rep. 12 (2022). 10.1038/s41598-022-19137-0.

[57] R.S. Katiyar, P.K. Jha, Molecular simulations in drug delivery: Opportunities and challenges, Wiley Interdiscip. Rev. Comput. Mol. Sci. 8 (2018). 10.1002/wcms.1358.

[58] W.F. Drew Bennett, S. He, C.L. Bilodeau, D. Jones, D. Sun, H. Kim, J.E. Allen, F.C. Lightstone, H.I. Ingólfsson, Predicting small molecule transfer free energies by combining molecular dynamics simulations and deep learning, J. Chem. Inf. Model. 60 (2020). 10.1021/acs.jcim.0c00318.

[59] M.A. Maria-Solano, E. Serrano-Hervás, A. Romero-Rivera, J. Iglesias-Fernández, S. Osuna, Role of conformational dynamics in the evolution of novel enzyme function, Chemical Communications 54 (2018). 10.1039/c8cc02426j.

[60] C. Wang, F. He, K. Sun, K. Guo, S. Lu, T. Wu, X. Gao, M. Fang, Identification and characterization of 7-azaindole derivatives as inhibitors of the SARS-CoV-2 spike-hACE2 protein interaction, Int. J. Biol. Macromol. 244 (2023). 10.1016/j.ijbiomac.2023.125182.

[61] S. Ghahremanian, M.M. Rashidi, K. Raeisi, D. Toghraie, Molecular dynamics simulation approach for discovering potential inhibitors against SARS-CoV-2: A structural review, J. Mol. Liq. 354 (2022). 10.1016/j.molliq.2022.118901.

[62] A. Shrivastava, K. Mathur, R.K. Verma, S.K. Jayadev Magani, D.K. Vyas, A. Singh, Molecular dynamics study of tropical calcific pancreatitis (TCP) associated calcium-sensing receptor single nucleotide variation, Front. Mol. Biosci. 9 (2022). 10.3389/fmolb.2022.982831.

[63] Md.T. Islam, Md. Aktaruzzaman, A. Saif, A.R. Hasan, Md.M.H. Sourov, B. Sikdar, S. Rehman, A. Tabassum, S. Abeed-Ul-Haque, M.H. Sakib, Md. M.A. Muhib, Md.A.A. Setu, F. Tasnim, R. Rayhan, M.M. Abdel-Daim, Md.O. Raihan, Identification of acetylcholinesterase inhibitors from traditional medicinal plants for Alzheimer’s disease using *in silico* and machine learning approaches, RSC Adv. 14 (2024) 34620–34636. 10.1039/D4RA05073H.

[64] E. Durham, B. Dorr, N. Woetzel, R. Staritzbichler, J. Meiler, Solvent accessible surface area approximations for rapid and accurate protein structure prediction, J. Mol. Model. 15 (2009). 10.1007/s00894-009-0454-9.

[65] L. Lin, M.K.E. Hasan, X. Gu, S.U. Khan, A. Hossain, M.O. Faruqe, S. Shahriar, M.N.H. Joy, M.M.H. Sourov, M.I. Khan, L. Zhang, M. Lv, Y. Shi, Human tripartite motif-containing protein 71 NCL-1/HT2A/LIN-41 domain crystal structure and its potential natural inhibitors, Int. J. Biol. Macromol. 309 (2025). 10.1016/j.ijbiomac.2025.142764.

[66] Md. N.H. Joy, M.K.E. Hasan, Md.S. Hossan, Md.M.H. Sourov, S. Shahriar, Md.F. Hasan, A.K. Dutta, Md.E. Haque, Computational structure modeling, functional characterization, and identification of potential inhibitors for the cyclic-di-GMP mediated biofilm forming membrane protein in Vibrio cholerae, In Silico Pharmacol. 14 (2026) 221. 10.1007/s40203-026-00719-z.

