## Supplementary material for "Recombinant SIRT1 expression and SIRT1–METTL3 interaction analysis with computational identification of phytochemical SIRT1 activators and inhibitors": Table S1

**Table S1: List of 41 Indian Medicinal plants with phytochemical compounds**

| **Serial no** | **Name of plants** | **Name of Chemical Compounds** | **PubChem**  **CID** | **References** | |
| --- | --- | --- | --- | --- | --- |
| 1 | Acorus calamus | Zingiberene | 92776 | ( Pooja, 2017 ;  Bisht et al., 2011 ;  Samaneh Rahamooz  Haghighi et al.,  2017 ) | |
|  |  | cyclohexanemethanol | 7507 |  |  |
|  |  | calacorene | 528708 |  |  |
|  |  | valencene | 9855795 |  |  |
|  |  | isocalamendiol | 12302240 |  |  |
|  |  | aristolene | 530421 |  |  |
|  |  | naphthalene | 931 |  |  |
|  |  | eugenol | 3314 |  |  |
|  |  | Limonene | 22311 |  |  |
|  |  | camphene | 6616 |  |  |
|  |  | safrol | 5144 |  |  |
|  |  | 𝛼-Asarone | 636822 |  |  |
|  |  | 𝛽-Asarone | 5281758 |  |  |
|  |  | Caryophyllene | 5281515 |  |  |
|  |  | Methyl isoeugenol | 637776 |  |  |
| 2 | Aegle marmelos | Lupeol | 259846 | ( Pooja, 2017 ;  S. Sarla & Subhash,  2011 ; Bisht et al., 2011 ; Revathi &  Lukmanul, 2019 ) | |
|  |  | acetamide | 178 |  |  |
|  |  | benzoic acid | 243 |  |  |
|  |  | pyranocoumarin | 10606 |  |  |
|  |  | scopoletin | 5280460 |  |  |
|  |  | marmesin | 334704 |  |  |
|  |  | psoralen | 6199 |  |  |
|  |  | skimmianine | 6760 |  |  |
|  |  | eugenol | 3314 |  |  |
| 3 | *Ajuga* *parviflora* | senecionine | 5280906 | ( Ankit et al., 2019 ;  Xia et al., 2017 ) | |
|  |  | integerrimine | 5281733 |  |  |
| 4 | Allium cepa | myricetin | 5281672 | ( Ankit et al., 2019 ;  Zeljana et al., 2017 ) | |
|  |  | delphinidin-3-glucoside | 165558 |  |  |
|  |  | quercetin-3-4-diglucoside | 5320835 |  |  |
|  |  | peonidin-3-glucoside | 443654 |  |  |
|  |  | malvidin-3-glucoside | 443652 |  |  |
|  |  | Cepa-2 | 5281672 |  |  |
|  |  | myricetin | 165558 |  |  |
|  |  | delphinidin-3-glucoside | 5320835 |  |  |
| 5 | Allium wallichii | Allixin | 86374 | ( Ankit et al., 2019 ; Jaya &  Muhammad, 2017 ;  Seied & Afra, 2018 )  ( Ankit et al., 2019 ; Jaya & | |
|  |  | 𝛽-Chlorogenin | 12303065 |  |  |
|  |  | Ajoene | 5386591 |  |  |
|  |  | S-Allylcysteine | 9793905 |  |  |
|  |  | Arginine | 6322 |  |  |
|  |  | Glutamic acid | 33032 |  |  |
|  |  | Allicin | 65036 |  |  |
|  |  | Diallyl trisulfide | 16315 |  |  |
|  |  | Methionine | 6137 |  |  |
| 6 | *Aloe* *vera* | Doxorubicin | 31703 | ( Pooja, 2017 ;  Bisht et al., 2011 ;  Karpagam et al.,  2019 ) | |
|  |  | Barbaloin | 12305761 |  |  |
|  |  | Aloe-emodin | 10207 |  |  |
|  |  | Aloesin | 160190 |  |  |
|  |  | Emodin | 3220 |  |  |
|  |  | Campesterol | 173183 |  |  |
|  |  | 6-Methyl-4-chromanone | 584354 |  |  |
| 7 | Andrographis paniculata | Lupeol | 259846 | ( Pooja, 2017 ;  Bisht et al., 2011 ;  Avni et al., 2008 ;  Rajeshkumar1 et al.,  2015 ) | |
|  |  | Stigmasterol | 5280794 |  |  |
|  |  | Andrographolide | 5318517 |  |  |
|  |  | 14-Deoxyandrographolide | 11624161 |  |  |
| 8 | Asparagus racemosus | Rutin | 5280805 | ( Pooja, 2017 ;  Bisht et al., 2011 ;  Verma et al., 2014 )  ( Pooja, 2017 ; | |
|  |  | Quercetin | 5280343 |  |  |
|  |  | Glutathione | 124886 |  |  |
|  |  | Kaempferol | 5280863 |  |  |
| 9 | Annona atemoya / muricata | Chlorogenic Acid | 1794427 | ( Avni et al., 2008 ;  Saranya et al., 2019 ;  S.M. Bassam et al.,  2020 ; Amudha &  Vanitha, 2017 ;  S.M. Bassam et al.,  2020 ; Joseph et al., 2019 ) | |
|  |  | Ledol | 92812 |  |  |
|  |  | Ferulic Acid | 445858 |  |  |
|  |  | Vanillic Acid | 8468 |  |  |
|  |  | Myricetin | 5281672 |  |  |
|  |  | Cleistopholine | 457732 |  |  |
|  |  | Anonaine | 160597 |  |  |
|  |  | Myrcene | 31253 |  |  |
|  |  | Asimilobine | 160875 |  |  |
|  |  | Lanuginosine | 97622 |  |  |
|  |  | Catechin | 9064 |  |  |
| 10 | *Annona* *squamosa* | Limonene | 22311 | ( Saranya et al.,  2019 ; Manoj et al., 2021 ;  Gajalakshmi et al.,  2011 ) | |
|  |  | Anonaine | 160597 |  |  |
|  |  | Aporphine | 114911 |  |  |
|  |  | Isocorydine | 10143 |  |  |
|  |  | Norcorydine | 179491 |  |  |
|  |  | Glaucine | 16754 |  |  |
|  |  | Carvone | 7439 |  |  |
|  |  | Linalool | 6549 |  |  |
|  |  | Eugenol | 3314 |  |  |
|  |  | Bisabolene | 62346 |  |  |
|  |  | Borneol | 64685 |  |  |
|  |  | Geraniol | 637566 |  |  |
|  |  | Germacrene-D | 91723653 |  |  |
| 11 | Arbutus andrachne/unedo | Myricetin | [5281672](https://pubchem.ncbi.nlm.nih.gov/compound/5281672) | (Saranya et al., 2019; A. Emna et al., 2016; A. Emna et al., 2016; Eman et al., 2016; Maria et al., 2014) | |
|  |  | betulin | [72326](https://pubchem.ncbi.nlm.nih.gov/compound/72326) |  |  |
|  |  | kaempferol | [5280863](https://pubchem.ncbi.nlm.nih.gov/compound/5280863) |  |  |
|  |  | betulinic acid | [64971](https://pubchem.ncbi.nlm.nih.gov/compound/64971) |  |  |
|  |  | quercetin | [5280343](https://pubchem.ncbi.nlm.nih.gov/compound/5280343) |  |  |
|  |  | vanillic acid | [8468](https://pubchem.ncbi.nlm.nih.gov/compound/8468) |  |  |
|  |  | gallic acid | [370](https://pubchem.ncbi.nlm.nih.gov/compound/370) |  |  |
|  |  | gentisic acid | [3469](https://pubchem.ncbi.nlm.nih.gov/compound/3469) |  |  |
|  |  | catechins | [1203](https://pubchem.ncbi.nlm.nih.gov/compound/1203) |  |  |
|  |  | procyanidins | [107876](https://pubchem.ncbi.nlm.nih.gov/compound/107876) |  |  |
|  |  | galloylquinic acid | [129650210](https://pubchem.ncbi.nlm.nih.gov/compound/129650210) |  |  |
|  |  | gallotannin | [16129778](https://pubchem.ncbi.nlm.nih.gov/compound/16129778) |  |  |
|  |  | α-linolenic | [5280934](https://pubchem.ncbi.nlm.nih.gov/compound/5280934) |  |  |
|  |  | linoleic | [5280450](https://pubchem.ncbi.nlm.nih.gov/compound/5280450) |  |  |
|  |  | oleic acid | [445639](https://pubchem.ncbi.nlm.nih.gov/compound/5280590) |  |  |
|  |  | lupeol | [259846](https://pubchem.ncbi.nlm.nih.gov/compound/259846) |  |  |
| 12 | Aristolochia ringens | Aristolactam | [96710](https://pubchem.ncbi.nlm.nih.gov/compound/96710) | (Saranya et al.,  2019; Taiye et al.,  2015; Latha et al.,  2015;  Tian-Shung et al.,  2004) | |
|  |  | aporphine | [114911](https://pubchem.ncbi.nlm.nih.gov/compound/114911) |  |  |
|  |  | isoquinoline | [8405](https://pubchem.ncbi.nlm.nih.gov/compound/8405) |  |  |
|  |  | p-cymen-8-ol | [14529](https://pubchem.ncbi.nlm.nih.gov/compound/14529) |  |  |
|  |  | limonene | [22311](https://pubchem.ncbi.nlm.nih.gov/compound/22311) |  |  |
|  |  | 𝛽-caryophyllene | [5281515](https://pubchem.ncbi.nlm.nih.gov/compound/5281515) |  |  |
|  |  | isocaryophyllene | [5281522](https://pubchem.ncbi.nlm.nih.gov/compound/5281522) |  |  |
|  |  | coumarins | [54678486](https://pubchem.ncbi.nlm.nih.gov/compound/54678486) |  |  |
|  |  | tetralone | [10724](https://pubchem.ncbi.nlm.nih.gov/compound/10724) |  |  |
| 13 | Aristolochia bracteolata | Aristolochic acid-I | [2236](https://pubchem.ncbi.nlm.nih.gov/compound/2236) | (Latha et al., 2015;  Tian-Shung et al.,  2004;  Abdelgadir et al.,  2011) | |
|  |  | Aristolochic acids-II | [108168](https://pubchem.ncbi.nlm.nih.gov/compound/108168) |  |  |
|  |  | aristolactam | [96710](https://pubchem.ncbi.nlm.nih.gov/compound/96710) |  |  |
|  |  | aporphine | [114911](https://pubchem.ncbi.nlm.nih.gov/compound/114911) |  |  |
|  |  | protoberberine | [114943](https://pubchem.ncbi.nlm.nih.gov/compound/114943) |  |  |
|  |  | isoquinoline | [8405](https://pubchem.ncbi.nlm.nih.gov/compound/8405) |  |  |
|  |  | coumarins | [54678486](https://pubchem.ncbi.nlm.nih.gov/compound/54678486) |  |  |
|  |  | tetralone | [10724](https://pubchem.ncbi.nlm.nih.gov/compound/10724) |  |  |
| 14 | Artemisia nilagirica | borneol | [64685](https://pubchem.ncbi.nlm.nih.gov/compound/64685) | (Ankit et al., 2019;  Suresh et al., 2011;  Pandey &  Singh, 2017) | |
|  |  | terpinene-4-ol | [11230](https://pubchem.ncbi.nlm.nih.gov/compound/11230) |  |  |
|  |  | 1,8-cineole | [2758](https://pubchem.ncbi.nlm.nih.gov/compound/2758) |  |  |
|  |  | β-caryophyllene | [5281515](https://pubchem.ncbi.nlm.nih.gov/compound/5281515) |  |  |
|  |  | l-linalool | [443158](https://pubchem.ncbi.nlm.nih.gov/compound/443158) |  |  |
|  |  | azulene | [9231](https://pubchem.ncbi.nlm.nih.gov/compound/9231) |  |  |
|  |  | coumarins | [54678486](https://pubchem.ncbi.nlm.nih.gov/compound/54678486) |  |  |
|  |  | acetylene | [6326](https://pubchem.ncbi.nlm.nih.gov/compound/6326) |  |  |
|  |  | 𝛽-eudesmol | [91457](https://pubchem.ncbi.nlm.nih.gov/compound/91457) |  |  |
|  |  | 𝛼-gurjunene | [15560276](https://pubchem.ncbi.nlm.nih.gov/compound/521243) |  |  |
|  |  | para-cymene | [7463](https://pubchem.ncbi.nlm.nih.gov/compound/7463) |  |  |
|  |  | 𝛼-pinene | [6654](https://pubchem.ncbi.nlm.nih.gov/compound/6654) |  |  |
|  |  | stigmasterol | [5280794](https://pubchem.ncbi.nlm.nih.gov/compound/5280794) |  |  |
| 15 | Artemisia herba-alba | Chlorogenic acid | [1794427](https://pubchem.ncbi.nlm.nih.gov/compound/1794427) | (Pandey &  Singh, 2017;  Mohamed et al.,  2010) | |
|  |  | rutin | [5280805](https://pubchem.ncbi.nlm.nih.gov/compound/5280805) |  |  |
|  |  | vicenin-2 | [442664](https://pubchem.ncbi.nlm.nih.gov/compound/442664) |  |  |
|  |  | dihydrocostunolide | [5367639](https://pubchem.ncbi.nlm.nih.gov/compound/5367639) |  |  |
|  |  | piperitone | [6987](https://pubchem.ncbi.nlm.nih.gov/compound/6987) |  |  |
| 16 | Beberis aristata | Berberine | [6987](https://pubchem.ncbi.nlm.nih.gov/compound/6987) | | (Saranya et al.,  2019; S. Sarla &  Subhash, 2011;  Deepti et al., 2014;  Sharma et al., 2011;  Ibrahim et al., 2016) |
|  |  | 𝛽-sitosterol | [2353](https://pubchem.ncbi.nlm.nih.gov/compound/2353) | |  |
|  |  | palmatine | [222284](https://pubchem.ncbi.nlm.nih.gov/compound/222284) | |  |
|  |  | aromoline | [19009](https://pubchem.ncbi.nlm.nih.gov/compound/19009) | |  |
|  |  | berbamine | [362574](https://pubchem.ncbi.nlm.nih.gov/compound/362574) | |  |
|  |  | quercetin | [275182](https://pubchem.ncbi.nlm.nih.gov/compound/275182) | |  |
|  |  | columbamine | [5280343](https://pubchem.ncbi.nlm.nih.gov/compound/5280343) | |  |
|  |  | caffeic acid | [72310](https://pubchem.ncbi.nlm.nih.gov/compound/72310) | |  |
|  |  | chlorogenic acid | [689043](https://pubchem.ncbi.nlm.nih.gov/compound/689043) | |  |
|  |  | rutin | [1794427](https://pubchem.ncbi.nlm.nih.gov/compound/1794427) | |  |
| 17 | Bergenia ciliata | ellagitannin | [10033935](https://pubchem.ncbi.nlm.nih.gov/compound/10033935) | | (Ankit et al., 2019;  Mohammad &  Vimal, 2016;  Ruby et al., 2012;  Vinesh &  Devendra, 2013;  Farman et al., 2016;  Roheena et al., 2019;  Bhupendra et al.,  2020) |
|  |  | gallic acid | [370](https://pubchem.ncbi.nlm.nih.gov/compound/370) | |  |
|  |  | benzaldehyde | [240](https://pubchem.ncbi.nlm.nih.gov/compound/240) | |  |
|  |  | benzeneacetaldehyde | [998](https://pubchem.ncbi.nlm.nih.gov/compound/998) | |  |
|  |  | decadienal | [16899](https://pubchem.ncbi.nlm.nih.gov/compound/16899) | |  |
|  |  | catechin-7-O-glucoside | [10789789](https://pubchem.ncbi.nlm.nih.gov/compound/10789789) | |  |
|  |  | p-hydroxybenzoic acid | [135](https://pubchem.ncbi.nlm.nih.gov/compound/135) | |  |
|  |  | afzelechin | [442154](https://pubchem.ncbi.nlm.nih.gov/compound/442154) | |  |
|  |  | catechin | [9064](https://pubchem.ncbi.nlm.nih.gov/compound/9064) | |  |
|  |  | arbutin | [440936](https://pubchem.ncbi.nlm.nih.gov/compound/440936) | |  |
|  |  | bergenin | [66065](https://pubchem.ncbi.nlm.nih.gov/compound/66065) | |  |
|  |  | protocatechuic acid | [72](https://pubchem.ncbi.nlm.nih.gov/compound/72) | |  |
| 18 | Betula utilis | 𝛽-sitosterol | [222284](https://pubchem.ncbi.nlm.nih.gov/compound/222284) | | (Pooja, 2017;  Bisht et al., 2011;  Tripti et al., 2016) |
|  |  | lupeol | [259846](https://pubchem.ncbi.nlm.nih.gov/compound/259846) | |  |
|  |  | betulinic acid | [64971](https://pubchem.ncbi.nlm.nih.gov/compound/64971) | |  |
|  |  | oleanolic acid | [10494](https://pubchem.ncbi.nlm.nih.gov/compound/10494) | |  |
|  |  | myristic | [11005](https://pubchem.ncbi.nlm.nih.gov/compound/11005) | |  |
|  |  | linalool | [6549](https://pubchem.ncbi.nlm.nih.gov/compound/6549) | |  |
|  |  | palmitic | [985](https://pubchem.ncbi.nlm.nih.gov/compound/5988) | |  |
|  |  | lupenone | [92158](https://pubchem.ncbi.nlm.nih.gov/compound/92158) | |  |
|  |  | oleic | [445639](https://pubchem.ncbi.nlm.nih.gov/compound/445639) | |  |
|  |  | linoleic | [5280450](https://pubchem.ncbi.nlm.nih.gov/compound/5280450) | |  |
|  |  | geranic acid | [5275520](https://pubchem.ncbi.nlm.nih.gov/compound/5275520) | |  |
|  |  | methyl betulonate | [10766700](https://pubchem.ncbi.nlm.nih.gov/compound/10766700) | |  |
|  |  | 𝛽-amyrin | [73145](https://pubchem.ncbi.nlm.nih.gov/compound/73145) | |  |
|  |  | betulin | [72326](https://pubchem.ncbi.nlm.nih.gov/compound/72326) | |  |
|  |  | ursolic acid | [64945](https://pubchem.ncbi.nlm.nih.gov/compound/64945) | |  |
|  |  | karachic acid | [102117094](https://pubchem.ncbi.nlm.nih.gov/compound/102117094) | |  |
|  |  | 1,8-cineol | [2758](https://pubchem.ncbi.nlm.nih.gov/compound/2758) | |  |
|  |  | champacol | [227829](https://pubchem.ncbi.nlm.nih.gov/compound/227829) | |  |
| 19 | Bidens bipinnata | phenylpropanoid | [3314](https://pubchem.ncbi.nlm.nih.gov/compound/3314) | | (Bisht et al., 2011;  Parimalakrishnan et al.,  2006) |
|  |  | glycosides | [637579](https://pubchem.ncbi.nlm.nih.gov/compound/637579) | |  |
|  |  | sesquiterpenes | [667450](https://pubchem.ncbi.nlm.nih.gov/compound/667450) | |  |
|  |  | acetylacetone | [31261](https://pubchem.ncbi.nlm.nih.gov/compound/31261) | |  |
| 20 | Bidens pilosa | Phenylheptatriyne | 77981 | | (Parimalakrishnan et al.,2006) |
|  |  | Linolenic acid | 5280934 | |  |
|  |  | Friedelin | 91472 | |  |
| 21 | Boswellia serrata | Boswellic acid | 168928 | | (Avni et al., 2008;Sudhanshu et al.,2020; Aman &Balu, 2009;Mahe et al., 2012;Nand et al., 2019) |
|  |  | arabinose | 439195 | |  |
|  |  | galactose | 6036 | |  |
|  |  | xylose | 135191 | |  |
|  |  | alpha-pinene | 6654 | |  |
|  |  | borneol | 64685 | |  |
|  |  | myrcene | 31253 | |  |
|  |  | cadinene | 3032853 | |  |
|  |  | verbenone | 29025 | |  |
|  |  | limonene | 22311 | |  |
|  |  | verbenol | 61126 | |  |
|  |  | p-cymene | 7463 | |  |
|  |  | thujene | 18818 | |  |
|  |  | copaene | 76319594 | |  |
|  |  | terpinyl acetate | 111037 | |  |
|  |  | methyl chavicol | 8815 | |  |
|  |  | linalool | 6549 | |  |
|  |  | terpineol | 17100 | |  |
| 22 | Centella asiatica | sistosterol | 222284 | | (Bisht et al., 2011; Avni et al., 2008; Iwan et al., 2016) |
|  |  | asiaticoside | 11954171 | |  |
|  |  | oxyasiaticoside | 18129691 | |  |
| 23 | Cassia fistula | Rhein | 10168 | | (Bisht et al., 2011; Duraipandiyan et al.,2012) |
|  |  | fistucacidin | 51412023 | |  |
|  |  | sitosterol | 222284 | |  |
|  |  | hexacosanol | 68171 | |  |
|  |  | lupeol | 259846 | |  |
|  |  | caprylic | 379 | |  |
|  |  | myristic acids | 481185477 | |  |
|  |  | lecithinphospholipids | 5287971 | |  |
|  |  | epiafzelechin | 443639 | |  |
|  |  | proanthocyanidins | 108065 | |  |
|  |  | epicatechin | 72276 | |  |
| 24 | Catharanthus roseus | Vinorelbine | 5311497 | | (Pooja, 2017;Bisht et al., 2011;Harshini et al.,2020) |
|  |  | vinblastine | 13342 | |  |
|  |  | vincristine | 5978 | |  |
|  |  | vindesine | 40839 | |  |
| 25 | Centella asiatica | Madecassoside | 45356919 | | (Bisht et al., 2011; Iwan et al., 2016) |
|  |  | asiatic acid | 119034 | |  |
|  |  | hydrocotylin |  | |  |
|  |  | brahmoside | 395762 | |  |
|  |  | madecassic acid | 73412 | |  |
|  |  | asiaticosides | 22344906 | |  |
|  |  | centelloside | 274010308 | |  |
|  |  | sitosterol | 222284 | |  |
|  |  | brahminoside | 395762 | |  |
| 26 | Cedrus deodara | Dihydromyricetin | 161557 | | (Avni et al., 2008;Sumeet et al., 2011;Dwaipayan, 2019;Chandur et al., 2011;Amit et al., 2018) |
|  |  | oleoresin | 11980947 | |  |
|  |  | dibenzylbutyrolactol | 18191624 | |  |
|  |  | himachalene | 11586487 | |  |
|  |  | sitosterol | 222284 | |  |
|  |  | ethyl stearate | 8122 | |  |
|  |  | isocentdarol |  | |  |
|  |  | 3-beta‑hydroxy-oleanolic acid | 10494 | |  |
|  |  | shikimic acid | 8742 | |  |
|  |  | ferulic acid | 445858 | |  |
| 27 | Cleome viscosa | 3-glucuronide | 5484731 | | (Bisht et al., 2011;Anuj et al., 2018;Ravi, 2015;Subhash et al., 2021) |
|  |  | lupeol | 259846 | |  |
|  |  | arginine | 6322 | |  |
|  |  | aspartic acid | 5960 | |  |
|  |  | ursolic acid | 64945 | |  |
|  |  | chrysosplenetin | 5281608 | |  |
|  |  | glutamic acid | 33032 | |  |
|  |  | anthroquinones |  | |  |
| 28 | Curcuma longa | Curcumin | 969516 | | (Bisht et al., 2011;Ankit et al., 2019;Avni et al., 2008;Antonio &Giuseppina, 2019) |
|  |  | glucuronide | 5281877 | |  |
|  |  | demethoxycurcumin | 5469424 | |  |
|  |  | bisdemethoxycurcumin | 5315472 | |  |
| 29 | Dioscorea bulbifera | Diosgenin | 99474 | | (Ankit et al., 2019;Gang et al., 2009;Jin-Song et al.,2017; Hilda et al.,2019; S. Sarla &Subhash, 2012) |
|  |  | kaempferol | 5280863 | |  |
|  |  | lutein | 5281243 | |  |
|  |  | zeaxanthin | 5280899 | |  |
|  |  | tristin | 15736297 | |  |
|  |  | protocatechuic acid | 72 | |  |
|  |  | adenosine | 60961 | |  |
|  |  | stigmasterol | 5280794 | |  |
|  |  | azelaic acid | 2266 | |  |
|  |  | caryatin | 5489501 | |  |
|  |  | catechin | 9064 | |  |
| 30 | Punica granatum | Punicic acid | 5281126 | | (Ankit et al., 2019;  Sharrif &  Hamed, 2012; Sheng  & Li, 2017;  Arshad et al., 2017) |
|  |  | myricetin | 5281672 | |  |
|  |  | kaempferol | 5280863 | |  |
|  |  | luteolin | 5280445 | |  |
|  |  | brevifolin | 66654 | |  |
|  |  | isolariciresinol | 160521 | |  |
|  |  | 𝛼-conidendrin | 457194 | |  |
|  |  | pelletierine | 92987 | |  |
|  |  | ursolic acid | 64945 | |  |
|  |  | phloretin | 4788 | |  |
|  |  | coumestrol | 5281707 | |  |
|  |  | asiatic acid | 119034 | |  |
|  |  | isopelletierine | 92987 | |  |
|  |  | matairesinol | 119205 | |  |
|  |  | hovetrichoside-C | 42607791 | |  |
|  |  | betulinic acid | 64971 | |  |
|  |  | pseudopelletierine | 11096 | |  |
|  |  | methionine | 6137 | |  |
|  |  | vanillic acid | 8468 | |  |
|  |  | corilagin | 73568 | |  |
|  |  | chlorogenic acid | 1794427 | |  |
|  |  | neochlorogenic acid | 5280633 | |  |
|  |  | hippomanin-A | 323958 | |  |
|  |  | sinapic acid | 637775 | |  |
|  |  | castalagin | 168165 | |  |
|  |  | isocorilagin | 10077799 | |  |
|  |  | maslinic acid | 73659 | |  |
|  |  | oenothein-B | 16129800 | |  |
| 31 | Rubia Cordifolia  /manjith | Alizarin | 6293 | | (Bisht et al., 2011;  Patel et al., 2011) |
|  |  | rubiadin | 124062 | |  |
|  |  | purpurin | 6683 | |  |
|  |  | munjistin | 160476 | |  |
| 32 | Rumex nepalensis | Chrysophanol | 10208 | | (Ankit et al., 2019;  Yilma et al., 2021;  Samrin et al., 2018;  Nusrat et al., 2017) |
|  |  | physcion | 10639 | |  |
|  |  | emodin | 3220 | |  |
|  |  | aloesin | 160190 | |  |
|  |  | catechin | 9064 | |  |
|  |  | resveratrol | 445154 | |  |
|  |  | orcinol glucoside | 5280450 | |  |
|  |  | linoleic acid | 5280450 | |  |
|  |  | stearic acids | 5281 | |  |
|  |  | isovanillin | 6915 | |  |
|  |  | pentadecanoic acid | 10457 | |  |
|  |  | 1-octadecene | 8223 | |  |
|  |  | ferulic acid | 445858 | |  |
|  |  | palmitic acid | 985 | |  |
|  |  | methylorsellinate | 76658 | |  |
|  |  | neopodin | 23166 | |  |
| 33 | Saussurea costus | Naringenin | 439246 | | (Ankit et al., 2019;  Mohamed et al.,  2021) |
|  |  | kaempferol | 5280863 | |  |
|  |  | ferulic acid | 445858 | |  |
|  |  | malic acid | 525 | |  |
|  |  | ellagic acid | 5281855 | |  |
|  |  | gallic acid | 370 | |  |
|  |  | cinnamic acid | 444539 | |  |
|  |  | chlorogenic acid | 1794427 | |  |
|  |  | vanillin | 1183 | |  |
|  |  | taxifolin | 439533 | |  |
|  |  | catechin | 9064 | |  |
|  |  | syringic acid | 10742 | |  |
|  |  | methyl gallate | 7428 | |  |
|  |  | rutin | 5280805 | |  |
| 34 | Saussurea lappa | Costunolide | 5281437 | | (Ravinder et al.,  2017;  Mohammad et al.,  2013) |
|  |  | linalool | 6549 | |  |
|  |  | α-pinene | 2723720 | |  |
|  |  | myrcene | 31253 | |  |
|  |  | γ-terpinene |  | |  |
|  |  | isodihydrocostunolide | 24893995 | |  |
|  |  | sabinene | 18818 | |  |
|  |  | anethole | 637563 | |  |
|  |  | limonene | 22311 | |  |
|  |  | thymol | 6989 | |  |
|  |  | estragole | 8815 | |  |
|  |  | camphene | 6616 | |  |
|  |  | p-cymene | 7463 | |  |
|  |  | α-terpinolene | 11463 | |  |
|  |  | menthone | 26447 | |  |
|  |  | terpinen-4-ol | 11230 | |  |
|  |  | cryptone | 92780 | |  |
|  |  | ocimene | 5281553 | |  |
|  |  | isozaluzanin | 470970 | |  |
|  |  | zaluzanin-C | 72646 | |  |
|  |  | cynaropicrin | 119093 | |  |
|  |  | lappalone | 10979139 | |  |
|  |  | reynosin | 482788 | |  |
|  |  | isocostic acid | 10922464 | |  |
|  |  | daucosterol | 5742590 | |  |
|  |  | pregnenolone | 8955 | |  |
| 35 | Taxus baccata | Paclitaxel | 36314 | | (Bisht et al., 2011;  Ankit et al., 2019;  Milena et al., 2015) |
|  |  | taxodione | 73588 | |  |
|  |  | gibberellin A-12 | 5459783 | |  |
|  |  | carnosol | 442009 | |  |
|  |  | sugiol | 94162 | |  |
|  |  | ferruginol | 442027 | |  |
|  |  | phenylbutyl | 54678486 | |  |
|  |  | isolariciresinol | 160521 | |  |
|  |  | taxiresinol | 10088963 | |  |
|  |  | lariciresinol | 332427 | |  |
|  |  | baccatin III | 65366 | |  |
| 36 | Terminalia arjuna | Casuarinin | 442673 | | (Bisht et al., 2011) |
|  |  | arjunetin | 21152828 | |  |
|  |  | luteolin | 5280445 | |  |
|  |  | arjunone | 14034821 | |  |
|  |  | 𝛽-sitosterol | 222284 | |  |
|  |  | gallic acid | 370 | |  |
|  |  | ellagic acid | 5281855 | |  |
| 37 | Tinospora  cordifolia | Palmatine | 19009 | | (Pooja, 2017;  Bisht et al., 2011;  Ankit et al., 2019;  Avni et al., 2008;  Rumana &  Srivastava, 2015) |
|  |  | columbin | 442015 | |  |
|  |  | 𝛽-sitosterol | 222284 | |  |
|  |  | isocolumbin | 24721165 | |  |
|  |  | tetrahydropalmatine | 5417 | |  |
|  |  | magnoflorine | 73337 | |  |
|  |  | berberine | 2353 | |  |
|  |  | tinosporic acid |  | |  |
| 38 | Trigonella  foenum-graecum | Diosgenin | 99474 | | (Bisht et al., 2011;  Faris et al., 2021) |
|  |  | cyclophosphamide | 2907 | |  |
|  |  | gitogenin | 441887 | |  |
|  |  | Diosgenin | 99474 | |  |
|  |  | cyclophosphamide | 2907 | |  |
|  |  | gitogenin | 441887 | |  |
| 39 | Urtica dioica | Kaempferol | 5280863 | | (Bisht et al., 2011;  Avni et al., 2008;  Saranya et al., 2019) |
|  |  | isorhamnetin | 5281654 | |  |
|  |  | secoisolariciresinol | 65373 | |  |
|  |  | gentisic acid | 3469 | |  |
|  |  | amentoflavone | 5281600 | |  |
|  |  | ferulic acid | 445858 | |  |
|  |  | quercitrin | 5280459 | |  |
|  |  | catechin | 9064 | |  |
|  |  | esculetin | 5281416 | |  |
|  |  | protocatechuic acid | 72 | |  |
|  |  | scopoletin | 5280460 | |  |
|  |  | chrysoeriol | 5280666 | |  |
|  |  | acetylcholine | 187 | |  |
|  |  | moroidin | 23247762 | |  |
|  |  | carvacrol | 10364 | |  |
|  |  | carvone | 7439 | |  |
|  |  | E-anethol | 637563 | |  |
|  |  | E-𝛽-ionone | 638014 | |  |
|  |  | phytol | 5280435 | |  |
| 40 | Withania  somnifera | Withanolides | 329485 | | (Bisht et al., 2011;  Avni et al., 2008;  Saranya et al., 2019) |
|  |  | anahygrine | 12306778 | |  |
|  |  | anaferine | 443143 | |  |
|  |  | 𝛽-sitosterol |  | |  |
|  |  | chlorogenic acid | 1794427 | |  |
|  |  | somniferine | 14106343 | |  |
|  |  | scopoletin | 5280460 | |  |
| 41 | Ziziphus  nummularia | Betulin | 72326 | | (Avni et al., 2008;  Joelle et al., 2021;  Sayed et al., 2017;  Sonia &  Sumitra, 2019) |
|  |  | betulinic acid | 64971 | |  |
|  |  | campesterol | 173183 | |  |
|  |  | stearic acid | 5281 | |  |
|  |  | linoleic acid | 5280450 | |  |
|  |  | palmitic acid | 985 | |  |
|  |  | squalene | 638072 | |  |
|  |  | lupeol | 259846 | |  |
|  |  | stigmasterol | 5280794 | |  |
|  |  | benzoic acid | 243 | |  |
|  |  | Gamma-sitosterol | 457801 | |  |
|  |  | oleic acid | 445639 | |  |
|  |  | squalene | 638072 | |  |
|  |  | tetratetracontane | 23494 | |  |
|  |  | tricosane | 12534 | |  |
|  |  | tetradecane | 12389 | |  |
|  |  | lapachol | 3884 | |  |
|  |  | 2‑methoxy-4-vinyl phenol | 332 | |  |
|  |  | ethyl alpha-d-glucopyranoside | 9815668 | |  |
